# *In silico* nanoscope to study the interplay of genome organization and transcription regulation

**DOI:** 10.1101/2024.10.22.619557

**Authors:** Soundhararajan Gopi, Giovanni B Brandani, Cheng Tan, Jaewoon Jung, Chenyang Gu, Azuki Mizutani, Hiroshi Ochiai, Yuji Sugita, Shoji Takada

## Abstract

In eukaryotic genomes, regulated access and communication between cis-regulatory elements (CREs) are necessary for enhancer-mediated transcription of genes. The molecular framework of the chromatin organization underlying such communication remains poorly understood. To better understand it, we develop a multiscale modeling pipeline to build near-atomistic models of the 200 kb *Nanog* gene locus in mouse embryonic stem cells comprising nucleosomes, transcription factors, co-activators, and RNA polymerase II-Mediator complexes. By integrating diverse experimental data, including protein localization, genomic interaction frequencies, cryo-electron microscopy, and single-molecule fluorescence studies, our model offers novel insights into chromatin organization and its role in enhancer-promoter communication. The models equilibrated by high-performance molecular dynamics simulations span a scale of ∼350 nm, revealing an experimentally consistent local and global organization of chromatin and transcriptional machinery. Our models elucidate that the sequence-regulated chromatin accessibility facilitates the recruitment of transcription regulatory proteins exclusively at CREs, guided by the contrasting nucleosome organization compared to other regions. By constructing an experimentally consistent near-atomic model of chromatin in the cellular environment, our approach provides a robust framework for future studies on nuclear compartmentalization, chromatin organization, and transcription regulation.

## INTRODUCTION

The genome of eukaryotic organisms is strategically organized and compartmentalized inside the nucleus to provide regulated access to cis-regulatory elements (CREs) for controlled transcription necessary for cell survival, replication, differentiation, and maturation^1–4^. Chromatin organization inferred by Hi-C experiments and its variants report a modular genome organization comprised of coarse chromatin compartments (A/B compartments) with distinct levels of transcriptional activity (high/low, respectively) and a finer organization called topologically associated domains (TADs)^5–8^. TADs are insulated from other genomic regions and are characterized by cis-interacting DNA segments often demonstrated to regulate the expression of encompassed genes^4,9^. The CREs are enriched in transcription factor (TF) binding and active epigenetic signatures, suggesting a strong interplay between genome organization and transcription regulation underlying the cell fate decisions^10–12^.

Despite extensive studies, the mechanistic details of the interplay between genome organization and transcription regulation remain elusive, with conflicting molecular picture in the literature^9,10,12,13^. These discrepancies have been discussed in the context of a range of transcription regulation models, such as contact^8^, network^14^, or diffusion^15,16^-driven transcription regulation differing in the extent of contribution from three-dimensional (3D) genome organization. To address the ambiguity, live-cell time-resolved nanoscopy experiments measure the transcriptional output together with the relative distance of distal genomic locations and molecular factors from the gene locus^10,13,17^. Such advanced in-situ imaging approaches reveal a hierarchical organization of the molecular factors at the transcription site, offering insights into the mechanistic details of transcription activation^18^.

Despite the extent of experimental information available on chromatin organization around transcription start sites - protein localization from ChIP-seq and chemical mapping studies^19–21^, cryoEM structures of ternary complexes of the transcriptional machinery^22,23^, hierarchical organization of protein factors^18^, and Micro-C informed interaction between CREs^8^ – a molecular-level understanding of the chromatin organization and the mechanistic basis of such organization is still lacking.

The transcription regulation of pluripotency factors in mouse embryonic stem cells (mESCs) is well characterized in the literature, supplemented with a plethora of chromatin interaction maps and CRE annotations, representing an ideal system to investigate the interplay between 3D genome organization and transcription at the molecular level. In this work, we develop a multiscale modeling pipeline to build a comprehensive model of mESC gene loci at a near-atomic resolution, integrating several experimental chromatin interaction maps and organization features into a consistent model. Such a model acts as an *in silico* nanoscope and is suitable for subsequent molecular dynamics (MD) simulations to study mechanistic details of the chromatin organization and the intertwined transcription regulation^24^. The molecular modeling of gene loci poses two major challenges – a) localizing protein complexes and factors along a 3-dimensionally organized DNA with characteristic local and global structural features and b) delineating the information from ensemble-averaged chromatin interaction maps (e.g., Micro-C experiments) to generate biologically relevant molecular models. We overcome these challenges by developing a multiscale modeling pipeline that encompasses the ensemble nature of these experiments.

Here, we describe our data-driven modeling pipeline by building comprehensive models of the 200 kb *Nanog* gene locus from mESCs, which contains three enhancers at various distances from the *Nanog* promoter (−45, −5, and +60kb), and is considered a model system for the understanding of enhancer-promoter (E-P) interactions and transcription regulation^25^. Our models integrate nucleosome chemical mapping, ChIP-seq data on protein-DNA binding and epigenetic modifications, and Micro-C genome contact frequencies into an experimentally consistent model of the *Nanog* locus at near-atomic resolution, providing a realistic molecular-level picture of chromatin organization consistent with *in vivo* single-gene imaging studies. The mapping of nucleosomes, linker histones, and transcription (co-)factors, together with the spatial chromatin organization observed from the high-resolution molecular model, reveals the distinct organization principles at CREs. The chromatin is locally expanded at CREs, strategically designed by weak nucleosome positioning signals (NPS) and a longer and asymmetric entry/exit linker DNA that favors local chromatin accessibility for transcription (co)factor binding and recruitment of RNA polymerase II-Mediator complex. The contrasting nucleosome positioning features along the genome guide the mutually excluded localization of linker histones and transcription machinery and suggest the DNA-sequence-guided interplay between chromatin organization and transcription regulation. The expanded chromatin segments at CREs, owing to the increased capture radius, facilitate non-local inter-nucleosome interactions, forming a road map for communication between CREs. The models are an excellent starting point for MD simulations to study the molecular basis of chromatin organization and the role of protein factors, as well as test enhancer-promoter communication models. Overall, the models are expected to be a valuable asset in exploring the structure-function relationships of the genome.

## RESULTS

### CRE interactions at *Nanog* gene locus

Our approach integrates three kinds of experimental data to build an experimentally consistent molecular model of the *Nanog* gene locus: Micro-C contact frequency maps, chemical mapping / ChIP-seq data on protein localization, and *in situ* microscopy on the local concentration of proteins and spatial organization (Figure 1A). The *Nanog* gene locus is particularly interesting as the 200 kb segment (Chr6: 122600-122800 kb) comprises four genes (*Gdf3, Dppa3, Nanog,* and *Slc2a3*), a partial gene at the 5’ end (*Apobec1*), and three super-enhancer (SE) elements, −45SE, −5SE, and +60SE, located at −45, −5, and 60 kb relative to the *Nanog* promoter (Figure 1B), respectively^4,26^. The interaction of the gene promoters with the three SEs is evident from the virtual-4C interaction maps^8^ from the viewpoints of promoters (Figure 1C; note the spike in contact frequency of promoters with other CREs). The contribution of SEs to the transcriptional output of *Nanog* and *Dppa3*^4,11,25^ and the hierarchical organization of TFs at the *Nanog* gene^18^ are well characterized in mESC.

**Figure 1.**
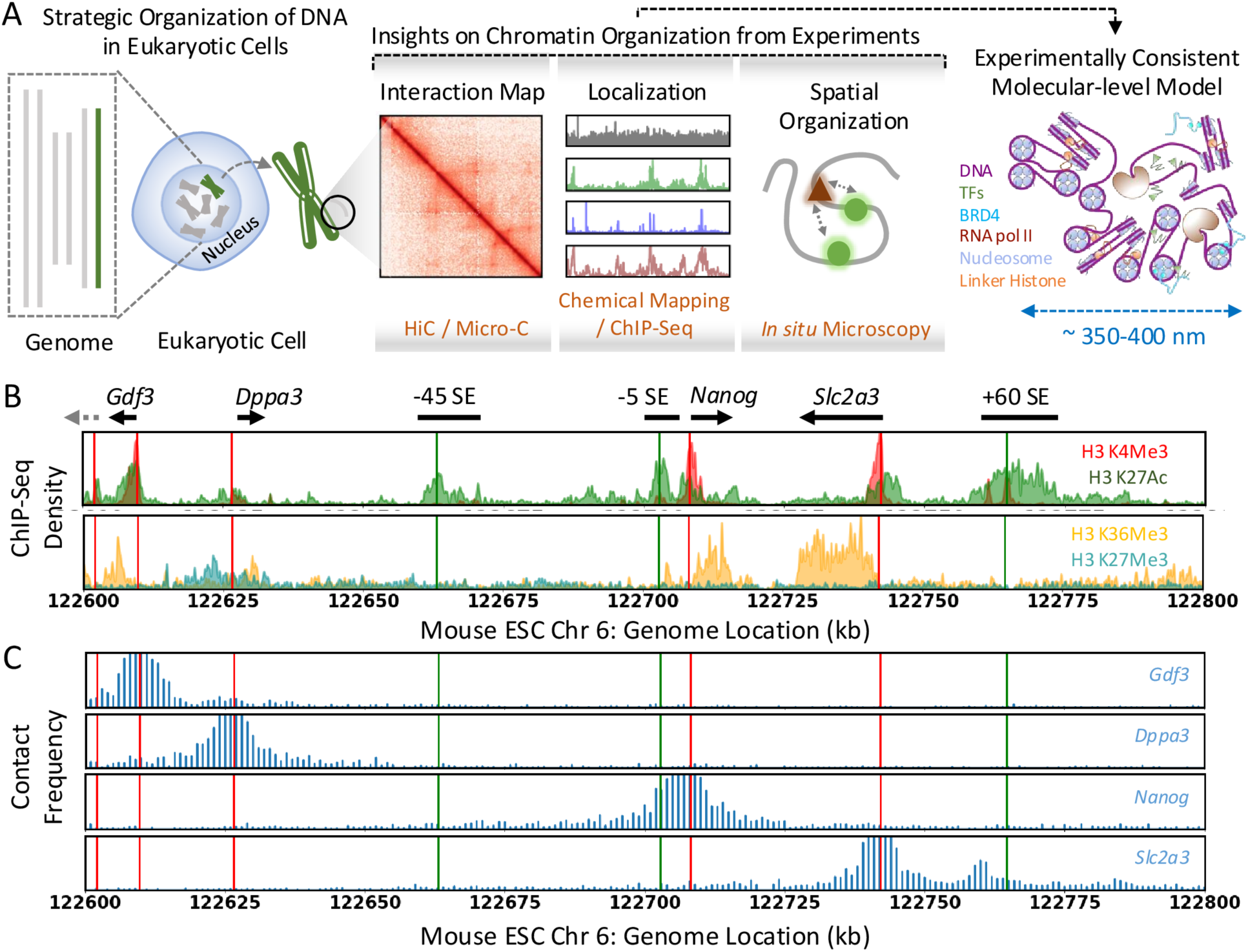
A) Molecular modeling of the mESC *Nanog* gene locus combining a range of experiments reporting on the distinct features of chromatin organization. B) Epigenetic signatures highlight the active enhancers (green), promoters (red), transcriptionally active (orange), and inactive (cyan) genes comprising the *Nanog* locus. The annotations are at the top, and the arrows indicate the orientation of the genes. The dashed grey arrow denotes the location of the partial *Apobec1* gene present inside the modeled chromatin segment. C) Virtual 4C interaction maps reconstructed from the Micro-C data at 1kb resolution from the viewpoint of gene promoters show the selective cis-interactions between promoters and three super-enhancer (SE) elements. As in panel B, the red and green vertical lines denote the location of enhancers and promoters. The y-axis scale is uniform for direct comparison across panels.

### 3D modeling of the *Nanog* Gene Locus

Most chromosome conformation capture methods characterize the 3D genome organization in terms of ensemble-averaged pairwise interaction frequency maps and mask chromatin conformation heterogeneity^27^. The replica-based Bayesian approach Hi-C metainference^28,29^ addresses this by reconstructing chromatin conformational ensembles from experimental contact frequencies and prior models and has been validated against synthetic and *in vivo* data^29^. Using this protocol (Figure 2A; see Methods), we modeled the 200 kb *Nanog* locus with 128 replicas based on mESC Micro-C data and a 1kb resolution chromatin model. The replica-averaged pairwise interaction frequencies match the ensemble-averaged Micro-C data (Pearson’s correlation coefficient *r* = 0.90, Figure 2A-B).

**Figure 2.**
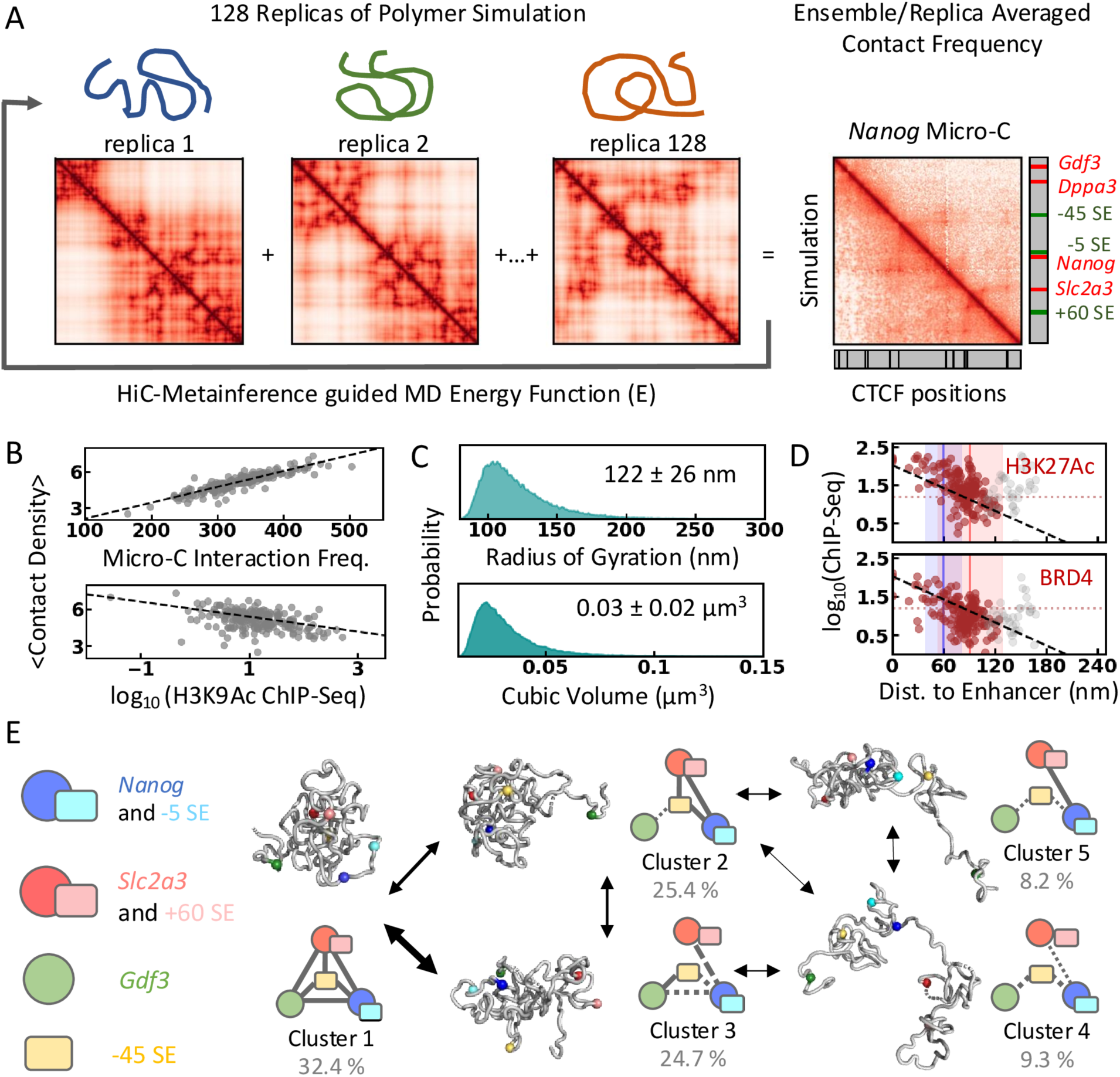
Mesoscopic model of the *Nanog* locus. A) Schematic representation of the Hi-C metainference protocol. 128 replicas of polymer simulations at 1 kb resolution (1 bead = 1 kb) are performed. Hi-C metainference uses Bayesian inference to introduce energy bias such that the replica-averaged pairwise interaction frequency (bottom left panel on the right) quantitatively resembles experimental Micro-C data (top-right panel on the right). B) The average contact density positively correlates with Micro-C contact frequency and shows weak negative dependence on the H3K9 acetylation signal. C) Distribution of radius of gyration and volume of the generated mesoscopic ensemble. D) The log-transformed ChIP-seq signals as the function of ensemble-averaged minimum distance from enhancers (−45, −5, and +60 SE) show a linear dependence. The colored and grey circles correspond to the 160 kb segment at the 3’ end (122640-122800 kb) and the remaining 40 kb, respectively. The blue and red vertical lines mark the average distance of the *Nanog* and *Slc2a3* promoters from the enhancers, respectively, and the shaded area corresponds to 1 standard deviation from the mean. The dotted horizontal line marks the mean ChIP-seq signal. E) Graphical representation of the conformational clusters obtained from k-means clustering and the observed transitions between them. Representative conformations from each cluster used for further modeling are shown in grey, and spherical beads represent the positions of CREs.

We estimate the chromatin ensemble size by rescaling spatial coordinates to the 1kb bead size of 22nm, derived from nucleosome-resolution 1CPN model simulations^29,30^. The average radius of gyration of the *Nanog* locus is 122 ± 26 nm, accounting for ∼0.01% of the nuclear volume (Figure 2C). Assuming a uniform density of ∼5 billion bp in the diploid mouse genome within a 10×10×5 μm ellipsoidal nucleus^31^, the expected *R_g_* is ∼136 nm, slightly larger than our estimate but within the observed range (Figure 2C). We observe a compact organization with a 1/3 power-law scaling of end-to-end distances with segment length^29^ (Figure S1C), characteristic of fractal globule architecture^32,33^, influenced by the bias towards Micro-C data.

The H3K9Ac signal, an epigenetic marker enriched at active enhancers and promoters, shows a linear dependence with average (local and non-local) contact density (*r* = −0.43; Figures 2B and S1A) and relative TF-accessible surface area (a proxy for accessibility of chromatin to TFs; *r* = 0.48; Figure S1B; see Methods). This indicates that the CREs are weakly packed and are preferentially accessible for TFs on the surface of the mesoscopic chromatin ensemble (Figure S1H), which is otherwise compact. This observation reflects the increased DNA accessibility of the acetylated regions probed by the MNase-seq^34^. The interactions between the H3K9Ac enriched beads are similar to non-specific chromatin interactions indicated by their comparable reconfiguration timescales (Figure S1D-E). The CTCF-(and, similarly, SMC1/SMC3-) bound sites show slow reconfiguration, also evident from the slowest independent components from the tICA (time-lagged independent component analysis)^35,36^ analysis of the ensemble (Figures S1D-E and S4D-E), indicative of cohesin’s action as a topological constraint.

ChIP-seq signals of H3K27Ac (active enhancers) and transcriptional cofactor (BRD4) decrease exponentially with distance from enhancers (Figure 2D; *r* = −0.53 for purple circles), suggesting that histone acetylation and cofactor recruitment depend on physical proximity to the three SE elements. Transcription signals extend ∼100 nm from enhancers, with background noise up to ∼200 nm, matching transcriptional condensate sizes observed in microscopy studies^14^. Genomic proximity alone cannot explain this, as the H3K27Ac signal extends up to ∼400 nm (∼18 kb) without spatial context (Figure S1F; *r* = −0.31 for purple circles). This suggests that the spatial proximity of CREs is crucial for enhancer-promoter communication. The 50 kb region upstream of −45 SE does not follow this trend due to the weak dependence of *Apobec1* and *Gdf3* on the three SEs ^4^ (Figure 2A) and may rely on upstream enhancers (Figure S1G).

We applied k-means clustering of the mesoscale ensemble based on the pairwise distance between 6 CREs (*Gdf3, Nanog*, and *Slc2a3* promoters and three SEs; Figures S2 and S3; see Methods) to understand the distinct organization of CREs. The optimal number of clusters is determined using the elbow method, resulting in five representative conformational clusters with distinct organization of CREs (Figures 2E and S4; Table S3). The projection of the mesoscopic chromatin ensemble over the first two tICA components reveals a heterogeneous conformational state within each cluster separated by a fuzzy boundary (Figure S4F-H).

The pairwise correlation map suggests three micro-domains with enhanced correlated motions indicating preferential intra-domain contacts (Figure S4K), each spanning ∼65 kb and bound by cohesin acting as a topological anchor – A) segment containing *Apobec1*, *Gdf3* and *Dppa3*, B) - 45SE to *Nanog* segment and C) *Slc2a3* - +60SE segment, and the relative association of these domains defines the CRE interactions across the clusters (Figures 2E and S4I-L). The contact probabilities (supplementary methods) among the CREs (probed by distinct epigenetic markers on histones) and the cohesin-enriched beads (based on SMC3, SMC1, and CTCF ChIP-seq signals) hint at the distinct topological anchors characterizing the clusters (Figure S5B). We observe a preferential dissociation of acetylated beads compared to non-specific chromatin interaction with identical pairwise sequence separation (Figure S5A). This suggests CRE interactions are not particularly favored but can instead be easily disrupted, possibly by a mechanism similar to the RNA-mediated disruption of transcriptional condensates^37^ (Figure S5A). The representative conformations from each cluster with E-P and E-E distances close to the cluster average (Figures 2E and S6) are used for further modeling.

### Mapping PIC and Nucleosomes

We mapped the transcription pre-initiation complex (PIC), comprising RNA polymerase II, Mediator, and general TFs, based on the overlapping RNA polymerase II and H3K9Ac ChIP-seq peaks (Figure 3A), resulting in 7 PICs at the CREs (see Methods). The number of PICs is close to the lower limit of the RNA polymerase II complexes estimated at the *Nanog* locus^18^. The RNA polymerases II-Mediator complexes are not mapped to the *Nanog* and *Slc2a3* gene termini, as the experimental model of the transcriptional machinery at the termination sites is unavailable.

**Figure 3.**
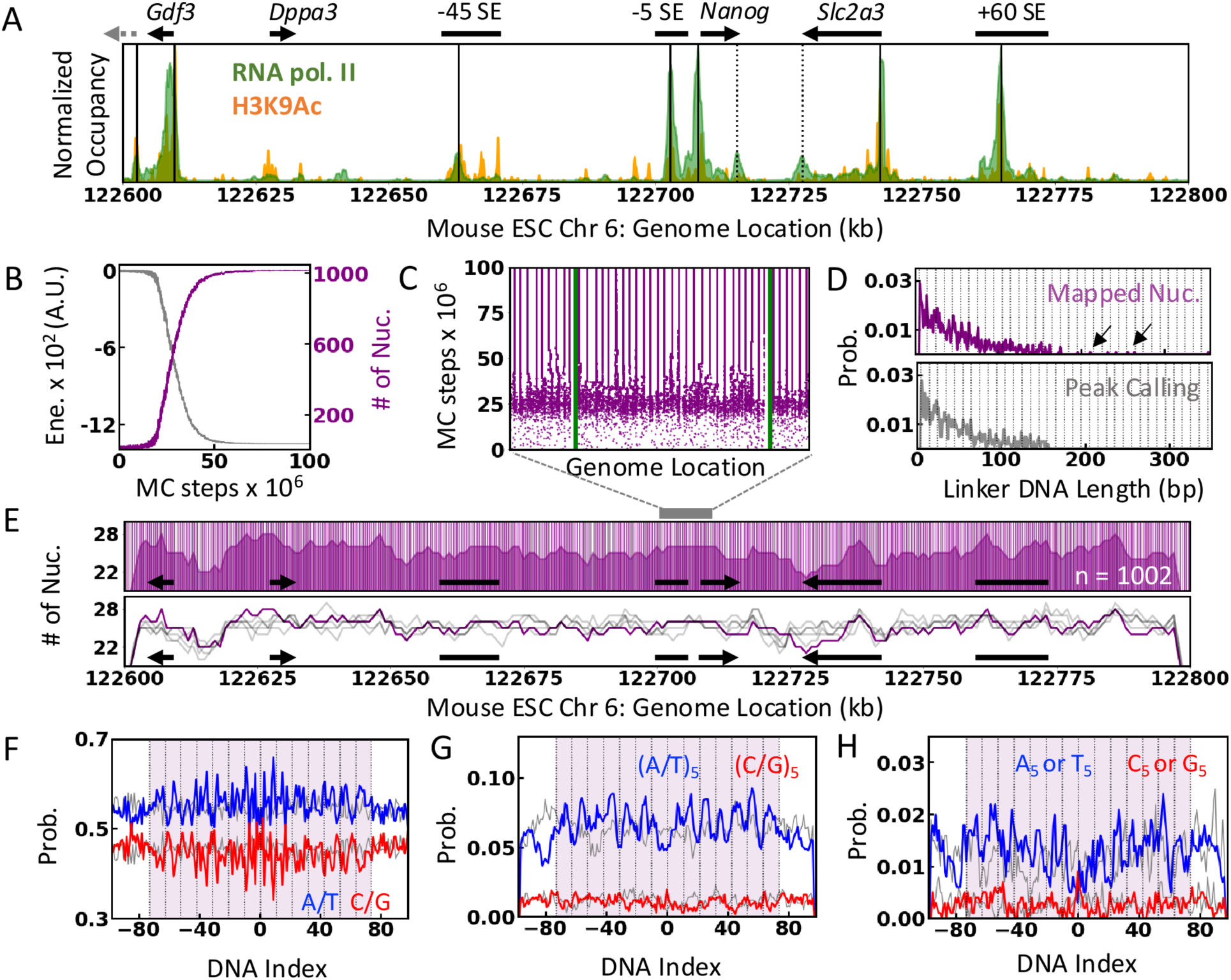
Mapping PIC and nucleosome positions. A) Mapped positions of PIC (solid black line) in the background of RNA pol II (green) and H3K9Ac (orange) ChIP-seq profiles. The CREs and gene annotations are shown at the top for reference. Dashed black lines denote the RNA pol II peaks at the gene ends. B and C) Converging number of nucleosomes, total energy, and nucleosome positions as the function MC steps. D) Distribution of DNA linker length obtained from the representative nucleosome map (top) and the peak-calling method (bottom). The arrows highlight the nucleosome-free regions (>185 bp), and the vertical dotted line indicates the (10n+5) periodicity. E) Nucleosome positions (vertical lines) and the number of nucleosomes per 5kb (shaded area) of the selected nucleosome position map (top). Stochastic variation in the nucleosome positions generated by independent MC simulations (grey) compared to the selected positional map (purple). F-H) Sequence features (X_5_ indicates 5mers of X centered at the position) calculated for the mapped nucleosome positions (colored) and randomly chosen 147bp DNA segments (grey). The dotted vertical lines indicate the position of SHL (10.5 bp) with the dyad position at 0. The shaded area marks the mapped nucleosome position.

The nucleosomes are mapped using Monte Carlo (MC) simulations guided by the nucleosome center positioning scores (NCPS) obtained from experimental chemical mapping^20^ (Figure S7A). The simulated annealing protocol converges to a well-defined positional map with strong NCPS scores compared to the randomly mapped positions (Figures 3B-C and S7B). Consistent with the genome-wide estimate of mESC, the typical nucleosome repeat length (NRL) in the *Nanog* locus ranges between 185-217 bp^38^, and the linker length distribution shows a periodicity of 10n+5^20^ (Figure 3D). Our protocol detects nucleosome-free regions (>185 bp) around the PIC-loaded regions (Figures 3D and S7D) that are not apparent in the conventional peak-calling methods. Individual MC runs generate slightly different nucleosome positional maps with a periodicity of 10n in positional shift (Figure S7C). This provides an ensemble view of nucleosome positions (Figure 3E) that retain a narrow distribution of nucleosome density measured over a 5 kb sliding window, *i.e.*, 25±2 nucleosomes per 5 kb (Figure 3E). Despite the stochastic nature of the mapping, strong nucleosome positioning is observed in the proximity of transcription start sites (TSS) along the direction of the gene in 50 independent runs, although the uniquely mapped ∼14% of the nucleosomes are roughly uniformly distributed in the 200 kb segment with minimal preference towards the genes and regions flanking enhancers (Figure S7E-F).

We selected a tentative nucleosome map (1002 nucleosomes) for further modeling and analysis. The mapped DNA positions reveal strong A/T phasing patterns with peaks corresponding to the DNA minor grooves facing the histone octamer (SHL ±0.5, ±1.5, etc.; Figure 3F), that becomes prominent for (A/T)_5_ 5-mers (Figure 3G; e.g., ATATA, TTTTA, AAAAA, etc.). The phasing (A/T)_5_ patterning is consistent with the analysis of whole-genome studies using chemical mapping^20^ and MNase-based ChIP-seq^39^, and corresponds to the strong nucleosome positioning sequences (NPS)^40^. The (A/T)_5_ phasing is not apparent at the linker DNA and the randomly mapped positions (grey in Figure 3F-H), validating the nucleosome positions (Figure 3F-G). The poly-A and poly-T stretches (A_5_ - AAAAA and T_5_ - TTTTT) are depleted at the dyad but populate further away from the dyad and at the linker DNA (Figure 3H). A_5_ and T_5_ stretches are reported to decrease the stability of DNA wrapping at the ends and increase the accessibility of nucleosome-wrapped DNA^41^. No apparent (G/C)_5_ patterning is observed, which is expected from the nucleosome mapping using MNase-based ChIP-seq data^42,43^, possibly due to the over-digestion of AT-rich DNA segments by MNase, unlike the chemical mapping methods^20,44^.

### Mapping nucleosome-associated protein factors

The linker histone and transcriptional cofactor BRD4 directly bind nucleosomes at dyad and acetylated histone tails, respectively^45,46^. Their positions are mapped using a similar Monte Carlo-based approach with the energy terms proportional to the ChIP-seq signals averaged over the mapped nucleosome positions (Methods). The energy terms are tuned to reproduce the *in vivo* H1/Nuc ratio (0.36-0.46)^19,47^ and the number of BRD4 measured in the proximity of *Nanog* locus^18^ (Methods), resulting in 418 linker histones (H1/Nuc ≈ 0.42) and 20 BRD4 molecules mapped onto the predetermined nucleosome positions (Figure 4A). The H1 density per 5 kb shows a broad distribution, ranging from 3 to 20 (∼11 ± 3) H1 per 5 kb, unlike the near-uniform distribution of nucleosomes (Figure 4B).

**Figure 4.**
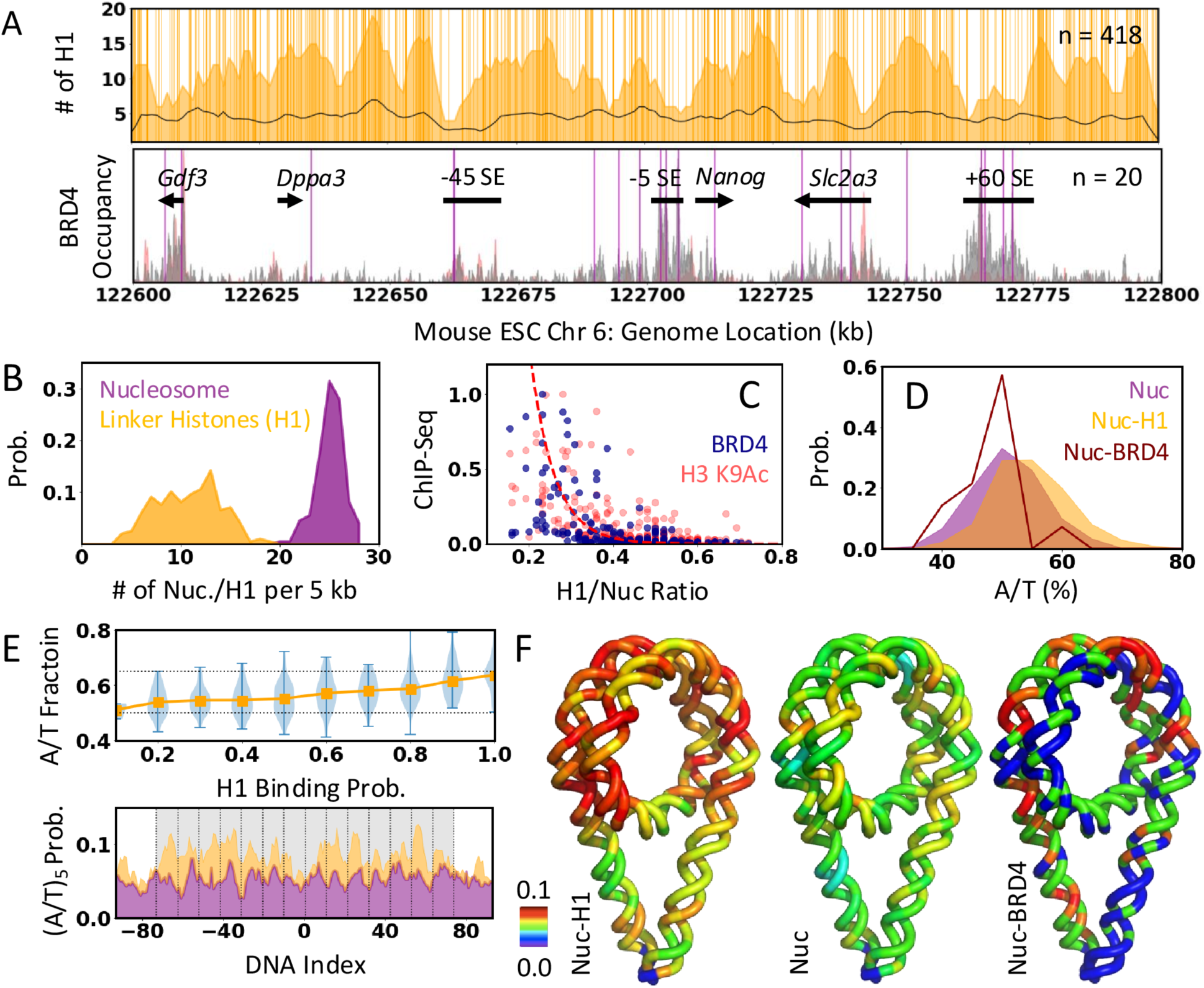
Mapping nucleosome-associated accessory proteins A) Top: Mapped genomic locations of linker histone (vertical lines), the number of linker histones mapped per 5 kb (orange shaded region), and the local contact frequency estimated from Micro-C data (black line). Bottom: Mapped positions of BRD4 (vertical lines) compared against BRD4 and H2K9Ac ChIP-seq profile (grey and red shaded region, respectively). The annotations are shown in black. B) Distribution of the number of nucleosomes (purple) and H1 (orange) per 5kb shown in panel A and Figure 3E. C) Comparison of normalized mean BRD4 and H3K9Ac ChIP-seq frequency and H1/nuc ratio per 5kb sliding window. The red dashed line is an exponential curve to visualize the apparent trend. D) Distribution of A/T% of different nucleosome classes based on the generated positional maps. E) A/T% as the function of H1 binding probability estimated from the experimental data (orange). The distribution of A/T% in each bin is shown in blue. Black dotted lines at 0.5 and 0.7 highlight the linear increase in the A/T%. The bottom panel shows the phasing poly A/T feature of Nuc-H1 (orange) and Nuc (purple). The grey-shaded area marks the position of the nucleosome. F) The poly A/T probability mapped on the nucleosomal DNA for easy visualization.

The H1 density calculated over the 5kb sliding window shows a positive correlation (*r* = 0.52) with the local contact frequency from Micro-C data (orange shaded region and black line in Figure 4A, respectively), indicating compact local organization mediated by the linker histones. The mapped BRD4 positions largely overlap with the BRD4 and H3K9Ac ChIP-seq peaks at the CREs (Figure 4A), consistent with the expectation that BRD4 binds acetylated nucleosomes^48^ and hence, BRD4 bound nucleosomes (Nuc-BRD4) can be used as a proxy for acetylated nucleosomes. Interestingly, the average ChIP-seq frequency of BRD4 (and H3K9Ac), calculated over 5kb sliding windows, exponentially decays with increasing mapped H1 occupancy (Figure 4A and 4C). This suggests that the association of linker histone and BRD4 is mutually exclusive, and the balance between H1 association and acetylation (as BRD4 binds acetylated histones) results in distinct local compaction of chromatin fibers.

The collective mapping of nucleosome, H1 and BRD4 provides a unique opportunity to identify sequence features that may assist the regulated association of the accessory proteins with nucleosome. The three classes of nucleosomes - Nuc-H1 (H1 bound nucleosome), Nuc-BRD4 (BRD4 bound nucleosome or acetylated nucleosome), and Nuc (other free nucleosomes) - show the distinct distribution of A/T fraction (A/T%; Figure 4D), with H1-bound nucleosomes relatively more enriched in A/T% compared to free nucleosomes and Nuc-BRD4 (p-value < 0.001, using two-sample Kolmogorov-Smirnov test). The A/T% of the nucleosomal DNA increases with the increasing H1 binding probability (Figure 4E, top panel), which could be explained by the previously observed higher affinity of linker histones for A/T-rich sequences^49^ (Figure 4E). The phasing of (A/T)_5_ motifs is retained for all classes of nucleosomes (Figure 4E-F) with a strong enrichment of (A/T)_5_ tracks specifically at half-integer SHL for Nuc-H1 and gradually decreases for Nuc followed by Nuc-BRD4.

### Mapping transcription factors and cofactors

We mapped the four pioneer TFs (SOX2, OCT4, NANOG, and KLF4) based on the ChIP-seq density and their cognate sequence bias, assuming a uniform nuclear concentration of 1 *μ*M based on the available experimental estimates^50,51^ (Figure 5A; Methods). Most TFs are mapped at CREs, and the number of each TFs mapped is equivalent to the number of SOX2 estimated at the *Nanog* gene locus *in vivo*^18^. We mapped three P300 molecules based on the theoretical estimate of one P300 per active gene^16^, guided by the ChIP-seq density (Figure 5A).

**Figure 5.**
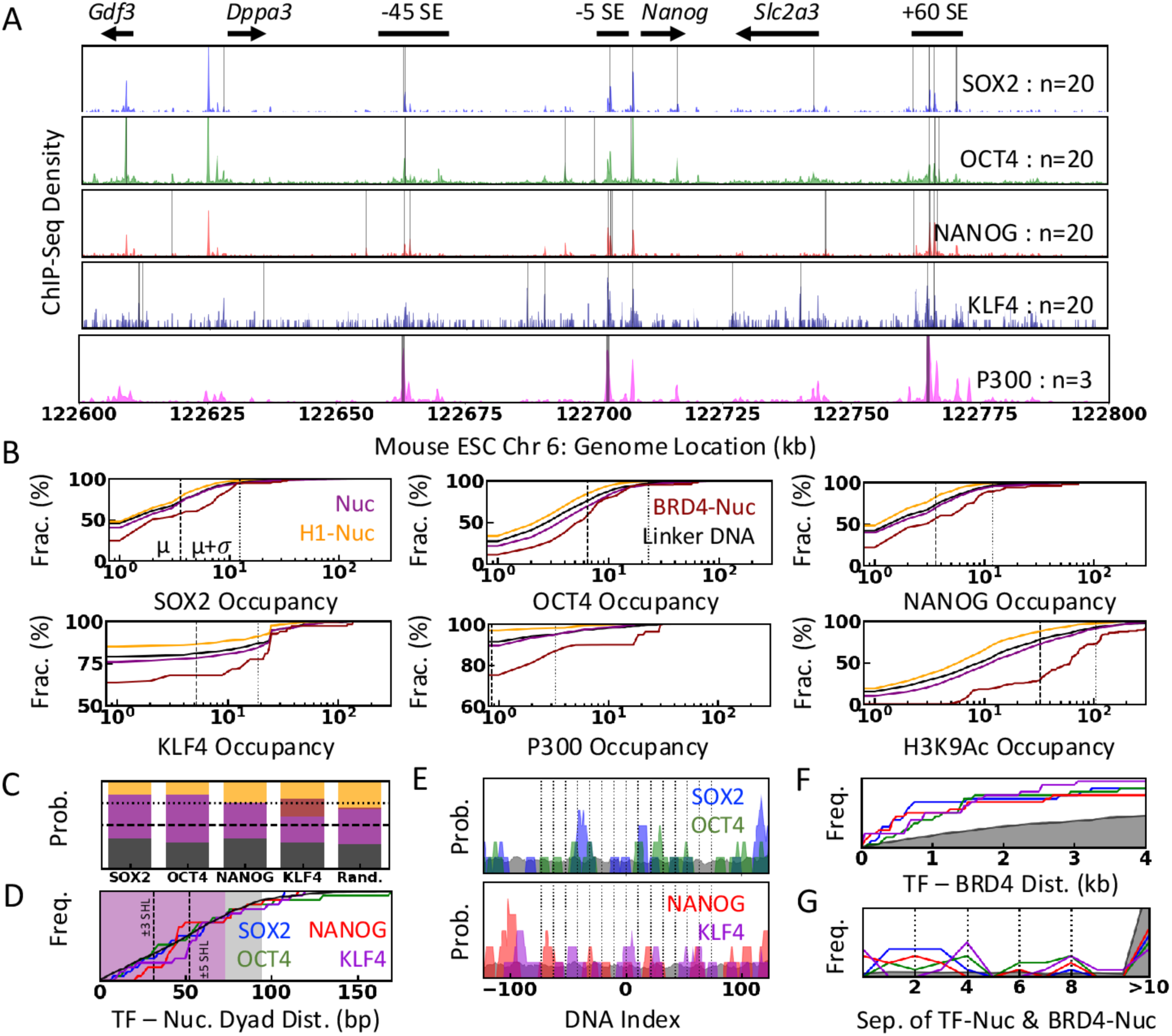
Mapping transcription (co)factors. A) Mapped genomic position of four pioneer transcription factors and P300 (grey vertical bar) alongside the ChIP-seq density (shaded area). The number of molecules mapped is denoted on the right, and the annotations are shown at the top. B) Distribution of ChIP-seq occupancy for each nucleosome class and linker DNA as indicated in the first two panels. The dashed and dotted vertical lines indicate the mean (*μ*) and one standard deviation from the mean (*μ* + *σ*) calculated for each data within *Nanog* locus, respectively. C) Probability of TFs mapped to different nucleosome classes following the color code as in panel B and the distribution expected of a randomly mapped TF with uniform binding preference to the 200kb DNA. The dashed and dotted lines indicate the 0.5 and 0.75 probability, respectively. D) Cumulative distribution of sequence separation between the center of the TF binding site and the dyad of proximal nucleosome. The purple and grey shaded region corresponds to the nucleosome and linker DNA (21 bp corresponding to 189 bp NRL), respectively. The black line indicates the cumulative frequency expected for a randomly mapped TF. E) Probability of mapped TFs as the function of nucleosomal DNA index. The dyad position is indicated as 0, and the dashed black lines denote the nucleosome SHL. The probability expected for a randomly mapped TF is shown as black shaded area. F) Cumulative frequency of the minimum genomic distance between the center of TF-bound DNA and the dyad of BRD4-Nuc following the color code in panel D. The black-shaded region represents the distribution expected for a randomly mapped TF. G) Distribution of shortest nucleosome separation between TF- and BRD4-bound nucleosomes following the same color code in panel F. Dotted black lines indicate the (i-i+2n) nucleosome separation.

The ChIP-seq density of the TFs, cofactors, and acetylation markers are strongly depleted at Nuc-H1 positions (equivalent to the background) while prominently enriched at linker DNA < free nucleosomes < Nuc-BRD4^52^ (Figures 5B and S8A). Based on the strong localization bias of the transcriptional signal, we hypothesize that the sequence-regulated chromatin accessibility, where H1-depleted regions spanning CREs are more accessible for TFs, leads to the cascading cycle of protein recruitment and acetylation required for transcription regulation. In line with these expectations, out of ∼75% of the TFs mapped to the nucleosome, only ∼20% are mapped to H1-Nuc (Figure 5C). The distribution of the TFs over the nucleosome and linker DNA is similar to the randomly mapped TFs except for a subtle decrease in the preference for H1-Nuc (Figure 5C).

Of the four TFs, SOX2, NANOG, and KLF4 bind the minor groove, and OCT4 binds the major groove of the DNA. Specifically, simulations have demonstrated that SOX2 can recognize the exposed target sites better and can induce nucleosome sliding to expose target sites for other TFs^53^. Consistent with this observation, we note that the SOX2 sites populate the exposed minor grooves (SHL −4 to 4; dashed lines on Figures 5D-E and S8D), and the OCT4 sites populate the major grooves facing the histone core. The target sites of the NANOG and KLF4 also show a subtle preference for the exposed minor grooves of the nucleosomal DNA (Figures 5D-E and S8D). About 10% of OCT4 and 35% of NANOG are colocalized with SOX2 on the same nucleosome (Figure S8B). Importantly, ∼50% of each TF is in close genomic proximity (<500 bp or ∼2 nucleosomes away) to all the other TFs, compared to <0.05% for the randomly-mapped TFs, suggesting a strong colocalization of the TFs (Figure S8C).

The combined positional maps reveal that the BRD4 are mapped near TFs (Figure 5F and G). Specifically, ∼50-60% of the mapped TFs are less than 1kb genomic distance from the BRD4-Nuc. Together with nucleosome mapping, ∼50(70)% of the TF-bound nucleosomes are <4(8) nucleosomes away from the BRD4-Nuc with preferential i-i+2n separation. When randomly mapped, TFs are almost always 10 nucleosomes away from the BRD4-Nuc (Figure 5G). We also note that ∼20-50% of the TFs are within 1kb of the three P300 molecules. The spatial proximity of the TFs, P300, and BRD4 likely reflects an underlying molecular mechanism where the TF-recruited acetylation factors (possibly P300) modify the histone tails of proximal nucleosomes^16^.

### Backmapping mesoscopic to near-atomistic resolution

We combine the positional maps generated by the ChIP-seq/Chemical-mapping guided Monte Carlo simulations and the mesoscopic conformations generated by Hi-C-metainference simulations using our backmapping protocol (Figures 6A and S10A-D). This protocol recognizes distinct DNA-protein complexes and generates the chromatin fiber by alternatively connecting the fiber modules – coarse-grained models of nucleosomes (free nucleosome, nucleosome bound to H1 and/or BRD4), PIC, and linker DNA with artificially compacted disordered regions (Figures S9 and S10B) – and minimizes steric clashes, distance to the reference mesoscopic fiber and distance between i-i+2 (D_i-i+2_) nucleosome by a Monte Carlo-like sampling of module conformations at each step (Methods; Figures S10A-D, supplementary videos 1-5). The backmapping protocol accommodates large nucleosome-free regions and simulates an apparent decrease in sedimentation coefficient (S_20,w_) with increasing NRL, qualitatively resembling the experiments^54^ (Figure S10F). Each run produces a structurally different chromatin model with a consistent local arrangement of nucleosomes.

**Figure 6.**
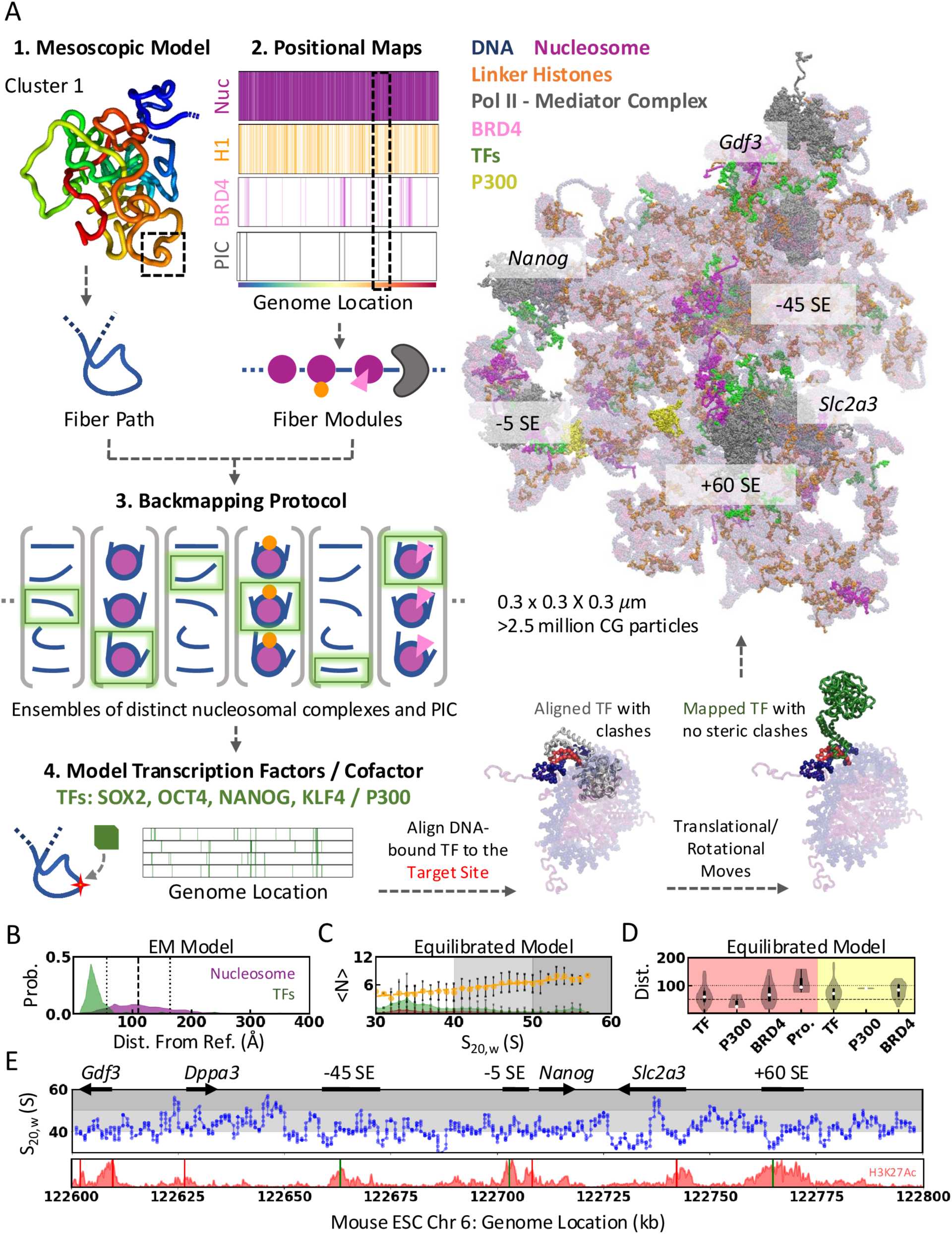
A) Schematic representation of the integrative modeling pipeline. Our semi-automated modeling pipeline combines multiscale modeling strategies to generate in-vivo relevant chromatin models at a nanoscopic resolution consistent with various experiments. B) Distribution of distance of the modeled nucleosomes (purple) and TFs (green) from their reference mesoscopic bead and DNA binding site, respectively, calculated from the energy minimized (EM) model. The radius of the mesoscopic bead (11 nm, black dashed line) ± radius of the nucleosome (5.5 nm, black dotted lines) are shown as references. C) The number of linker histones (orange), TFs (green), and BRD4 (red) mapped as the function of the sedimentation coefficient (S_20,w_) of 12-nucleosome chromatin segments from the equilibrated cluster 1 model. The error bars are calculated as the standard deviation in each bin. The shaded backgrounds indicate the broadly categorized expanded (white), moderately compact (light grey), and compact (dark grey) chromatin segments. D) Distance distribution of the four TFs, P300, BRD4, and promoters to the nearest enhancers (red shaded) and promoters (yellow shaded). E) The S_20,w_ of the 12 nucleosome segments as the function of genomic position. The shaded background is the same as in the panel C. The annotations are given at the top, and the H3K27Ac ChIP-seq signals are shown at the bottom panel for reference.

We generate two complete models for cluster 1 (Model 1 and 1*) to evaluate the reproducibility of the backmapping protocol and one each for the remaining mesoscopic chromatin structure cluster (Figures 2E, 6A, S11, and S12). The RMSD between the two independent models of cluster 1 based on the COM of nucleosomes is 20.3 nm, less than the resolution of the mesoscopic reference structure (22 nm). Across the six models, our backmapping procedure can model ∼45(80)% of the nucleosomes within 11(17) nm from the reference mesoscopic bead (Figures 6B, S10F, and S13A). TFs and P300 are iteratively placed within ∼4±1 and ∼20±10 nm (COM-COM distance; Figures 6B and S13A), respectively, from their target site by algorithmically orienting them to avoid steric clashes, as illustrated in Figure 6A.

The chromatin conformations at near atomistic resolution are energy minimized and equilibrated for 100 ns using GENESIS CGDYN^55^ to remove the modeling artifacts due to the artificially compacted fiber modules used in the backmapping procedure and to evaluate the stability of the generated gene locus models. The backmapping protocol favors compact local structure for convenient modeling, and accordingly, we observe local relaxation and subtle rearrangements favoring multivalent inter-nucleosome interactions after equilibration (RMSD = ∼15.5 nm; Figure S14). The TFs and P300 molecules diffuse freely or by interacting with spatially proximal molecules during the equilibrium process (supplementary methods), resulting in subtle differences in the intermolecular distances measured from the energy-minimized and equilibrated model (Figure S13B-C).

### Chromatin organization at the near-atomistic resolution

The local chromatin organization in our high-resolution models is compatible with the protein localization and mesoscopic ensemble generated using the Micro-C data (Figures 2B, 4A, 6C). The sedimentation coefficient (S_20,w_) calculated for the 12-nucleosome sliding window highlights that the compact chromatin segments have a high H1/Nuc ratio (orange circles in Figure 6C; *r* = 0.95). This aligns with our previous observation that local contact frequencies calculated from the Micro-C data are weakly correlated to the H1 density (Figure 4A) and several experimental evidence of H1-mediated chromatin compaction^54,56–58^. We also observe that chromatin segments mapped with TFs and BRD4 are highly expanded and accessible, as indicated by the low S_20,w_ values (green in Figures 6C, S12, and S15A). Specifically, the chromatin accessibility is higher at CREs (Figures 2B, 6E, S1B, and S16), and we observe an apparent increase in the H3K27Ac ChIP-seq signal with the increased chromatin accessibility observed from our models. The distinct local organization – compact (50-60 S) and expanded (30-40 S) – shows contrasting nucleosome organization and sequence features (Figure S15B-D). The compact chromatin segments have relatively short NRL (175 ± 3 bp) and smaller differences in the entry/exit linker DNA length (∼20 ± 5 bp) and linearly increase to 220 ± 4 bp (*r* = −0.93) and 65 ± 15 bp (*r* = −0.88), respectively, for the expanded segments. We also observe strong nucleosome positioning signals (high A/T%) at compact chromatin segments, corroborating our previous result that H1 favorably binds DNA regions with strong NPS.

The models span ∼350×350×350 nm^3^ and provide a physiologically realistic representation of the *in vivo* chromatin organization and spatial compartmentalization of transcriptional components. In the cluster 1 model, most transcriptional components are encased within a compartment of radius 125 nm, similar to the experimental estimated size of transcriptional condensates (∼100-200 nm)^14^. We observe a hierarchical spatial organization of transcription regulators from the enhancers – TFs (SOX2, OCT4, NANOG, KLF4 at ∼60 nm) < BRD4 (∼66 nm) < promoters (∼93 nm), corroborating with the expected molecular mechanism of transcription (co)factor recruitment, and the spatial molecular organization at the *Nanog* locus^14,16,18^ (Figures 6D and S13B-C). Despite the apparent higher median distance of the TFs from the enhancers compared to P300 (∼24 nm), there are more TFs in the proximity of enhancers compared to P300. We observe a similar hierarchical spatial organization of the TFs and cofactors from the enhancers across all five models, with a subtle increase in median distance with increasing distances between the CREs (Figure S13B-C).

A similar spatial organization is not observed from the viewpoint of promoters, where the distance to the transcription (co)factors is approximately equivalent to or larger than the distance to the enhancers (Figures 6D and S13B-C). Despite the differential spatial organization between promoters and enhancers, the average pairwise distance between the RNA polymerases and SOX2/BRD4 (∼130-220 nm, depending on the cluster) agrees with the estimates from experimental image tracking studies at the *Nanog* locus, further attesting to the quality of global organization represented in our models^18^.

The experimentally consistent local and global molecular organization in our models reveal a network of transcription (co)factors bridging the enhancers and promoters (Figure S17). These models can suggest potential molecular mechanisms underlying transcription regulation, although the lack of large-scale dynamics will make the discussion mostly speculative. A recent multiscale modeling study highlighted the role of nucleosome plasticity in promoting multivalent nucleosome interactions^59^. Considering the distinct local expansion of chromatin at CREs compared to other regions, we analyzed the inter-nucleosome interactions as the function of chromatin compaction. Interestingly, we identify that despite the relatively low inter-nucleosome contacts formed by the expanded 12-nucleosome chromatin segments (∼24 or valency = ∼2; Figure S18A), they favor non-local interactions (Figure S18B) preferably with other expanded segments, compared to the compact segments. The radius of gyration of the expanded 12 nucleosome segments ranges between 20 and 35 nm, facilitating the non-local nucleosomal interactions due to the increased accessibility, and can sufficiently bridge the promoters to the nearest enhancers (Figures 6D, S15A, and S18B-C). This suggests that the local chromatin organization contains the roadmap for communication between CREs and is encoded by the strategically organized nucleosomes.

## Discussion

In this work, we develop a multiscale modeling pipeline to explore the *Nanog* gene locus in mESCs at a near-atomistic resolution. We combine information from experimental ensemble-averaged protein localization, high-resolution pairwise interaction frequencies among genomic loci, cryo-electron microscopy, and *in vivo* single-molecule fluorescent studies using a multiscale approach. The generated model is a first step towards understanding the functional role of chromatin organization in facilitating E-P communication. Thanks to the availability of extensive experimental data, the proposed protocol is easily transferable to model other genomic segments and cell types or the same locus under different developmental or perturbed (e.g., CTCF/cohesin deletions) stages.

In addition to the detailed nanoscopic model, the adopted multiscale modeling methodology also provides insights into the principles of chromatin organization and molecular organization at CREs that are not directly accessible from experimental data of individual chromatin components. We explored how the observed chromatin ensemble informs the interplay between chromatin structure and gene regulation. The graded polyA/T phasing signal for Nuc-H1 > Nuc > Nuc-BRD4 suggests a sequence-regulated chromatin accessibility mechanism (Figure 4F). The strong nucleosome positioning sequences (NPS) may enhance the H1 association via favorable linker DNA geometry resulting from the strong DNA wrapping or decreased nucleosome dissociation and sliding. The resulting compact local nucleosome arrangements inhibit chromatin accessibility for TF/cofactor binding and histone tails for acetylation factors (Figures 4C and 5). Chromatin compaction is further ensured by smaller nucleosome repeat length (NRL) and symmetric entry/exit linker DNA length of the nucleosomes^54^ (Figures 6B-E and S15).

The transcriptional status of the genes is regulated by dynamic modification of the chromatin status at CREs, suggesting regulated access of TFs and cofactors to CREs. Accordingly, CREs are populated with weak NPS that favor the transient local nucleosome-nucleosome interactions due to a relatively high unwrapping rate^59^. The larger NRL and asymmetric entry/exit linker DNA length could disrupt the regular local organization and H1 binding due to increased electrostatic repulsion between the linker DNA and unfavorable linker DNA geometry^54,57,58^. The strategically orchestrated chromatin expansion and possibly increased nucleosome turnover ensure the DNA accessibility for protein binding and histone tail modifications. Together with the acetylation-dependent contact density and TF-ASA observed in the Hi-C metainference simulations (Figures 2B and S1A-B), the results reveal an organization principle at CREs favoring the accessibility of DNA for transcriptional (co)factors and other regulatory proteins. The contrasting chromatin organization at CREs and elsewhere reflects the preferential localization of CREs into kilobase-scale A compartments, observed in the recent *in situ* Hi-C studies^60^. We hypothesize that the recruited TFs and the subsequent histone modifications propagate to the proximal chromatin segments in a distance-dependent manner, as observed in the mesoscopic chromatin ensemble (Figure 2D), resulting in the hierarchical organization of the transcription (co)factors (Figures 6C and S13C).

The population distribution of the clusters from mesoscopic simulations suggests that the cluster with the most CRE interactions, cluster 1, is also the most populated compared to the clusters with weak interactions among CREs (Figure 2E). However, CREs show contact frequencies and distance reconfiguration comparable to the other mesoscopic beads, suggesting that the CRE interactions are comparable to non-specific chromatin interactions, unlike the strong association between CTCF-bound beads acting as topological restraints (Figures S1D-E, S4J, and S5). This could possibly result from using a uniform bead radius for 1 kb chromatin segments in the HiC-metainference simulations, whereas in reality, CREs are likely to be relatively expanded. Based on the molecular model, we also observe that the expanded chromatin segments favor transient non-local inter-nucleosome interactions with other expanded segments (Figures 7 and S18). We propose that such transient non-local inter-nucleosome contacts – “Nucleosomal Handshake” – form the roadmap for cis-regulatory interactions. Overall, the detailed analysis of our 3D models suggests that the design principle for such a roadmap is encoded by the nucleosome organization along the genome, which is summarized in Figure 7.

**Figure 7:**
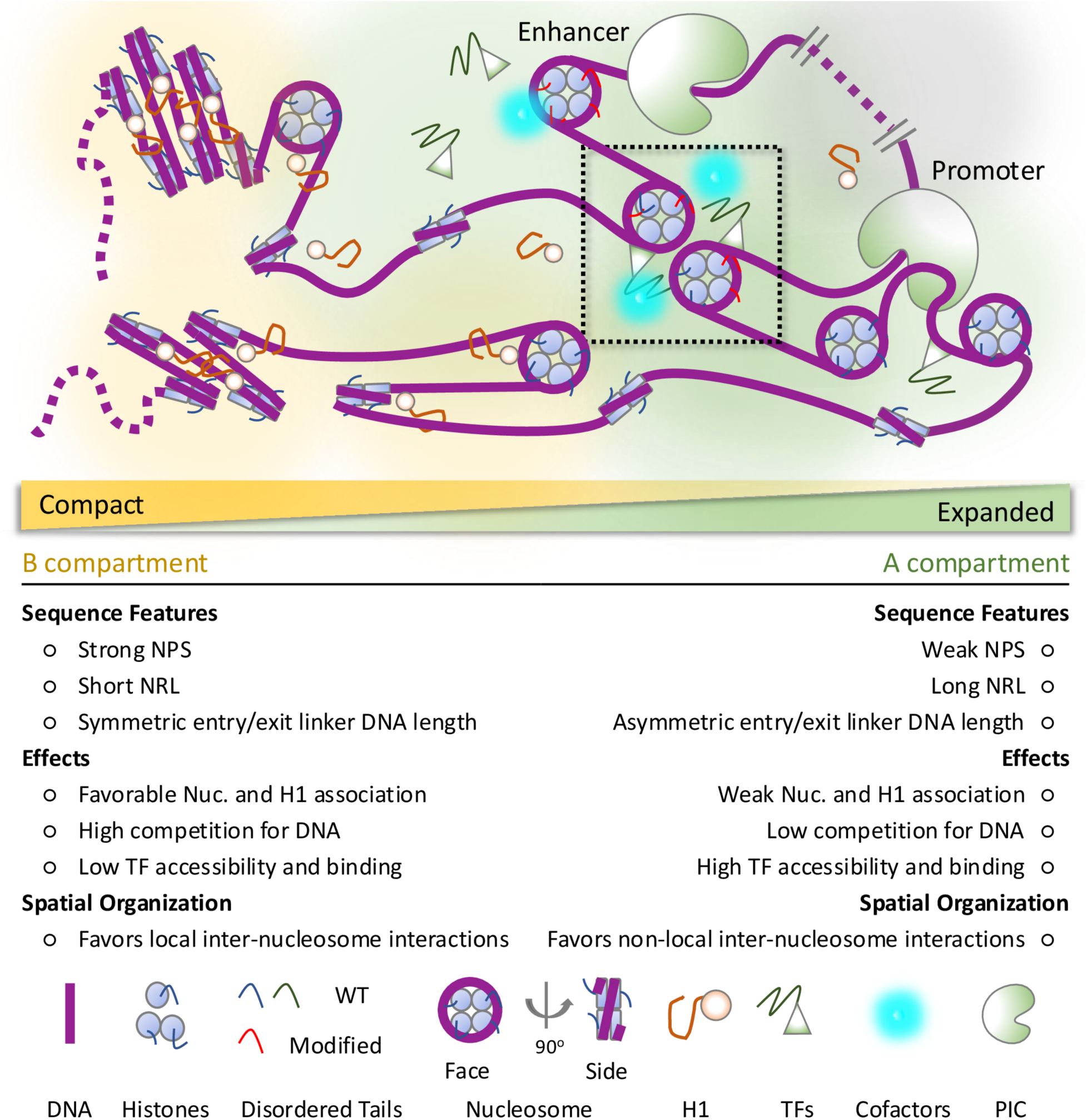
Cartoon representation of the sequence and nucleosome organization principles guiding the distinct local organization of the chromatin. The CREs are locally expanded, and the increased accessibility facilitates the binding of regulatory protein complexes. The transient non-local interactions between these expanded segments (dotted black box) – “Nucleosomal Handshake” – possibly form the roadmap for cis-interactions.

The high-resolution *Nanog* model provides a realistic molecular-level picture of chromatin organization consistent with *in vivo* single-gene imaging studies^18^. The model is also an excellent starting point for MD simulations to study the molecular basis of interplay between chromatin organization and communication between CREs, as well as the mechanistic details of the molecular interactions at CREs^14,18,61,62^. Past chromatin simulations established that the local nucleosome organization could sufficiently drive transient interactions between CREs and extensively studied the roles of NRLs, NDR, H1 association, and acetylation in local chromatin organization^29,63–70^. More recently, nucleosome-resolution integrative modeling and simulations of several 50-100 kb gene loci integrated the Micro-C and nucleosome positioning data to generate *in situ* relevant chromatin fiber models^71^. In our work, we integrate the site-specific protein association data and an ensemble view of chromatin organization to reveal molecular organization principles consistent with *in vivo* experiments. Importantly, our near-atomic large-scale models of the *Nanog* locus bridge the existing understanding of local chromatin organization with that of large-scale 3D genome architecture, revealing the interplay between chromatin structure and transcription regulation. While much work has explored the effect of protein association and histone tail modifications on chromatin^53,63,65,67,69,70,72,73^, our work provides insights into the features driving the orchestrated protein association along the genome.

A current limitation of the molecular models of the *Nanog* locus presented here is that they are simplistic regarding the constituting protein factors and include only the most fundamental components required for chromatin organization and enhancer-mediated transcription regulation. Our current method does not yet include important protein complexes such as cohesin, chromatin remodelers, topoisomerases, several hundred other TFs, the complete epigenetic status of chromatin, and the effect of prevalent histone variants. However, the protocol demonstrated here can accommodate the additional details depending on the purpose of the study and the spatial motions accessible within the timescales of the simulations. Finally, the proposed model is a step toward building an experimentally consistent atomistic molecular model of eukaryotic cells, expanding previous advances to build cellular-scale models of cytoplasm and synaptic vesicles^74–76^.

## METHODS

### Integrative Modelling Pipeline

The genome contact frequencies are used to build an ensemble of 3D chromatin conformations at a coarse resolution (1kb) using Bayesian polymer simulation^29^. Based on chemical mapping and ChIP-seq experimental data, we employ data-driven Monte Carlo simulations to generate an ensemble representation of nucleosome positioning and protein localization consistent with the experiments. The representative protein positional maps and coarse chromatin conformation are combined using a backmapping pipeline to generate the molecular model of the *Nanog* locus at near-atomistic resolution.

The DNA sequence of the *Nanog* gene locus (Chr6: 122600-122800 kb) and its annotations are obtained from the reference mouse genome^26^ (Accession ID: GCA_000001305.2; mm10; release date: 2012/01/09). The occupancy and epigenetic profiles obtained in various formats (supplementary methods) are converted into bedGraph format using the BEDOPS package^77^ and to mm10 genome assembly using the LiftOver tool (http://genome.ucsc.edu).

### Generating a mesoscopic model of the *Nanog* locus

The 3D conformational ensemble of chromatin in the *Nanog* locus was based on mESC Micro-C data^8^ at 1kb resolution, normalized by Juicebox^78^. To generate the conformations, a 128-replica Hi-C metainference MD simulation^28,29^ is performed using a prior 1kb-resolution chromatin model defined by harmonic bond, angle, and Lennard-Jones potentials, with the bead size of 22 nm and other parameters based on higher-resolution chromatin simulations with the 1CPN model^30^. Hi-C metainference performs replica polymer simulations based on the prior model and introduces an additional Bayesian energy score to ensure the agreement between experimental and replica-averaged contact frequencies calculated using a distance-dependent forward model (supplementary methods). The *Nanog* locus is then simulated as a polymer made of 200 1kb beads for 1 million MD steps, and the final half of the replica trajectories are accumulated and further analyzed. The conformational ensemble is clustered based on pairwise distances between the CREs into five conformational clusters with distinct combinations of cis-interactions (supplementary methods). Representative conformations from each cluster with E-P and enhancer-enhancer (E-E) distances close to the cluster average are selected for generating high-resolution molecular models.

For the analysis, a more detailed time-lagged independent component analysis^35^ (tICA) is performed over pairwise genomic contacts at 5kb resolution using the PyEMMA python package^36^. The relative TF accessible surface area (r. TF-ASA) is calculated using an 11 nm bead and 3 nm probe radius and normalized by the surface area of the beads. The distance autocorrelation function is calculated for residue pairs and fitted to the single-exponential functions to measure their decay rate (1⁄*τ_r_*) and hence their reconfiguration time *τ_r_*. The protein occupancy/epigenetic markers are mapped to the mesoscopic beads at 1kb resolution and are considered enriched if the Z-value = (*x* − *μ*)⁄*σ* is >1, where μ and σ the mean and standard deviation calculated over the 200 kb segment, respectively, and treated as background if Z<1.

### Positional Maps of Protein Factors and Complexes

The positions of RNA polymerase II–Mediator complexes (transcription pre-initiation complex; PIC) along the target locus are determined based on the ChIP-seq occupancy profile, and the orientation of the PIC is determined by the relative signal of the positive and negative strands from the GRO-seq experiment^79^. The nucleosomes are mapped on DNA using simulated annealing Monte Carlo simulations with protein association, dissociation, and translocation moves. The energy function is proportional to the nucleosome occupancy measured by *in vivo* chemical mapping experiments^20^, and the energy constant is optimized to achieve saturated nucleosome association. Similarly, linker histones and BRD4 (a transcriptional cofactor) are mapped onto the previously mapped nucleosome positions to reproduce the *in vivo* H1/Nuc—ratio and the number of BRD4 molecules observed by microscopy in the proximity of *Nanog* locus^18,19,47^ (supplementary methods).

Twenty copies of each TF (SOX2, OCT4, NANOG, and KLF4) are added, assuming a uniform 1 *μM* concentration inside the nucleus^50,51^ at positions ranked based on their ChIP-seq signal and the strength of their cognate DNA sequences (supplementary methods). Due to the low copy number of P300 in cells^16^, three copies of P300 are added to the model in the proximity of transcriptional factor clusters at super-enhancer (SE) regions.

### Backmapping mesoscopic model to near-atomistic model

Based on the positional maps of nucleosomes and the associated proteins (*i.e.*, nucleosome – DNA wound around core histones, Nuc-H1 – nucleosome bound to H1, Nuc-BRD4 – H3-acetylated nucleosome bound to BRD4; hereafter called nucleosome modules), the *Nanog* locus is deconstructed into fragments of nucleosome and PIC modules connected by linker DNA of various length (collectively referred to as fiber modules). The fiber modules at atomistic resolution are modeled using MODELLER^80^ if the structural template of homologous protein complexes is available – nucleosome: 1KX5, nucleosome bound to H1: 7K5Y, nucleosome bound to BRD4: 2WP1, and PIC: 7ENC and 6W1S (supplementary methods). The long-disordered regions (LDR) in PIC lacking structural templates are selectively modeled for subunits that are known to participate in liquid-liquid phase separation: MED1, MED14-15, Pol II subunits, TFIID3-5, and TFIID11^81,82^. The structural models for the LDRs are generated using AlphaFold^83^ and are added to the PIC model using MODELLER to account for any residual structures. About 83 out of 203 LDRs in the remaining subunits are trimmed (see supplementary methods) to avoid the risk of topological loops and knots formed while modeling LDRs at the interface of protein subunits. The all-atom structures of the other modules are modeled using MODELLER – SOX2: 1GT0, OCT4: 3L1P, NANOG: 2VI6, KLF4: 4M9E or obtained from the AlphaFold protein structure database^84^ – BRD4: Q9ESU6, and P300: B2RWS6. The molecular models are coarse-grained to residue level, and the disordered tails are artificially compacted using GENESIS to reduce the chances of clashes during backmapping (supplementary methods).

Using representative structures from the 1kb metainference mesoscopic model as a reference, we grow the chromatin fiber of the *Nanog* locus from one end to the other by adding one nucleosome module (+ PIC module, if PIC is at the consecutive position) at a time by generating structural ensembles on the fly (supplementary methods). The chromatin fiber grows by adding the nucleosome modules that vary in nucleosomal DNA unwrapping, orientation, and linker DNA bending to the current fiber (supplementary methods). The fiber modules are joined by aligning the phosphates of the three terminal bp of DNA. At each step, the structure of the growing chromatin fiber is sampled using a Monte Carlo-guided procedure attempting to avoid the clashes between the newly added nucleosome module and the structure generated until the previous step, and at the same time, minimize the distance from the reference 1kb mesoscopic bead and the distance between i-1 and i+1 nucleosomes to promote a compact local nucleosome organization and decrease the chances of topological knots. The protocol also ensures that the growing fiber is knot-free by rejecting knotted structures on the fly at each iteration^85,86^. The longer linker DNA is compacted by introducing decoy nucleosomes to avoid topological loops during the modeling. After chromatin fiber backmapping is complete for the reference mesoscopic conformations, the DNA component of the generated *Nanog* locus molecular models is coarse-grained to 25 bp resolution and analyzed for topological knots using KymoKnot^85^, and the backmapping procedure is repeated if knots are identified.

The performance of the backmapping protocol is evaluated by generating 25 nucleosome chromatin fibers with varying nucleosome repeat lengths (NRL=167-207) and reference polymer structures with five beads. The distance from the reference bead and the distance between i and i+2 nucleosomes are calculated as the distance between the center of mass (COM) of the nucleosome. The sedimentation coefficient of the nucleosome fibers is calculated as before^87^:

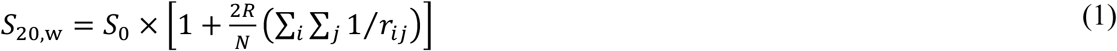

here, *S*_0_ = 11.1 *S* is the sedimentation coefficient of a mononucleosome, *R* = 54.5 Å is the spherical radius of a mononucleosome, *N* is the total number of nucleosomes, and *r_ij_* is the distance between the COM of nucleosomes *i* and *j*.

Finally, the compacted TFs and P300 models are added in the proximity of mapped genomic locations by iteratively sampling the translational- and rotational-moves to decrease steric clashes while retaining the binding orientation and proximity to the mapped genomic location (supplementary methods).

The molecular models of the *Nanog* locus (∼2.5 million CG particles) are energy-minimized and subsequently simulated at 150 mM ionic strength and 300 K using GENESIS CGDYN^55,88^ on the Fugaku supercomputer. The simulations employed the AICG2+ model^89^ for proteins, with a statistical flexible and solvation potential for disordered residues^90,91^, and the 3SPN2.C model^92^ for DNA with a −1.0 charge on phosphate groups (supplementary methods). The simulations are conducted for 100 ns (10^7^ MD steps with 10 fs integration timesteps) to relax the artificially compacted fiber modules used in the backmapping procedure and to demonstrate the stability of the generated molecular model for MD simulations.

## ASSOCIATED CONTENT

All processed input files, Hi-C metainference trajectory, all-atom and coarse-grained models of the fiber modules, 200 kb CG *Nanog* models, GENESIS CGDYN setup files, and modeling scripts used in this study are available upon request. Code to perform Hi-C metainference simulations is maintained on GitHub at https://github.com/gbrandani/Hi-C_Metainference.

## AUTHOR INFORMATION

### Author Contributions

S.T. designed the research; G.B.B and C.G. performed HiC-metainference simulations; S.G. and S.T. performed positional mapping of proteins; S.G., G.B.B., and A.M. generated all-atom molecular models of protein complexes; S.G., and G.B.B., performed backmapping; S.G., G.B.B., C.T., J. J., A.M., performed CG MD simulations; S.G., G.B.B., and S.T. analyzed the data; H. O., Y. S., and S.T. supervised the research; S.G. wrote the manuscript with help from all authors.

### Funding Sources

This work was supported by the MEXT grants JPMXP1020200101 as “Program for Promoting Research on the Supercomputer Fugaku” (S.T., Y.S.) and JPMXP1020230119 as “Program for Promoting Researches on the Supercomputer Fugaku” (S.T.), and by the Japan Society for the Promotion of Science KAKENHI grants 20H05934 (S.T.) and 21H02441 (S.T.). This work used computational resources of the supercomputer Fugaku provided by RIKEN through the HPCI System Research Project (Project ID: hp230095; S.G.).

### Notes

None

## ACKNOWLEDGMENT

We thank Diego Ugarte La Torre, Computational Biophysics Research Team, RIKEN Center for Computational Science, Kobe, Hyogo, Japan, for his help in modeling the mouse pre-initiation complex.

## ABBREVIATIONS

CRE: Cis-Regulatory Elements
TF: Transcription Factor
CG: Coarse Grained
MD: Molecular Dynamics
PIC: transcription Pre-Initiation Complex
tICA: time-lagged Independent Component Analysis
NPS: Nucleosome Positioning Signal
NCPS: Nucleosome Center Positioning Score

## Notes

### Competing Interest Statement

The authors have declared no competing interest.

## REFERENCES

1. Banigan, E. J. et al. Transcription shapes 3D chromatin organization by interacting with loop extrusion. Proceedings of the National Academy of Sciences 120, (2023).

2. Chacin, E. et al. Establishment and function of chromatin organization at replication origins. Nature 616, 836–842 (2023).

3. Dixon, J. R. et al. Chromatin architecture reorganization during stem cell differentiation. Nature 518, 331–336 (2015).

4. Blinka, S., Reimer, M. H., Pulakanti, K. & Rao, S. Super-Enhancers at the Nanog Locus Differentially Regulate Neighboring Pluripotency-Associated Genes. Cell Rep 17, 19–28 (2016).

5. Lieberman-Aiden, E. et al. Comprehensive Mapping of Long-Range Interactions Reveals Folding Principles of the Human Genome. Science (1979) 326, 289–293 (2009).

6. Nora, E. P. et al. Spatial partitioning of the regulatory landscape of the X-inactivation centre. Nature 485, 381–385 (2012).

7. Hsieh, T.-H. S. et al. Mapping Nucleosome Resolution Chromosome Folding in Yeast by Micro-C. Cell 162, 108–119 (2015).

8. Hsieh, T.-H. S. et al. Resolving the 3D Landscape of Transcription-Linked Mammalian Chromatin Folding. Mol Cell 78, 539–553.e8 (2020).

9. Batut, P. J. et al. Genome organization controls transcriptional dynamics during development. Science (1979) 375, 566–570 (2022).

10. Benabdallah, N. S. et al. Decreased Enhancer-Promoter Proximity Accompanying Enhancer Activation. Mol Cell 76, 473–484.e7 (2019).

11. Agrawal, P. & Rao, S. Super-Enhancers and CTCF in Early Embryonic Cell Fate Decisions. Front Cell Dev Biol 9, (2021).

12. Lee, R. et al. CTCF-mediated chromatin looping provides a topological framework for the formation of phase-separated transcriptional condensates. Nucleic Acids Res 50, 207–226 (2022).

13. Alexander, J. M. et al. Live-cell imaging reveals enhancer-dependent Sox2 transcription in the absence of enhancer proximity. Elife 8, (2019).

14. Sabari, B. R. et al. Coactivator condensation at super-enhancers links phase separation and gene control. Science (1979) 361, (2018).

15. Richter, W. F., Nayak, S., Iwasa, J. & Taatjes, D. J. The Mediator complex as a master regulator of transcription by RNA polymerase II. Nat Rev Mol Cell Biol (2022) doi:10.1038/s41580-022-00498-3.

16. Karr, J. P., Ferrie, J. J., Tjian, R. & Darzacq, X. The transcription factor activity gradient (TAG) model: contemplating a contact-independent mechanism for enhancer–promoter communication. Genes Dev 36, 7–16 (2022).

17. Su, J.-H., Zheng, P., Kinrot, S. S., Bintu, B. & Zhuang, X. Genome-Scale Imaging of the 3D Organization and Transcriptional Activity of Chromatin. Cell 182, 1641–1659.e26 (2020).

18. Li, J. et al. Single-Molecule Nanoscopy Elucidates RNA Polymerase II Transcription at Single Genes in Live Cells. Cell 178, 491–506.e28 (2019).

19. Cao, K. et al. High-resolution mapping of h1 linker histone variants in embryonic stem cells. PLoS Genet 9, e1003417 (2013).

20. Voong, L. N. et al. Insights into Nucleosome Organization in Mouse Embryonic Stem Cells through Chemical Mapping. Cell 167, 1555–1570.e15 (2016).

21. Chronis, C. et al. Cooperative Binding of Transcription Factors Orchestrates Reprogramming. Cell 168, 442–459.e20 (2017).

22. El Khattabi, L., et al. A Pliable Mediator Acts as a Functional Rather Than an Architectural Bridge between Promoters and Enhancers. Cell 178, 1145–1158.e20 (2019).

23. Chen, X. et al. Structures of the human Mediator and Mediator-bound preinitiation complex. Science (1979) 372, (2021).

24. Huertas, J., Woods, E. J. & Collepardo-Guevara, R. Multiscale modelling of chromatin organisation: Resolving nucleosomes at near-atomistic resolution inside genes. Curr Opin Cell Biol 75, 102067 (2022).

25. Blinka, S. & Rao, S. *Nanog* Expression in Embryonic Stem Cells - An Ideal Model System to Dissect Enhancer Function. BioEssays 39, 1700086 (2017).

26. Lee, B. T. et al. The UCSC Genome Browser database: 2022 update. Nucleic Acids Res 50, D1115–D1122 (2022).

27. Cheng, R. R. et al. Exploring chromosomal structural heterogeneity across multiple cell lines. Elife 9, (2020).

28. Bonomi, M., Camilloni, C., Cavalli, A. & Vendruscolo, M. Metainference: A Bayesian inference method for heterogeneous systems. Sci Adv 2, (2016).

29. Brandani, G. B., Gu, C., Gopi, S. & Takada, S. Multiscale Bayesian simulations reveal functional chromatin condensation of gene loci. PNAS Nexus 3, (2024).

30. Lequieu, J., Córdoba, A., Moller, J. & de Pablo, J. J. 1CPN: A coarse-grained multi-scale model of chromatin. J Chem Phys 150, 215102 (2019).

31. Zhou, Y., Basu, S., Laue, E. & Seshia, A. A. Single cell studies of mouse embryonic stem cell (mESC) differentiation by electrical impedance measurements in a microfluidic device. Biosens Bioelectron 81, 249–258 (2016).

32. Grosberg, A., Rabin, Y., Havlin, S. & Neer, A. Crumpled Globule Model of the Three-Dimensional Structure of DNA. Europhysics Letters (EPL) 23, 373–378 (1993).

33. Mirny, L. A. The fractal globule as a model of chromatin architecture in the cell. Chromosome Research 19, 37–51 (2011).

34. Schwartz, U. et al. Characterizing the nuclease accessibility of DNA in human cells to map higher order structures of chromatin. Nucleic Acids Res 47, 1239–1254 (2019).

35. Naritomi, Y. & Fuchigami, S. Slow dynamics in protein fluctuations revealed by time-structure based independent component analysis: The case of domain motions. J Chem Phys 134, (2011).

36. Scherer, M. K. et al. PyEMMA 2: A Software Package for Estimation, Validation, and Analysis of Markov Models. J Chem Theory Comput 11, 5525–5542 (2015).

37. Henninger, J. E. et al. RNA-Mediated Feedback Control of Transcriptional Condensates. Cell 184, 207–225.e24 (2021).

38. Fan, Y. et al. H1 Linker Histones Are Essential for Mouse Development and Affect Nucleosome Spacing In Vivo. Mol Cell Biol 23, 4559–4572 (2003).

39. Segal, E. et al. A genomic code for nucleosome positioning. Nature 442, 772–778 (2006).

40. Lowary, P. T. & Widom, J. New DNA sequence rules for high affinity binding to histone octamer and sequence-directed nucleosome positioning. J Mol Biol 276, 19–42 (1998).

41. Anderson, J. D. & Widom, J. Poly(dA-dT) Promoter Elements Increase the Equilibrium Accessibility of Nucleosomal DNA Target Sites. Mol Cell Biol 21, 3830–3839 (2001).

42. Kaplan, N. et al. The DNA-encoded nucleosome organization of a eukaryotic genome. Nature 458, 362–366 (2009).

43. Yoo, J., Park, S., Maffeo, C., Ha, T. & Aksimentiev, A. DNA sequence and methylation prescribe the inside-out conformational dynamics and bending energetics of DNA minicircles. Nucleic Acids Res 49, 11459–11475 (2021).

44. Chereji, R. V., Ramachandran, S., Bryson, T. D. & Henikoff, S. Precise genome-wide mapping of single nucleosomes and linkers in vivo. Genome Biol 19, 19 (2018).

45. Filippakopoulos, P. et al. Histone Recognition and Large-Scale Structural Analysis of the Human Bromodomain Family. Cell 149, 214–231 (2012).

46. Zhou, Y.-B., Gerchman, S. E., Ramakrishnan, V., Travers, A. & Muyldermans, S. Position and orientation of the globular domain of linker histone H5 on the nucleosome. Nature 395, 402–405 (1998).

47. Zhang, Y., Liu, Z., Medrzycki, M., Cao, K. & Fan, Y. Reduction of Hox Gene Expression by Histone H1 Depletion. PLoS One 7, e38829 (2012).

48. Devaiah, B. N. et al. BRD4 is a histone acetyltransferase that evicts nucleosomes from chromatin. Nat Struct Mol Biol 23, 540–548 (2016).

49. Käs, E., Izaurralde, E. & Laemmli, U. K. Specific inhibition of DNA Binding to nuclear scaffolds and histone H1 by distamycin. J Mol Biol 210, 587–599 (1989).

50. Chen, J. et al. Single-Molecule Dynamics of Enhanceosome Assembly in Embryonic Stem Cells. Cell 156, 1274–1285 (2014).

51. Xie, L. et al. A dynamic interplay of enhancer elements regulates *Klf4* expression in naïve pluripotency. Genes Dev 31, 1795–1808 (2017).

52. Peng, Y. et al. Detection of new pioneer transcription factors as cell-type-specific nucleosome binders. Elife 12, (2024).

53. Tan, C. & Takada, S. Nucleosome allostery in pioneer transcription factor binding. Proceedings of the National Academy of Sciences 117, 20586–20596 (2020).

54. Correll, S. J., Schubert, M. H. & Grigoryev, S. A. Short nucleosome repeats impose rotational modulations on chromatin fibre folding. EMBO J 31, 2416–2426 (2012).

55. Jung, J., Tan, C. & Sugita, Y. GENESIS CGDYN: large-scale coarse-grained MD simulation with dynamic load balancing for heterogeneous biomolecular systems. Nat Commun 15, 3370 (2024).

56. Song, F. et al. Cryo-EM Study of the Chromatin Fiber Reveals a Double Helix Twisted by Tetranucleosomal Units. Science (1979) 344, 376–380 (2014).

57. Hou, Z., Nightingale, F., Zhu, Y., MacGregor-Chatwin, C. & Zhang, P. Structure of native chromatin fibres revealed by Cryo-ET in situ. Nat Commun 14, 6324 (2023).

58. Gibson, B. A. et al. Organization of Chromatin by Intrinsic and Regulated Phase Separation. Cell 179, 470–484.e21 (2019).

59. Farr, S. E., Woods, E. J., Joseph, J. A., Garaizar, A. & Collepardo-Guevara, R. Nucleosome plasticity is a critical element of chromatin liquid–liquid phase separation and multivalent nucleosome interactions. Nat Commun 12, 2883 (2021).

60. Harris, H. L. et al. Chromatin alternates between A and B compartments at kilobase scale for subgenic organization. Nat Commun 14, 3303 (2023).

61. Kagey, M. H. et al. Mediator and cohesin connect gene expression and chromatin architecture. Nature 467, 430–5 (2010).

62. Shrinivas, K. et al. Enhancer Features that Drive Formation of Transcriptional Condensates. Mol Cell 75, 549–561.e7 (2019).

63. Chang, L. & Takada, S. Histone acetylation dependent energy landscapes in tri-nucleosome revealed by residue-resolved molecular simulations. Sci Rep 6, (2016).

64. Kenzaki, H. & Takada, S. Linker DNA Length is a Key to Tri-nucleosome Folding. J Mol Biol 433, 166792 (2021).

65. Bascom, G. D. & Schlick, T. Chromatin Fiber Folding Directed by Cooperative Histone Tail Acetylation and Linker Histone Binding. Biophys J 114, 2376–2385 (2018).

66. Perišić, O., Collepardo-Guevara, R. & Schlick, T. Modeling Studies of Chromatin Fiber Structure as a Function of DNA Linker Length. J Mol Biol 403, 777–802 (2010).

67. Bascom, G. D., Myers, C. G. & Schlick, T. Mesoscale modeling reveals formation of an epigenetically driven HOXC gene hub. Proceedings of the National Academy of Sciences 116, 4955–4962 (2019).

68. Sridhar, A. et al. Emergence of chromatin hierarchical loops from protein disorder and nucleosome asymmetry. Proceedings of the National Academy of Sciences 117, 7216–7224 (2020).

69. Portillo-Ledesma, S. et al. Nucleosome Clutches are Regulated by Chromatin Internal Parameters. J Mol Biol 433, 166701 (2021).

70. Forte, G. et al. Transcription modulates chromatin dynamics and locus configuration sampling. Nat Struct Mol Biol 30, 1275–1285 (2023).

71. Li, Z. & Schlick, T. Hi-BDiSCO: folding 3D mesoscale genome structures from Hi-C data using brownian dynamics. Nucleic Acids Res 52, 583–599 (2024).

72. Collepardo-Guevara, R. et al. Chromatin Unfolding by Epigenetic Modifications Explained by Dramatic Impairment of Internucleosome Interactions: A Multiscale Computational Study. J Am Chem Soc 137, 10205–10215 (2015).

73. Watanabe, S., Mishima, Y., Shimizu, M., Suetake, I. & Takada, S. Interactions of HP1 Bound to H3K9me3 Dinucleosome by Molecular Simulations and Biochemical Assays. Biophys J 114, 2336–2351 (2018).

74. McGuffee, S. R. & Elcock, A. H. Diffusion, Crowding & Protein Stability in a Dynamic Molecular Model of the Bacterial Cytoplasm. PLoS Comput Biol 6, e1000694 (2010).

75. Wilhelm, B. G. et al. Composition of isolated synaptic boutons reveals the amounts of vesicle trafficking proteins. Science (1979) 344, 1023–1028 (2014).

76. Feig, M. et al. Complete atomistic model of a bacterial cytoplasm for integrating physics, biochemistry, and systems biology. J Mol Graph Model 58, 1–9 (2015).

77. Neph, S. et al. BEDOPS: high-performance genomic feature operations. Bioinformatics 28, 1919–1920 (2012).

78. Robinson, J. T. et al. Juicebox.js Provides a Cloud-Based Visualization System for Hi-C Data. Cell Syst 6, 256–258.e1 (2018).

79. Williams, L. H. et al. Pausing of RNA Polymerase II Regulates Mammalian Developmental Potential through Control of Signaling Networks. Mol Cell 58, 311–322 (2015).

80. Webb, B. & Sali, A. Comparative Protein Structure Modeling Using MODELLER. Curr Protoc Bioinformatics 54, (2016).

81. Palacio, M. & Taatjes, D. J. Merging Established Mechanisms with New Insights: Condensates, Hubs, and the Regulation of RNA Polymerase II Transcription. J Mol Biol 434, 167216 (2022).

82. Wang, X. et al. LLPSDB v2.0: an updated database of proteins undergoing liquid–liquid phase separation *in vitro*. Bioinformatics 38, 2010–2014 (2022).

83. Jumper, J. et al. Highly accurate protein structure prediction with AlphaFold. Nature 596, 583–589 (2021).

84. Varadi, M. et al. AlphaFold Protein Structure Database: massively expanding the structural coverage of protein-sequence space with high-accuracy models. Nucleic Acids Res 50, D439–D444 (2022).

85. Tubiana, L., Polles, G., Orlandini, E. & Micheletti, C. KymoKnot: A web server and software package to identify and locate knots in trajectories of linear or circular polymers. The European Physical Journal E 41, 72 (2018).

86. Dabrowski-Tumanski, P., Rubach, P., Niemyska, W., Gren, B. A. & Sulkowska, J. I. Topoly: Python package to analyze topology of polymers. Brief Bioinform 22, (2021).

87. Arya, G., Zhang, Q. & Schlick, T. Flexible Histone Tails in a New Mesoscopic Oligonucleosome Model. Biophys J 91, 133–150 (2006).

88. Tan, C. et al. Implementation of residue-level coarse-grained models in GENESIS for large-scale molecular dynamics simulations. PLoS Comput Biol 18, e1009578 (2022).

89. Li, W., Terakawa, T., Wang, W. & Takada, S. Energy landscape and multiroute folding of topologically complex proteins adenylate kinase and 2ouf-knot. Proceedings of the National Academy of Sciences 109, 17789–17794 (2012).

90. Terakawa, T. & Takada, S. Multiscale Ensemble Modeling of Intrinsically Disordered Proteins: p53 N-Terminal Domain. Biophys J 101, 1450–1458 (2011).

91. Tesei, G., Schulze, T. K., Crehuet, R. & Lindorff-Larsen, K. Accurate model of liquid– liquid phase behavior of intrinsically disordered proteins from optimization of single-chain properties. Proceedings of the National Academy of Sciences 118, (2021).

92. Freeman, G. S., Hinckley, D. M., Lequieu, J. P., Whitmer, J. K. & de Pablo, J. J. Coarse-grained modeling of DNA curvature. J Chem Phys 141, 165103 (2014).

