## Supplementary File for "*In silico* nanoscope to study the interplay of genome organization and transcription regulation"

METHODS

1. Mesoscopic model of *Nanog* gene locus

The ensemble of chromatin conformation at mesoscopic resolution is generated using Hi-C metainference approach as previously described^1^. Figure 2A in the main text demonstrates the schematics of the HiC-metainference protocol. In short, replica exchange simulations are performed with a constraint to generate a replica-averaged signal comparable to the experimental ensemble-averaged signal guided by a Bayesian inference method^2^. In our protocol, the experimental signal is the pairwise interaction frequency obtained from the Micro-C experiments^3^, normalized using Juicebox^4^. The mesoscopic model has a resolution of 1kb (200 beads representing the 200kb *Nanog* gene locus). The prior chromatin model is a generalized polymer model with beads linked using harmonic bond and angle potential and interacts with non-neighboring beads via the Weeks-Chandler-Anderson potential optimized to reproduce the 1/2 scaling of the end-to-end distances with segment length. The forward model to calculate the contact frequency is defined as,

$$f_{ij}=\frac{\alpha}{N}\sum_{r} \frac{1-\left[ {(r_{ij}-r_{o})}/{d_{o}} \right]^{6}}{1-\left[ {(r_{ij}-r_{o})}/{d_{o}} \right]^{12}}$$

(1)

where, $r_{ij}$ is the distance between beads *i* and *j*, $r_{o}=0.5d_{o}$ is the radius of the bead set to 11 nm, $\alpha$ is the scaling factor to convert the contact probability to frequency (*i.e.,* Micro-C reads) and *N* is the total number of replicas *r*. The HiC metainference simulations are performed using LAMMPS patched with PLUMED 2, with 128 replicas, to reproduce the experimental Micro-C data^3,5,6^. The simulation data is generated for 10^6^ steps using a single Gaussian error model to sample the scaling factor ($\alpha)$. The conformational ensemble from the final 9x10^5^ steps of each replica is used for further analysis. Code to perform Hi-C metainference simulations with LAMMPS^6^ and PLUMED^5^ is maintained on Github at https://github.com/gbrandani/Hi-C_Metainference.

1.1 Clustering Analysis

The clustering of mesoscopic chromatin conformations is performed using k-means clustering, and the number of clusters is optimized using the elbow method (Figure S4). The conformational ensemble is clustered based on non-redundant enhancer-promoter (E-P) and enhancer-enhancer (E-E) distances after eliminating the weakly correlated E-P/E-E (-5SE-E/P and *Dppa3*-E/P) distances (Figures S2 and S3). We identified five conformational clusters with distinct combinations of E-P and E-E interactions. The conformation with minimum root mean squared difference (RMSD) compared to the average E-P/E-E distances of each cluster is used as the representative conformation for further modeling at near-atomistic resolution (Figure S6). tICA analysis over the contact map, calculated using 1 in 5 mesoscopic beads, is performed using PyEMMA python package^7^ (Figure S3).

2. Positional Maps of Proteins and Protein Complexes

The chromatin model comprises nucleosomes, linker histones, RNA polymerase II-Mediator complexes (or transcription pre-initiation complex; PIC), transcriptional co-factors such as BRD4, P300, and transcriptional factors such as SOX2, OCT4, NANOG, and KLF4. The experimental occupancy profiles are obtained as ChIP-seq or chemical mapping reads (Table S1). The occupancy profiles obtained in various formats are converted into bedgraph format using the BEDOPS package^8^. The mouse genome assembly, GRCm39 (mm10; release date: 2012/01/09), is used as the reference, and the genome coordinates from various assemblies are appropriately converted using the LiftOver tool (<http://genome.ucsc.edu>).

The occupancy profiles of the region of interest – Mouse chromosome 6: 122600 – 122800 kb - are isolated, and the genomic positions of proteins are mapped subsequently.

2.1 Mapping of RNA polymerase-Mediator (PIC) complex: The positions of nine RNA polymerases are selected based on the signal peaks of RNA polymerase II ChIP-seq occupancy and the orientation of the PIC is determined by the relative signal of the positive and negative strands from the GRO-seq experiment^9^. The number of PIC closely matches the experimental estimate of 10-25 RNA polymerase molecules at the *Nanog* Locus^10^. Based on the available experimentally resolved structure of RNA polymerase II in complex with the Mediator subunits (PDB ID: 7enc), we added seven RNA polymerase II-Mediator complexes in the model, excluding the two positions at the gene termination sites.

2.2 Mapping of Nucleosomes: We performed a Monte-Carlo simulation guided by the nucleosome center positioning score (NCPS) obtained using experimental chemical mapping. In the simulation, nucleosomes can associate, dissociate, and slide ($\pm$10 bp) stochastically along the genomic positions. The energy function of any given occupancy state is defined as follows,

$V=V_{single}+V_{pair}+V_{RNAp}$

(2)

$V_{single}$ is the energy of nucleosome association defined as $V_{single}=\sum_{i_{dyad} \in nuc} -\log\left( \boldsymbol{\varepsilon}\times{NCPS}_{i} \right)$, here, *nuc* is the set of nucleosomes, and *ε* is the proportionality constant tuned to obtain saturated levels of the nucleosome (supplementary figure S3).

$V_{pair}$ and $V_{RNAp}$ prevent the overlapping of nucleosomes with other nucleosomes (occupies 151 bp assuming a 2bp linker DNA at each end) and RNA polymerases (occupies 74 bp assuming a 5 bp linker DNA at each end), respectively.

$$V_{pair}= \sum_{i_{dyad}\neq j_{dyad} \in nuc} \left\{ \begin{matrix} \infty abs\left( i-j \right)<151 \\ 0 otherwise \end{matrix} \right.$$

(3)

$V_{RNAp}= \sum_{i_{dyad} \in nuc, j_{bp} \in RNAp} \left\{ \begin{matrix} \infty abs\left( i-j \right)<109 \\ 0 otherwise \end{matrix} \right.$

(4)

2.3 Mapping of Linker Histones: The linker histone depletion scores are converted to binding profiles (1-bound, 0-unbound) using a cutoff (-0.2) equivalent to the background score^11^. The average over experimental replicates is used to calculate the probability of linker histone occupancy, and the linker histone positioning score (LHPS) is calculated as an average probability over the mapped nucleosome positions. The linker histone position map is obtained using the Monte Carlo simulation, similar to the nucleosome mapping procedure, involving association and dissociation moves. The *ε* is tuned to reproduce the experimentally obtained H1/Nuc ratio (0.36 - 0.46)^11,12^.

2.4 Mapping Transcriptional Factors and Cofactors: The *in vivo* nuclear concentrations of SOX2 and OCT4 are estimated to be 0.7-1.3 µM and 2-3 µM, respectively^13,14^. The nuclear concentration of several other pluripotency factors is also in the micromolar range^14^. For simplicity, we assume a uniform 1µM concentration for all the transcription factors and cofactors except for P300^15^. Based on the average volume of the *Nanog* gene locus estimated from the mesoscopic model, this concentration translates to ~20±10 molecules each for SOX2, OCT4, NANOG, KLF4, and BRD4. As measured using single-molecule nanoscopy experiments, the number of SOX2 and BRD4 molecules at the *Nanog* gene locus is 9-17 and 10-25, respectively. The concentration of modeled transcription factors and cofactors is close to the upper limit of the experimental estimates. The nuclear concentration of the P300 is limited, and the number of molecules is equivalent to the number of active genes^13,15^. Hence, three P300 molecules are added near transcription factor clusters at the super-enhancers (-45SE, -5SE, and +60SE), overlapping with P300 ChIP-seq peaks.

BRD4 molecules are added similarly to the linker histones using the Monte-Carlo simulations guided by the corresponding ChIP-seq data. The proportionality constant is tuned to reproduce the number of molecules estimated based on the experiments^10^.

The binding sequence profiles of the transcription factors are obtained from JASPER – SOX2 (MA0143.3), OCT4 (MA0142.1), NANOG (UN0383.1), and KLF4(MA0039.2)^16^. The position-weighted binding (PWM) score is calculated over the genomic regions in 5’ and 3’ directions, identified using the respective ChIP-seq peaks^17^.

$$PWM= \sum_{i=1}^{n} p(i,N_{i})$$

(5)

where *i* is the position along the consensus sequence and $N_{i}$ is the identity of the DNA nucleotide at position *i*. The transcription factors are modeled onto the top 20 genomic locations ranked based on the PWM score in a strand-specific manner.

2.5 Sequence analysis of the positional maps

The DNA sequence of the *Nanog* gene locus (Chr6: 122600-122800 kb) and the annotations are obtained from the UCSC genome browser (Accession ID: GCA_000001305.2; mm10)^18^. The generated protein maps are consistent with the mm10 mouse genome assembly, and the sequences are directly extracted from the corresponding genomic locations. The A/T fraction is the probability of A or T at a specific position (-73 to 73 bp) relative to the dyad (set to 0) calculated from 1002 mapped nucleosome positions. Similarly, (A/T)_5_ probability is the frequency of finding a 5-mer composed solely of combinations of A and T (e.g., ATATA, ATTTT, TTAAA, etc.) centered at a given position. A_5_/T_5_ probability is the chance of finding 5-mer composed of A or T alone (e.g., AAAAA and TTTTT) centered at a given position. The A/T fraction as the function of H1-binding probability is calculated by binning the nucleosomes based on the H1-binding probability calculated in 2.2.3, and the distribution of the A/T fraction is calculated for each bin.

The shift in the selected nucleosome positions compared to 50 independent simulated annealing simulations are calculated as follows,

$${Nuc. Shift}_{i,j}=min\left( \left| {Dyad Pos}_{nuc=i}^{Ref}-{Dyad Pos.}_{nuc=0 to n}^{Run=j} \right| \right)$$

(6)

here, ${Dyad Pos}_{nuc=i}^{Ref}$ is the position of nucleosome *i* used in the modeling and ${Dyad Pos.}_{nuc=0 to n}^{Run=j}$ is the set of nucleosome positions obtained from the *j^th^* run.

3. Protein Structure Modeling

The structures of the mouse embryonic stem cell (mESC)-specific protein complexes are modeled through homology modeling by MODELLER and ab-initio models predicted by Alphafold^19–21^. The mESC-specific nucleosome is modeled by aligning appropriate sequences – H3.3 (P84244), H4 (P62806), H2A.1B (C0HKE1), and H2B.3A (Q9D2U9) - to the nucleosome structure of Xenopus laevis (1KX5) using MODELLER. Similarly, the dominant linker histone variant - H1.3 (P43277) - in mESC is modeled at the dyad position using the structure of human nucleosome with linker histone as the template (7K5Y)^12^.

The BRD4 (Q9ESU6) structure from the Alphafold protein structure database is modeled to bind the H3 tail of the nucleosome based on the BRD4-H3 tail structure (2WP1) as the template. The structure of SOX2 (P48432), OCT4 (P20263), NANOG (Q80Z64), and KLF4 (Q60793) are modeled using 1GT0, 3L1P, 2VI6, and 4M9E structures as templates, respectively^22^, and the structure of P300 (B2RWS6) from the Alphafold protein structure database is used. The modeled structures contain an extensive array of disordered tails (at the protein terminus and interdomain segments), either identified by the missing structural information from the template structures or with a low pLDDT score of Alphafold predicted structures. The modeled structures of these segments are not well-defined; however, they provide an initial configuration suitable for subsequent simulation studies.

The cryo-EM structure of the human transcription pre-initiation complex (PIC; 7ENC) comprising 25 Mediator subunits, 12 RNA polymerase II subunits, and the components of general transcription factors TFIIA, TFIIB, TFIID, TFIIE, TFIIF, and TFIIH, is used as a template to model mouse PIC. The template is supplemented with the structural information from the atomic model of the mouse Mediator complex (6W1S). Table S2 lists the uniport IDs of the protein sequences used to model the mouse PIC.

The modeling of mouse PIC is performed in a series of steps to make minimal perturbations to the experimental structures owing to the homology between mouse and human protein sequences.

Step 1: The 7ENC structure does not include the MED25 subunit and contains partial structural information of the MED14 and MED16 subunits available in 6W1S. We created a new structural template by aligning the MED23-MED25 subunit from 6W1S onto 7ENC using the MED23 structure as a reference. The structural overlap and clashes are resolved by loop modeling by MODELLER and positional adjustments using PyMOL (The PyMOL Molecular Graphics System, Version 2.0 Schrödinger, LLC.). Similarly, the MED14 and MED16 from 6W1S are aligned to 7ENC to include the complete structural information of the Mediator subunits. The combined PIC structure from 7ENC and 6W1S is used as the template for further modeling.

Step 2: The missing atomic details in the template are modeled using SCRWL4^23^. About 825 residues with missing side chain information are fixed.

Step 3: The residues on the template structure are mutated to match the mouse sequence using PyMOL based on the local pairwise sequence alignment.

Step 4: Modeling the longer disordered regions consistent with higher order organization of PIC is not straightforward, as the loops modeled without a template often result in physiologically restricted topological loops and knots. To overcome this bottleneck, we trimmed the protein segments without templates to the length of 14 amino acids by combining 7 amino acids, flanking the termini of the segment. Assuming the additional two amino acids on the termini of the template to be flexible, the disordered segments are modeled using the loop modeling protocol from MODELLER. The modeled trimmed loops are visually analyzed for topological loops and knots and are resolved by further loop modeling efforts.

Step 5: The complete model of the subunits suspected to participate in the liquid-liquid phase separation is generated selectively based on the previous literature reports^24,25^. The selectively modeled subunits are MED1, MED14, MED15, RNA Pol II, TFIID3, TAF4(2), TFIID5(2), TFIID11. The structure of the protein segments that are larger than 40 amino acids are generated using Alphafold and are appropriately incorporated into the model to account for any missing residual structure information. The structural clashes and knots are resolved by loop modeling protocol or manual adjustments using the PyMOL.

All the modeled structures are energy-minimized using Gromacs^26^ employing amber ff99SB*-ILDN forcefield^27^. The structures are subsequently coarse-grained to residue-level representations using the AICG2+ model for protein and the 3SPN2.C model for nucleic acids implemented in the GENESIS package^28^.

4. Backmapping mesoscopic model to near-atomistic resolution

The near-atomistic model of the *Nanog* gene locus is built by combining positional maps of protein complexes and the five representative conformations from the mesoscopic chromatin model. The backmapping pipeline is a series of streamlined procedures, as described in Figure 6 in the main text and Figure S10.

Step 1: The genome segment of interest (mESC chr 6: 122600-122800 kb) can be represented as a chromatin fiber made up of fiber modules – (a) nucleosome made up of histone octamer wrapped by 147 bp of DNA, (b) nucleosome bound to linker histone at the dyad position (Nuc-H1), (c) nucleosome with BRD4 bound to the H3 tail (Nuc-BRD4), (d) nucleosome bound to both linker histone and BRD4 and (e) transcription pre-initiation complex (PIC) – connected by linker DNA of variable length.

Step 2: Given the fiber modules, the modules are connected by a Monte Carlo-based approach to follow the fiber path defined by the representative conformations from the mesoscopic model. The automated end-to-end modeling of the fiber, described below, is optimized based on several considerations to generate an experimentally consistent initial configuration of the *Nanog* gene locus suitable for subsequent simulation studies.

Step 2a – Preprocessing fiber modules: The CG fiber modules contain extended disordered tails, making it hard to build compact fiber representations without steric clashes. To make the hassle-free initial model, the fiber modules are artificially compacted using a strong HPS interaction ($\varepsilon=$ 0.8, default = 0.2)^29^ for disordered tails at low ionic strength (10 mM) and temperature (278 K) in GENESIS (Figure S10B). Disordered tails are identified as protein segments without structural information in the template structures and residues with pLDDT score < 0.5 for Alphafold modeled structures (section 2.3). The charges on the DNA phosphate are set to -1.0 further to facilitate the interaction of charged tails with DNA. Conformations with little or no tail interactions with the DNA terminus (up to 25 bp of DNA) are chosen for end-to-end fiber modeling.

Step 2b – Preprocessing mesoscopic fiber: The representative mesoscopic configurations with 1 bead diameter (in reduced units) are energy minimized using GENESIS to remove the overlapping non-local beads (due to the Weeks-Chandler-Anderson potential) using the reference bond length, angle, and pair-wise native interactions defined by the corresponding configurations.

Step 2c – Generating structural ensemble of fiber modules: The ensembles of nucleosome containing fiber modules (fiber modules a-d described in step 1) are generated by trimming the terminus of nucleosomal DNA and combining it with the fragments of straight B-DNA, generated using web 3DNA^30^, of variable length. Two straight DNA fragments are combined with the nucleosomal DNA by aligning the terminal phosphates (3 phosphates of both strands at the termini of DNA; Figure S10C) to generate ensembles representing DNA unwrapping and nucleosome sliding (by extending DNA length on one end compared to the other; Figure S10D). The unwrapping length and nucleosome sliding are sampled from the normal distribution as follows,

$$length of unwrapped DNA=\left\lceil\left| n_{bp,U}^{max}\times p\left( normal, \mu=0, \sigma=1 \right) \right| \right\rceil\% n_{bp,U}^{max}$$

(7)

$$Sliding Length=\left\lceil n_{bp,S}^{max} \times p\left( normal,\mu=0, \sigma=1 \right) \right\rceil\% n_{bp,S}^{max}$$

(8)

where, $p$ is the probability drawn from the normal distribution with mentioned mean ($\mu$) and standard deviation ($\sigma$), % operation gives the reminder, $n_{bp,U}^{max}$ and $n_{bp,S}^{max}$ represent the maximum length of unwrapped DNA and nucleosome sliding sampled during the ensemble generation and are set to 25 and 5bp, respectively. The limits are defined based on the estimates available in the literature^31–33^.

The ensemble of linker DNA is generated by combining two straight DNA with a fragment of curved nucleosomal DNA, as previously mentioned. The variable length of the three DNA fragments determines the relative distance and orientation of the fiber modules added at the ends (Figure S10D). During the conformational sampling, only one bent DNA is added for each linker DNA segment of length $n_{bp}^{l,DNA}$ as follows,

$$l_{bent DNA}= \left\lceil n_{bp}^{l,DNA}\times p\left( normal,\mu=0, \sigma=1 \right) \right\rceil\% n_{bp}^{l,DNA}$$

(9)

$$l_{straight DNA 1}= \left\lceil\left( n_{bp}^{l,DNA}-l_{bent DNA} \right)\times p(uniform) \right\rceil$$

(10)

$$l_{straight DNA 2}= n_{bp}^{l,DNA}-l_{bent DNA}-l_{straight DNA 1}$$

(11)

where, $n_{bp}^{l,DNA}$ is the length of the linker DNA segment, p is the probability drawn from the normal distribution with mentioned mean ($\mu$) and standard deviation ($\sigma$) for sampling the length of bent DNA and the probability drawn from a uniform distribution for sampling the length of straight DNA, and % operation gives the reminder. In the case of linker DNA length >167 bp, a nucleosome decoy is added during the modeling and treated equivalent to the nucleosome fiber module without the protein components.

The PIC ensemble is generated by combining five fragments of DNA – straight DNA fragment + curved nuc. DNA fragment + DNA fragment from PIC model (60 bp modeled using 7ENC as the template; refer section 2.3) + curved nuc. DNA fragment + straight DNA fragment. The sharp bending of DNA around the PIC complex facilitates the flexible modeling of the PIC ensemble together with nucleosome modules.

Step 2d – Conformational Sampling and Scoring: The fiber is modeled end-to-end by sampling the conformations of two nucleosome-containing modules connected by a linker DNA at each instance for a predefined number of iterations. The conformations are selected by the fitness score (F) that minimizes the distance of the nucleosome modules from the reference fiber path, the compactness of the sampled conformation, and the distance between i-i+2 nucleosomes.

$$F_{i=1}^{n}= \left\langle\left[ d1\left( R_{nuc,i}^{com},R_{nuc,i}^{ref} \right),d1\left( R_{nuc,i+1}^{com},R_{nuc,i+1}^{ref} \right),\max\left( d\left( R_{i}^{conf},R_{nuc,i}^{ref} \right) \right), d\left( R_{nuc,i-1}^{com},R_{nuc,i+1}^{com} \right) \right] \right\rangle$$

(12)

here, function $d(R_{x},R_{y})$ calculates the pairwise distance between two sets of coordinates, $R_{nuc,x}^{com}$ is the center of mass coordinate of the nucleosome $x$ sampled during the modeling, $R_{nuc,x}^{ref}$ is the reference coordinate calculated based on the representative conformation from the mesoscopic model, $R_{i}^{conf}$is the coordinates of the conformation (${Nuc. module}_{i}+{linker DNA}_{i}^{i+1}+{Nuc. module}_{i+1}$) sampled at instance $i$ and $n$ is the total number of nucleosome containing modules. The triangular brackets indicate the average over the four distances that are being optimized. The $d1$ function is calculated as,

$$d1\left( R_{nuc,i}^{com},R_{nuc,i}^{ref} \right)=\max\left[ {f_{width}}/3,d\left( R_{nuc,i}^{com},R_{nuc,i}^{ref} \right) \right]$$

(13)

here, $f_{width}$ is the fiber width set to 22 nm equivalent to the bead diameters assumed in the mesoscopic model, and $R_{nuc,x}^{ref}$ is calculated as the linear projection of the genomic location of the nucleosome $x$ considering each bead in the mesoscopic model correspond to 1 kb chromatin segment.

$$R_{nuc,i}^{ref}= \left[ R_{x}^{meso}+\left( x-{Loc}_{i} \right) R_{x+1}^{meso} \right]\times f_{width}$$

(14)

$$x=\left\lfloor{Loc}_{i}/1000 \right\rfloor$$

(15)

here, $R_{x}^{meso}$ is the coordinate of the bead $x$ from the representative mesoscopic conformation and ${Loc}_{i}$ is the genomic location of the fiber module $i$ indexed from 0 for the modeled genomic segment.

This treatment of the $d1$ function allows our modeling pipeline to avoid overfitting the nucleosome modules to the reference fiber at the expense of compactness and *i*-*i*+2 interactions. For each instance $i$, the conformations are sampled for 1000-3000 iterations, and the conformation with minimal fitness score and minimal steric clashes with the conformation generated up till $i-1$ instance. The conformation that forms local knots, with conformation upstream 5kb, are identified and ignored during the sampling procedure^34^. Sampling nucleosome modules *i* and *i*+1 together ensures that the chromatin model grows in the desired direction of the reference fiber and the linker DNA orientation favors the addition of consecutive nucleosome modules. However, only the ${NCP module}_{i}$ and ${linker DNA}_{i}^{i+1}$ is added to the chromatin model and the conformation of ${NCP module}_{i+1}$ is sampled again during the $i+1$ instance.

The size of the compacted PIC complex is >22 nm and is larger than the diameter of the mesoscopic fiber beads. Hence, additional care is taken when adding the PIC module to the chromatin model. At instance $i$, if the $i+2$ module is a PIC module, then in addition to the two nucleosome modules sampled during the other instances, the PIC ensemble is sampled as described in the previous step. In this case, the fitness score also includes the distance of the center of mass of the modeled PIC-DNA from the corresponding reference coordinate calculated from the fiber path.

$$F_{i}^{PIC \in i+2}=\left[ F_{i}+d1\left( R_{PIC,i+2}^{com},R_{PIC,i+2}^{ref} \right) \right]/2$$

(16)

here, $R_{PIC,i+2}^{ref}$ is calculated the same as $R_{nuc,i}^{ref}$ as described before. Finally, the entire $R_{i}^{conf}$is added to the chromatin model, unlike other instances, to accommodate the PIC module alongside the two nucleosome modules.

Step 2e: The modeled chromatin model at near-atomistic resolution is analyzed for steric clashes and the distribution of the distance between the reference mesoscopic fiber and the modeled fiber module. Since the conformation at each instance is sampled for a predefined number of iterations (generally, 2000 iterations but 3000 iterations while modeling PIC), it is still possible to generate chromatin models with steric clashes, especially with the PIC molecules or in compact mesoscopic configurations. To generate a better model, at least ten independent models are generated for each representative mesoscopic conformation and the model with least steric clashes and best fit are selected. The double-stranded DNA in the final chromatin model is coarse-grained to a single chain polymer with 25 bp resolution and are analyzed for knots using KymoKnot software package^35^. The best chromatin models for each representative mesoscopic conformation are used for further modeling.

Step 3: Modeling transcription factors: The transcription factors (SOX2, OCT4, NANOG, KLF4) and cofactor (P300) are later added to the chromatin models near the identified genomic locations.

Step 3a: Preprocessing transcription (co)factors: The transcription factors are artificially compacted, in the presence of short DNA fragments (8 bp) with a strong positional restraint for DNA and transcription factor globular domain, using strong HPS interaction ($\varepsilon=$ 0.8, default = 0.2)^29^ for disordered tails at low ionic strength (10 mM) and temperature (278 K) in GENESIS.

Step 3b: The initial model is generated by aligning the transcription factor-bound DNA strand in the preprocessed model to the mapped genomic location. The molecules are translational- and orientationally adjusted iteratively to minimize clashes and the distance between the center of mass of DNA-binding residues and the center of mass of the mapped DNA segment. The process is repeated $n$ times for each transcription factor. In the case of P300, the initial model is generated by randomly positioning the compacted P300 model at the desired location and progressively minimizing the clashes and the distance to the center of mass of the mapped location.

Step 4: Energy minimization and CG implicit solvent MD simulation: The model is energy minimized and subsequently relaxed, employing the AICG2+ model for proteins and the 3SPN.2C model for double-stranded DNA in GENESIS^28,36,37^. The native interactions of the globular domains are defined based on the modeled structures (refer to section 2.3). The native interactions in the linker histone globular domain are 1.05 times stronger than the AICG2+ model to stabilize the tertiary structure. The disordered segments, identified as those not resolved in the template structures or with pLDDT<0.5 for Alphofold models, are simulated using a statistical potential based on the comprehensive loop-structure-library^38^ with solvation interactions defined by the HPS model^29^. The inter-domain interactions in the Alphafold predicted structures are identified based on the predicted aligned error (PAE) and native interactions are defined between residues with PAE$\leq$5Å.

The interaction between the core histone proteins and DNA is modeled using the non-specific hydrogen-bond-like interaction as previously done with the energy constant set to 2.7 k_B_T^33^. Similarly, linker histone CG residues within 4Å from the DNA in the modeled structure interact with DNA using the hydrogen-bond-like interaction. The DNA binding residues of transcription factors are identified using the reference structures (SOX2:1GT0, OCT4:3L1P, NANOG:2VI6 and KLF4:2WBU) and interact with DNA via a sequence-specific PWMcos potential^39^. The parameters of the PWMcos potential are set to $\gamma$=2.0 and $\varepsilon$=-0.5 to facilitate strong cognate sequence bias and interaction with DNA, respectively. BRD4 is connected to the H3 N-terminal tail using a virtual bond to retain its association with the mapped nucleosome.

The charge on the DNA phosphates is set to -1.0, and the charged residues on the protein have integer charges corresponding to pH 7. The system is energy minimized at 150 mM ionic strength conditions and 300 K. The MD simulations are conducted by Langevin dynamics for three million MD steps, with the friction coefficient $\gamma$=0.01, using GENESIS CGDYN^37^ in the Fugaku supercomputer.

| Protein/Protein Complex | Experimental Data | GEO accession | Reference Genome Assembly | Source |
| --- | --- | --- | --- | --- |
| Nucleosome | Chemical Mapping | GSE82127 | mm10 | ^40^ |
| RNA polymerase | ChIP-seq | GSM22562 | mm8 | ^41^ |
| Linker Histone | ChIP-seq | GSE46134 | mm9 | ^11^ |
| BRD4 (cofactor) | ChIP-seq | GSE87064 | mm9 | ^42^ |
| P300 (cofactor) | ChIP-seq | GSE29218 | mm9 | ^43^ |
| Transcription Factors – SOX2, OCT4, NANOG, KLF4 | ChIP-seq | GSE90985 | mm9 | ^17^ |
| Epigenetic Marks – H3K27Ac, H3K9Ac | ChIP-seq | GSE90985 | mm9 | ^17^ |
| Cohesin components – SMC3, SMC1, CTCF | ChIP-seq | GSE22562 | mm8 | ^41^ |
| Mediator Complex – Med1, Med12 | ChIP-seq | GSE22562 | mm8 | ^41^ |

Table S1: Experimental positional maps used for mapping proteins and protein complexes

| Protein Complex | Uniprot ID | Protein | Uniprot ID | Protein |
| --- | --- | --- | --- | --- |
| Mediator | Q925J9 | MED1 | Q8C1S0 | MED19 |
|  | Q9CQA5 | MED4 | Q9R0X0 | MED20 |
|  | Q921D4 | MED6 | Q9CQ39 | MED21 |
|  | Q9CZB6 | MED7 | Q62276 | MED22 |
|  | Q9D7W5 | MED8 | Q80YQ2 | MED23 |
|  | Q8VCS6 | MED9 | Q99K74 | MED24 |
|  | Q9CXU0 | MED10 | Q8VCB2 | MED25 |
|  | Q9D8C6 | MED11 | Q7TN02 | MED26 |
|  | A2ABV5 | MED14 | Q9DB40 | MED27 |
|  | Q924H2 | MED15 | Q920D3 | MED28 |
|  | Q6PGF3 | MED16 | Q9DB91 | MED29 |
|  | Q8VCD5 | MED17 | Q9CQI9 | MED30 |
|  | Q9CZ82 | MED18 | Q9CXU1 | MED31 |
| RNA Polymerase II | P08775 | RPB1 | Q8CFI7 | RPB2 |
|  | P97760 | RPB3 | Q9D7M8 | RPB4 |
|  | Q80UW8 | RPAB1 | P61219 | RPAB2 |
|  | P62488 | RPB7 | Q923G2 | RPAB3 |
|  | P60898 | RPB9 | P62876 | RPAB5 |
|  | O08740 | RPB11 | Q63871 | RPAB4 |
| General Transcription Factors | P29037 | TBP | Q99PM3 | TFIIA1 |
|  | Q80ZM7 | TFIIA2 | P62915 | TFIIB |
|  | Q9D0D5 | TFIIE1 | Q9D902 | TFIIE2 |
|  | Q3THK3 | TFIIF1 | Q8R0A0 | TFIIF2 |
|  | P49135 | TFIIH -XPB | O08811 | TFIIH - XPD |
|  | Q9DBA9 | TFIIH1 | Q9JIB4 | TFIIH2 |
|  | Q8VD76 | TFIIH3 | O70422 | TFIIH4 |
|  | Q8K2X8 | TFIIH5 | P51949 | MAT1 |
|  | Q03147 | CDK7 | Q61458 | Cyclin-H |
|  | Q80UV9 | TFIID1 | Q8C176 | TFIID2 |
|  | Q5HZG4 | TFIID3 | E9QAP7 (2) | TAF4 |
|  | Q8C092 (2) | TFIID5 | Q62311 (2) | TFIID6 |
|  | Q9R1C0 | TFIID7 | Q9EQH4 | TFIID8 |
|  | Q8VI33 (2) | TFIID9 | Q8K0H5 (2) | TFIID10 |
|  | Q99JX1 | TFIID11 | Q8VE65 (2) | TFIID12 |
|  | P61216 | TFIID13 |  |  |

Table S2: UniProt sequence identifiers of the protein components constituting mouse transcription pre-initiation complex (PIC).

| Cluster ID | P1-E1 | P2-E1 | P3-E1 | P1-P2 | P2-P3 | P1-P3 | P3-E3 |
| --- | --- | --- | --- | --- | --- | --- | --- |
| 1 | 142 ± 54 | 122 ± 46 | 128 ± 45 | 155 ± 60 | 122 ± 47 | 163 ± 58 | 102 ± 42 |
| 2 | 174 ± 64 | 131 ± 52 | 144 ± 53 | 203 ± 76 | 130 ± 52 | 213 ± 73 | 107 ± 46 |
| 3 | 151 ± 58 | 152 ± 61 | 193 ± 57 | 163 ± 66 | 159 ± 57 | 182 ± 67 | 122 ± 52 |
| 4 | 148 ± 58 | 161 ± 71 | 251 ± 75 | 186 ± 76 | 184 ± 67 | 269 ± 85 | 132 ± 58 |
| 5 | 213 ± 73 | 176 ± 76 | 205 ± 83 | 303 ± 97 | 142 ± 57 | 330 ± 95 | 113 ± 50 |

Table S3. Ensemble-averaged CRE distances (in nm) in the five clusters identified using K-means clustering of Hi-C metainference simulation, and the errors indicate standard deviation. The *Gdf3*, *Nanog*, and *Slc2a3* promoters are P1, P2, and P3, respectively, and -45 SE, -5SE, and +60 SE are indicated as E1, E2 and E3, respectively.

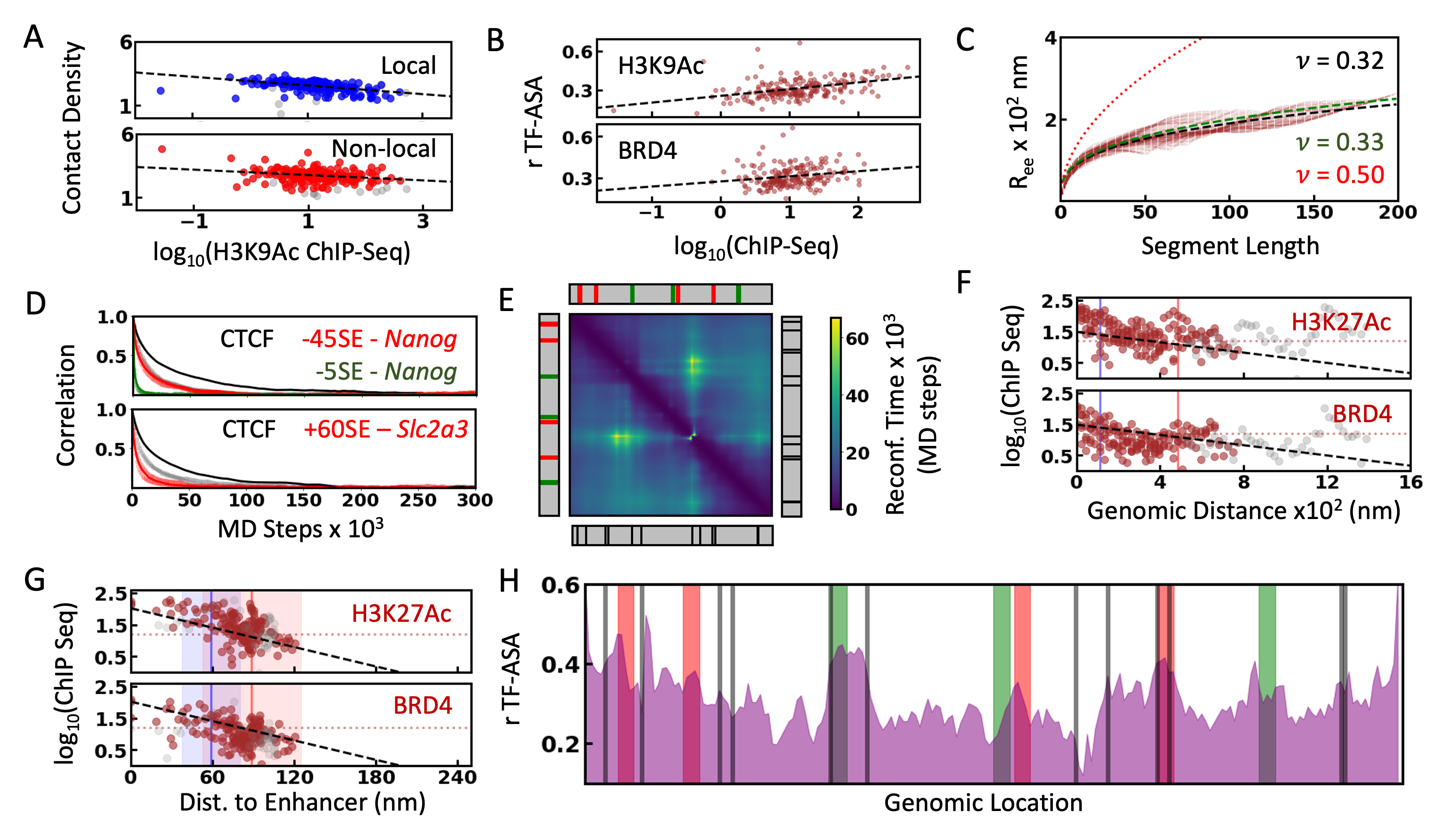

Figure S1: A) Decomposition of the contact density to local ($\leq$5kb) and non-local (>5kb) contact density as the function of log-transformed acetylation signal from ChIP-seq experiments. The colored and grey circles correspond to the 160 kb segment at the 3’ end (122640-122800 kb) and the remaining 40 kb, respectively. Dashed black lines show the linear fit to the data. B) Average relative TF-accessible surface area as the function of log-transformed H3K9Ac and BRD4 ChIP-seq signals. C) Average end-to-end distance (R_ee_) as the function of segment length (L). Black dashed lines are the fit to the data (R_ee_ = aL^ν^), and the reference lines with ν = 0.33 (green) and 0.5 (red) are shown for the same prefactor (a) obtained from the fit. D) Decay of the distance autocorrelation function of CRE pairs (red and green) and the CTCF sites flanking those CREs. The distance autocorrelation function of the randomly selected beads having the same length as the CRE/CTCF pairs are shown as faded circles. E) Reconfiguration time obtained from the single exponential fit to distance autocorrelation function of each mesoscopic bead pairs. F) log-transformed ChIP-seq signals as the function of genomic distance assuming local nucleosome organization (1kb = 22 nm). The color scheme is the same as that of the Figure 2D in the main text. G) Same as figure 2D but assumes the H3K27Ac peak upstream the *Gdf3* promoter as the enhancer in addition to the other three SE. H) Relative TF-ASA as the function of genomic distance. Red and green shaded areas represent the genomic position of promoters and enhancers, respectively. The CTCF binding sites are shown as vertical black lines.

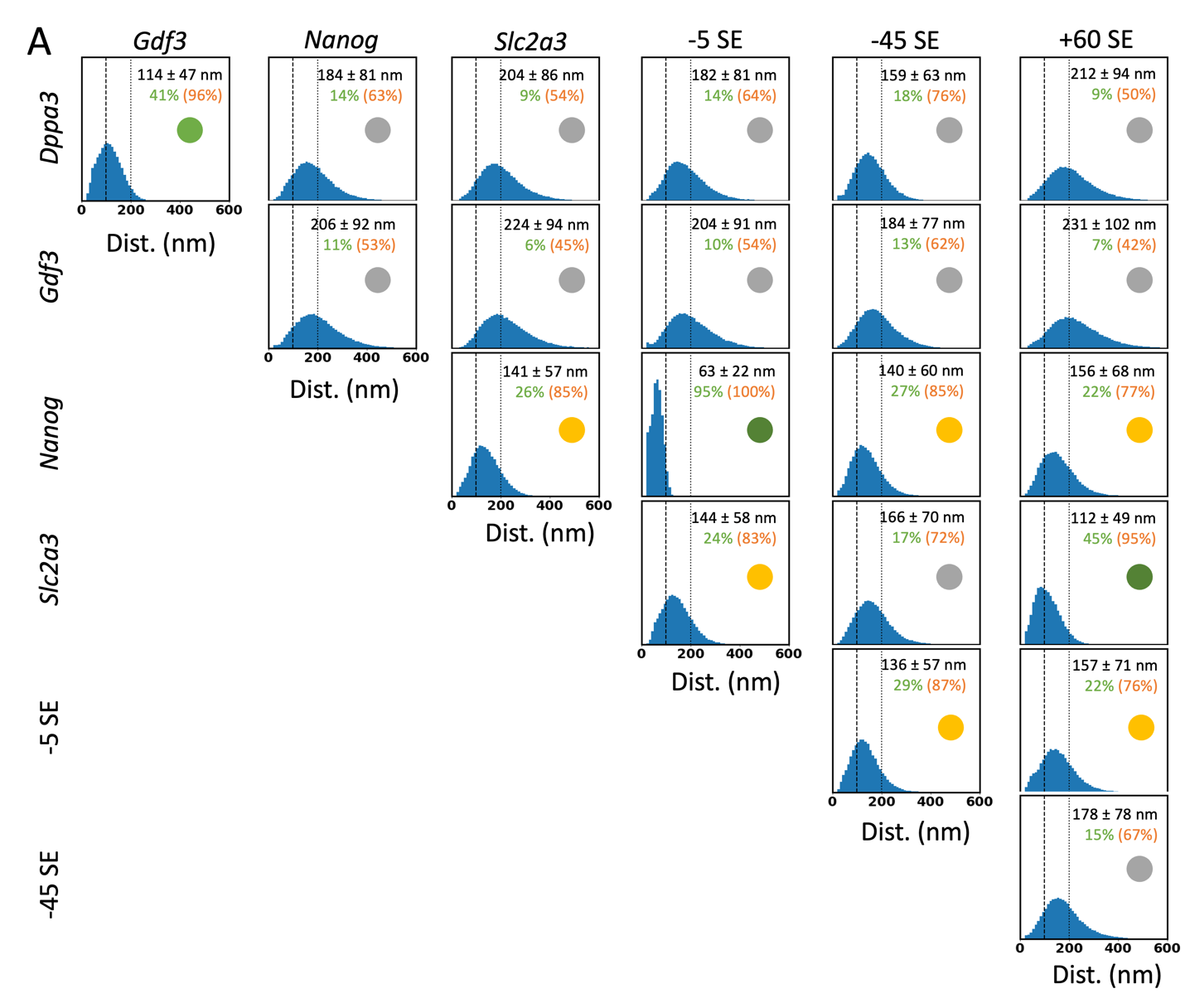

Figure S2: Distribution of pairwise distance between the six cis-regulatory elements (three promoters: *Gdf3*, *Nanog*, and *Slc2a3* and three super-enhancers -45, -5, and +60 SE) in the *Nanog* gene locus. The percentage of conformational ensemble with distances <100 and <200 nm is shown in green and orange, respectively, along with the mean and standard deviation in the top-right corner. The pairs with ~20% and >40% of the distribution <100 nm are marked with green and yellow circles on the top right and are otherwise grey.

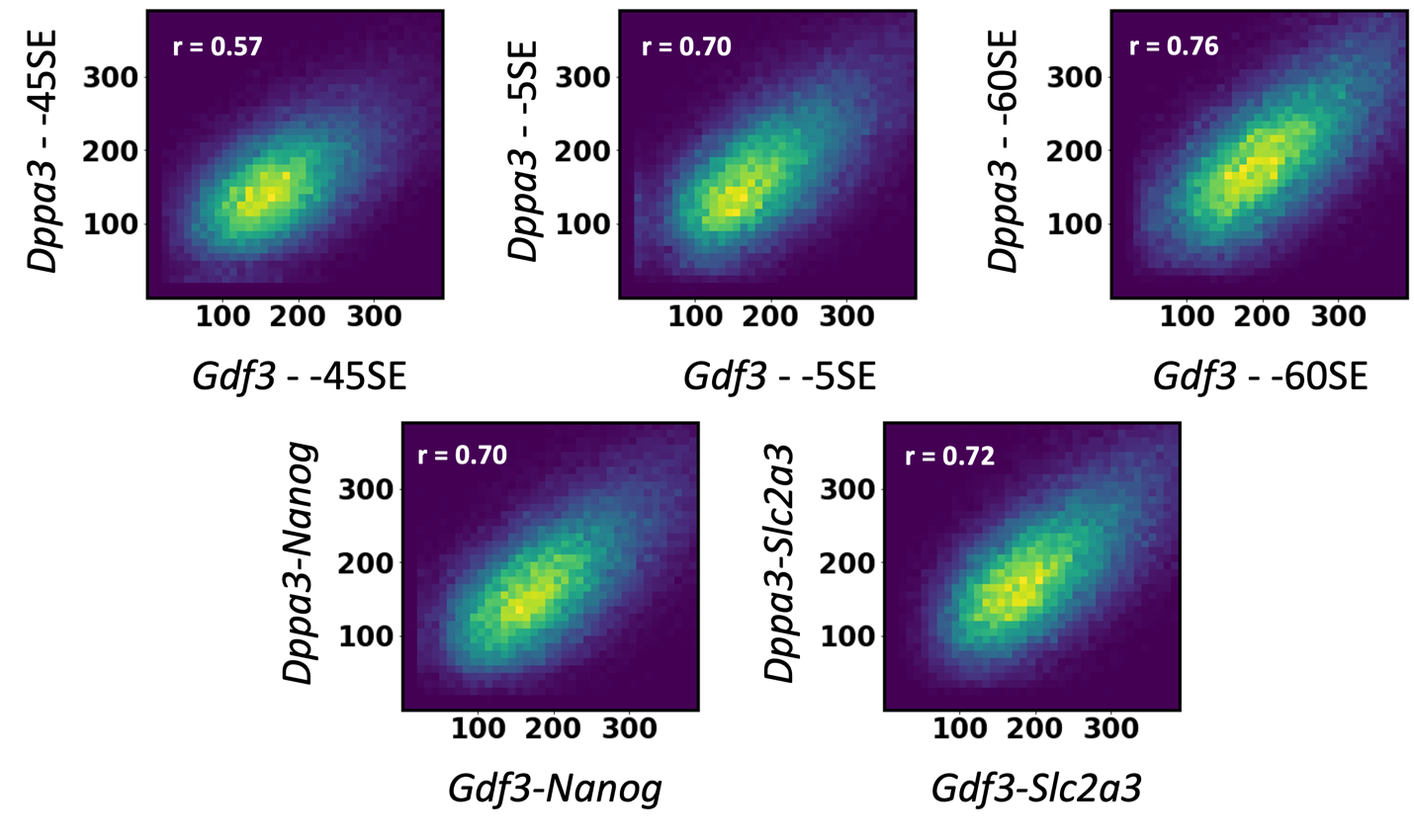

Figure S3: 2D distribution of *Dppa3*-CRE distances and *Gdf3*-CRE distances showing weak positive correlation. The color spectrum of bright to dark corresponds to a decreasing probability.

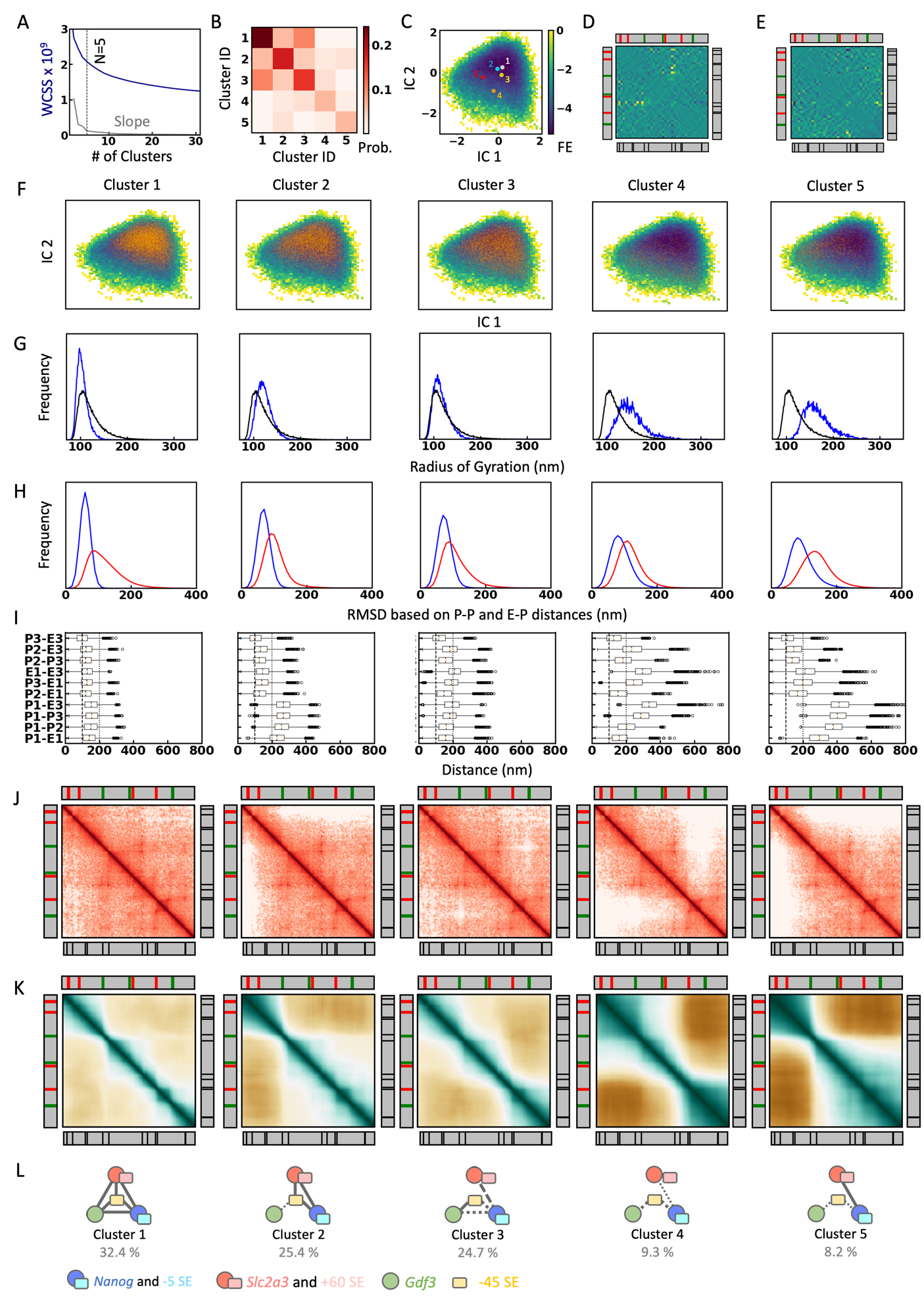

Figure S4: A) Within-Cluster-Sum-of-Squares (WCSS) is calculated as the function of the number of clusters (N) in the k-means clustering. N=5 is selected as the optimal number of clusters corresponding to the elbow point on the slope (grey) of the WCSS curve (blue). B) Cluster transition probabilities calculated from the HiC-metainference simulations. C) Free-energy profile calculated based on the tiCA projections over the first two eigenvectors. The k-means cluster centroids are projected onto the landscape and are numbered in descending order of the cluster population. D and E) independent components 1 and 2 calculated from the tiCA analysis. The color spectrum of bright to dark colors indicates the value range from 1 to -1. The positions of promoters (red), enhancers (green), and CTCF (black) are shown around the margin for reference. F) Projection of k-means cluster data points onto the tiCA landscape displaying the distinct occupancy of individual clusters with hazy boundaries. G) Distribution of radius of gyration of the overall HiC-metainference simulation (black) compared to the distribution of Rg from individual clusters (blue). H) Distribution of pairwise intra– and inter-cluster dRMSD show that the inter-cluster variations are more considerable than intra-cluster differences in the CRE distances. I) Boxplot of the six CRE distances used in the k-means clustering. P1 – *Gdf3* promoter, P2 – *Nanog* promoter, P3 – *Slc2a3* promoter, E1 - -45 SE, E2 - -5SE and E3 - +60SE. J) Average contact maps with intensity proportional to the contact frequency. K) Pairwise correlation map represented using a color spectrum of brown (-1)-white(0)-green(1). The positional maps are the same as in panel D. L) Cartoon representation of clusters, as in Figure 2E.

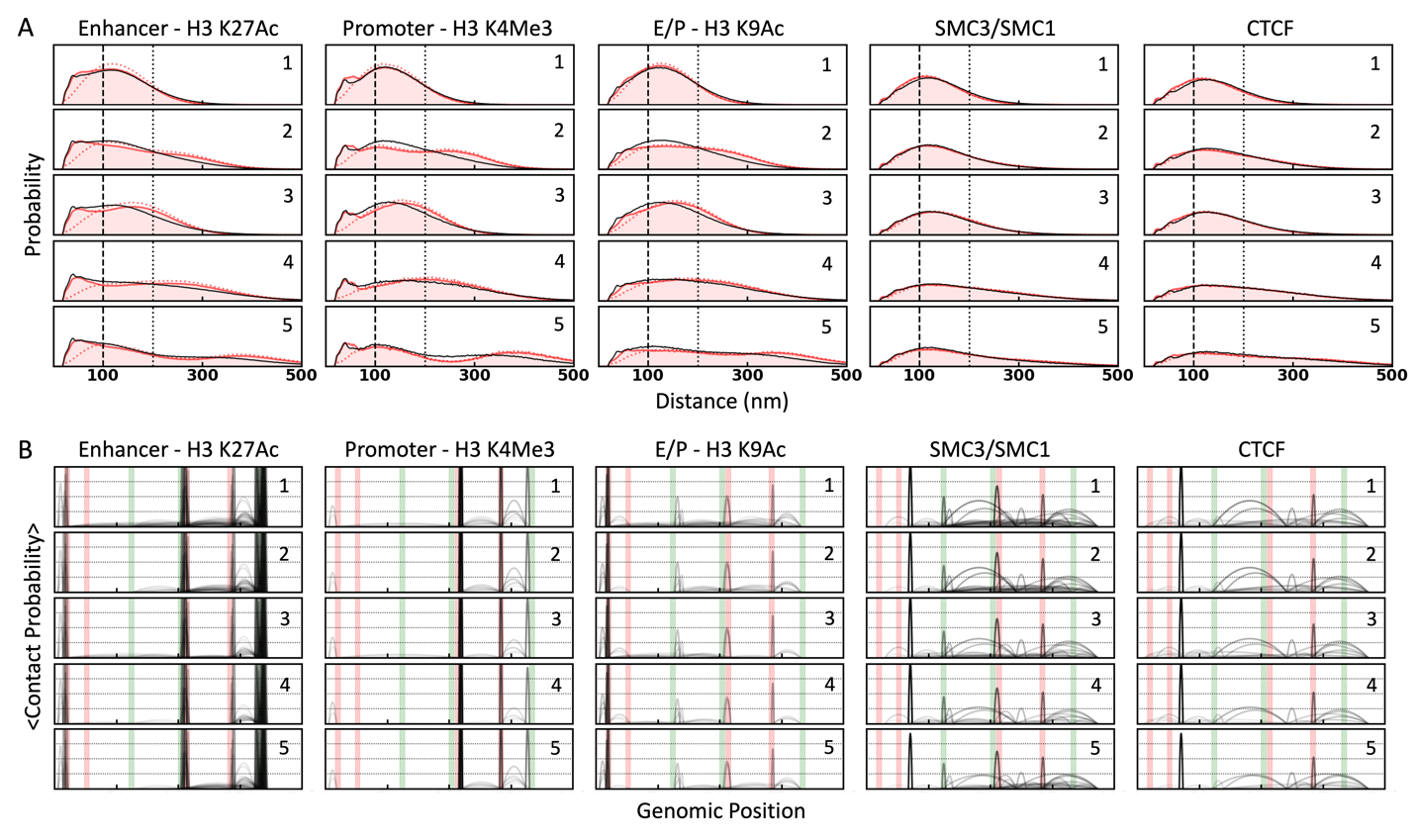

Figure S5: A) Distance distribution of mesoscopic beads, excluding nearest-neighbors (|i-j|>1 in red solid line and |i-j|>5 in red dotted line), enriched with epigenetic markers or protein (z>1). The distance distribution of the randomly selected mesoscopic beads with the same sequence separation as the marker-specific mesoscopic beads is shown as a reference (black solid line). Reference distances 100 and 200 nm are shown as dashed and dotted black lines, respectively. The cluster numbers in the top right corner are the same as in Figure 2E. B) Interaction between mesoscopic beads enriched with epigenetic markers are protein (z>1), where the height and opacity are proportional to the interaction frequency. Positions of the CREs (red – promoters and green – enhancers) are marked as the shaded area for reference.

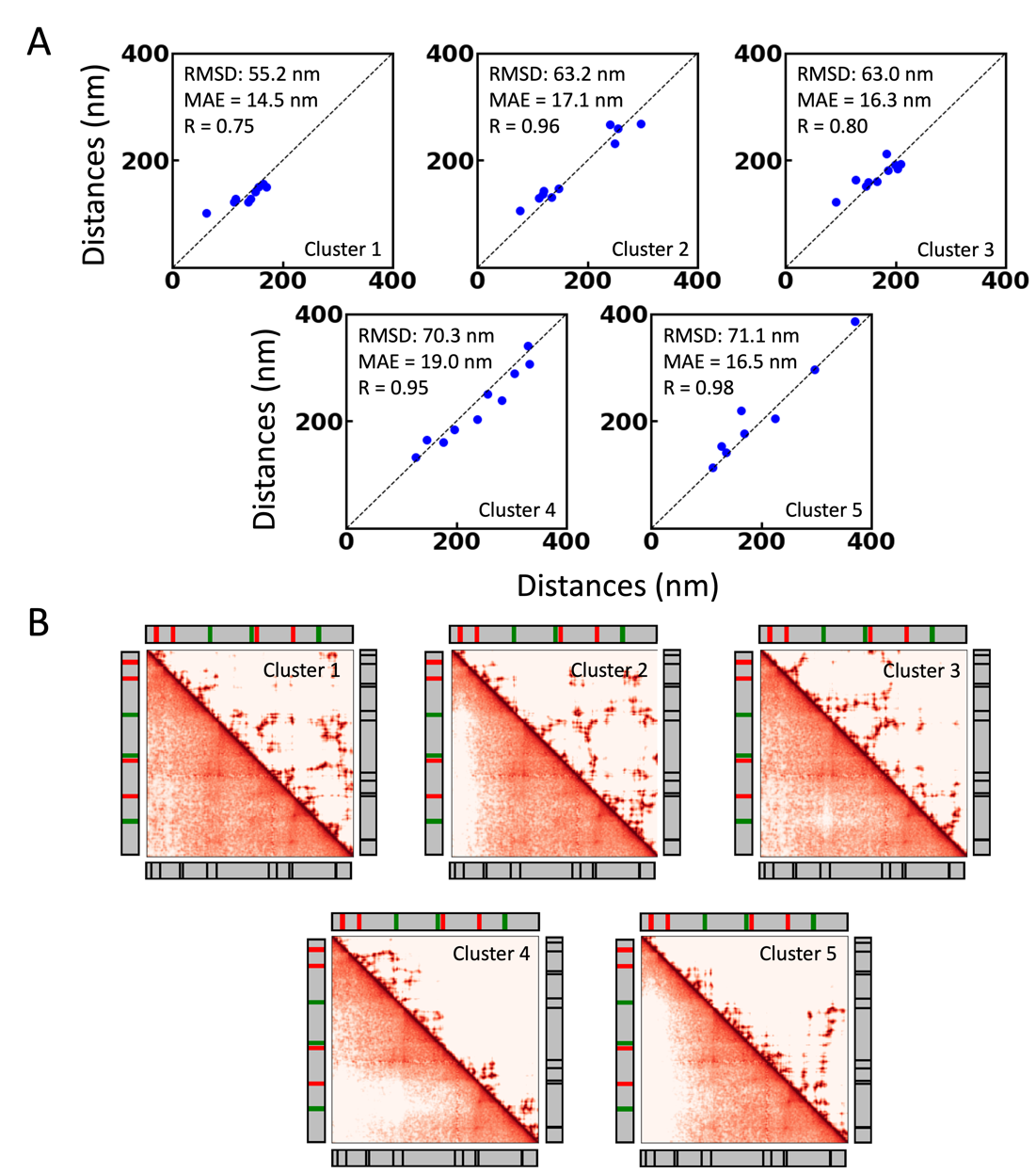

Figure S6: A) Comparison of the CRE distances from the representative structure (x-axis) with the average CRE distances from each cluster (y-axis). The root mean square difference (RMSD), mean absolute error (MAE), and Pearson's correlation (R) are shown in the top-left corner. B) The average compact map of each cluster is shown in the bottom-left, and the contact map of the representative conformation is shown in the top-right. The positions of enhancers (green), promoters (red), and CTCF sites (black) are shown for reference on the side.

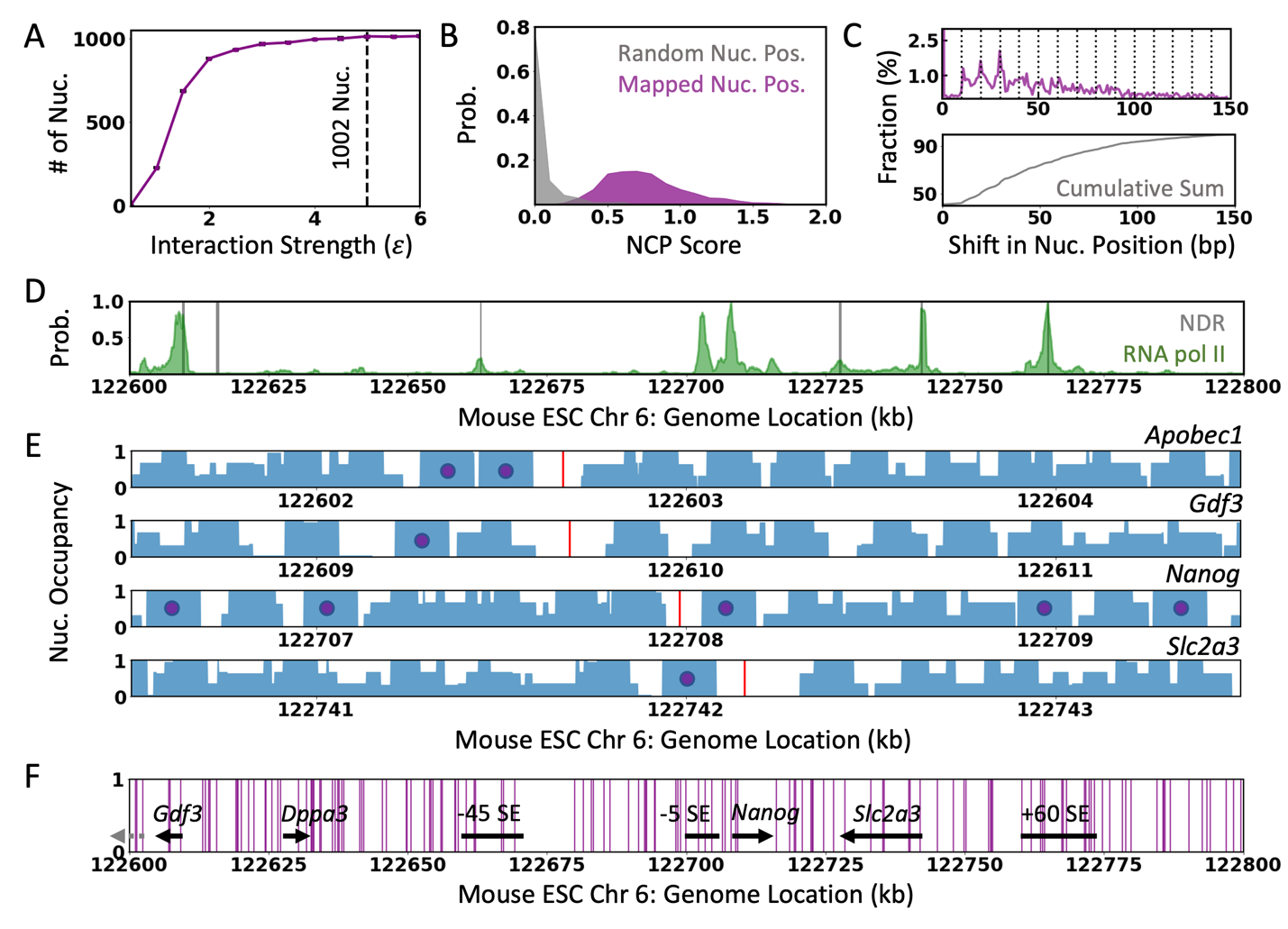

Figure S7: A) Optimizing the proportionality constant in the energy function of Monte Carlo simulations to achieve saturated nucleosome association. The dashed line indicates the final value used to generate the positional maps of nucleosomes. B) Distribution of the nucleosome center positioning (NCP) score at the dyad position of mapped nucleosome positions (purple) and random genomic positions (grey). C) Distribution of the shift in nucleosome positions obtained from multiple independent nucleosome position mapping (top) and the cumulative distribution (bottom). The dashed line indicates 10n periodicity in the nucleosome sliding. D) The position of nucleosome-depleted regions (NDR, >185bp: grey) overlaps with the ChIP-seq densities of the RNA polymerase II (green). E) Probabilistic nucleosome positional map from multiple independent simulations shows well-defined nucleosome positioning (purple circles) in the proximity of the promoters (red). F) Nucleosome positions that are consistently mapped across 50 independent MC runs (purple).

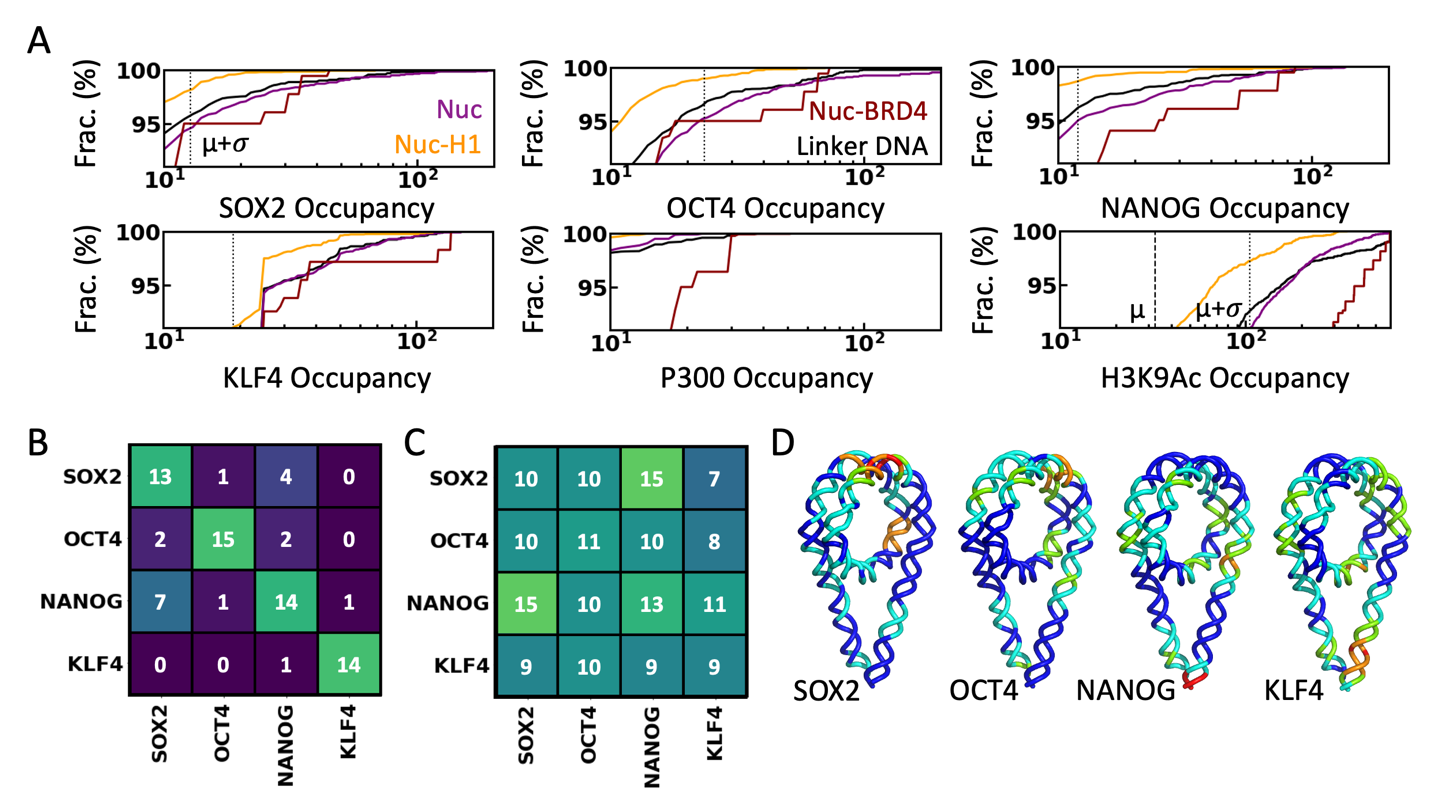

Figure S8: A) Distribution of ChIP-seq occupancy for each nucleosome class and linker DNA as indicated in the first two panels. The dashed and dotted vertical lines indicate the mean ($\mu$) and one standard deviation from the mean ($\mu+\sigma$) calculated for each data within *Nanog* Locus, respectively. B) Frequency of colocalization of the TFs (y-axis) in the same nucleosome mapped with the TF on the x-axis. Numbers on the diagonal indicate the number of TFs mapped onto the nucleosome. The off-diagonal elements are not symmetrical, as multiple TFs are mapped to the same nucleosome. The dark to bright color spectrum of the cells indicates increasing frequency. C) Frequency of TFs (in the y-axis) within 500 bp from the center of the DNA-binding position of the TF in the x-axis. D) Probability of mapped TFs as the function of nucleosomal DNA index mapped onto the nucleosomal DNA for easy visualization.

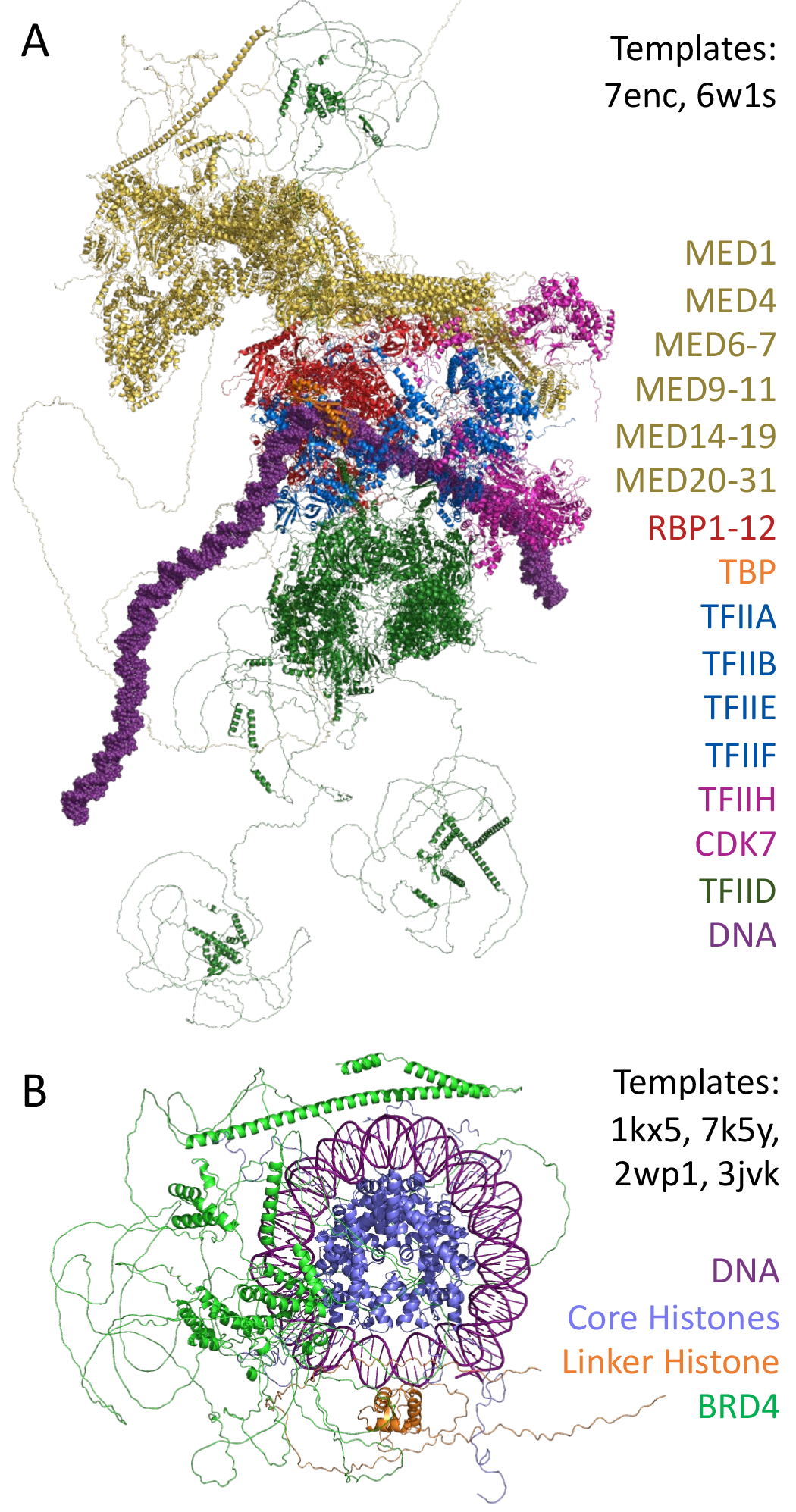

Figure S9: Molecular Modelling of protein complexes. All-atom model of the mouse transcription pre-initiation complex (PIC) (panel A) and mouse nucleosome bound to the linker histone (H1.3) and BRD4 (panel B). The templates used for the modeling are listed, and the subunits are color-coded for reference.

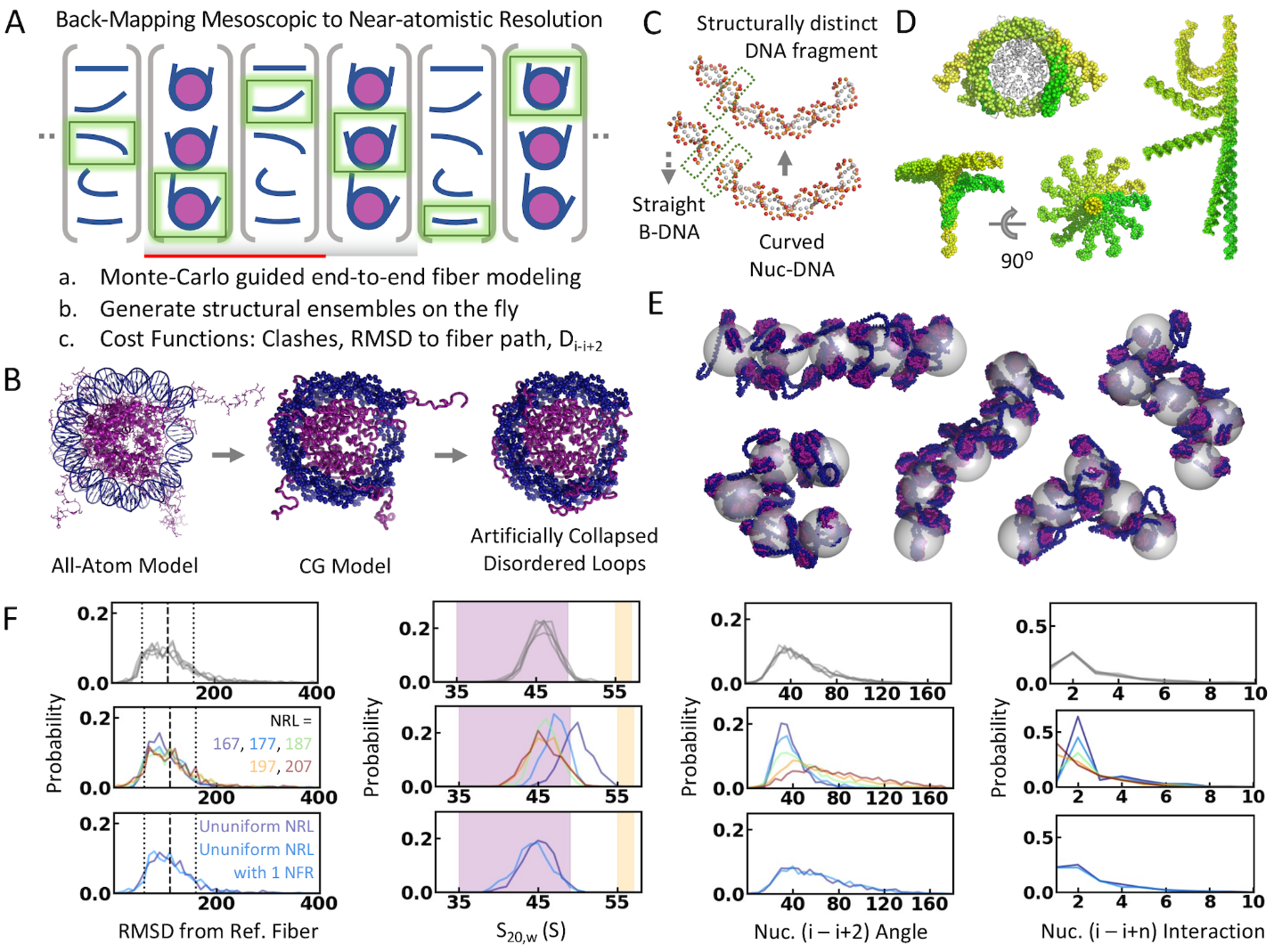
 Figure S10: Mesoscopic to near-atomistic resolution. A) Schematic representation of the backmapping protocol. The chromatin fiber is grown by one nucleosome at a time (red-lined segment) by choosing conformations of modules (grey-shaded segment) from an ensemble that best fits the fiber path. B) The nucleosome modules are coarse-grained, and the tails are compacted for easier modeling. C) The conformational ensemble is generated by combining DNA segments by aligning terminal base pairs. D) Heterogenous nucleosome and linker DNA conformations accessible to the protocol. E) chromatin fiber models at near-atomistic resolution (DNA in blue and nucleosome in purple) based on the structurally diverse 5 kb mesoscopic fibers used for evaluating the backmapping performance (grey). F) Performance of the protocol measured in terms of distance from the reference bead, sedimentation coefficient (S_20,w_), nuci,i+2 angle, and nuci,i+n interactions. The purple and orange shaded area highlights the range of experimental S_20,w_ values observed for 12 nucleosome arrays without and with H1, respectively. The top row shows the performance of 50 chromatin models generated for each representative mesoscopic fiber shown in panel E. Middle and bottom rows correspond to models generated with uniform and random NRLs with and without NFRs. The color codes in each row are the same as the first panel in the respective rows.

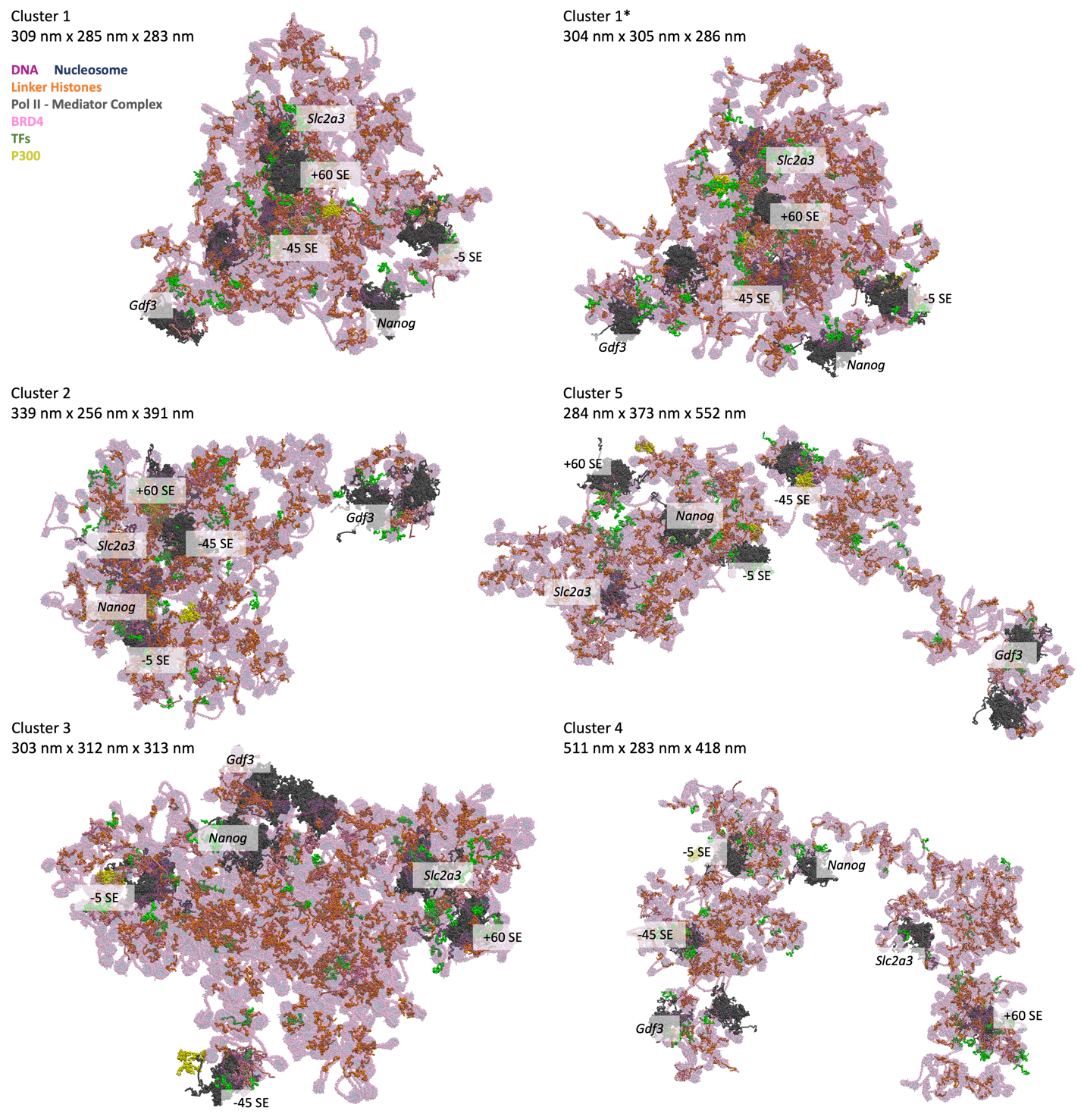
 Figure S11: Generated molecular models of *Nanog* locus at nanoscopic resolution. The models are oriented similarly to the mesoscopic representations in Figure 2E in the main text. Note: Cluster 1* is based on the mesoscopic reference structure of Cluster 1 but generated using an independent backmapping run.

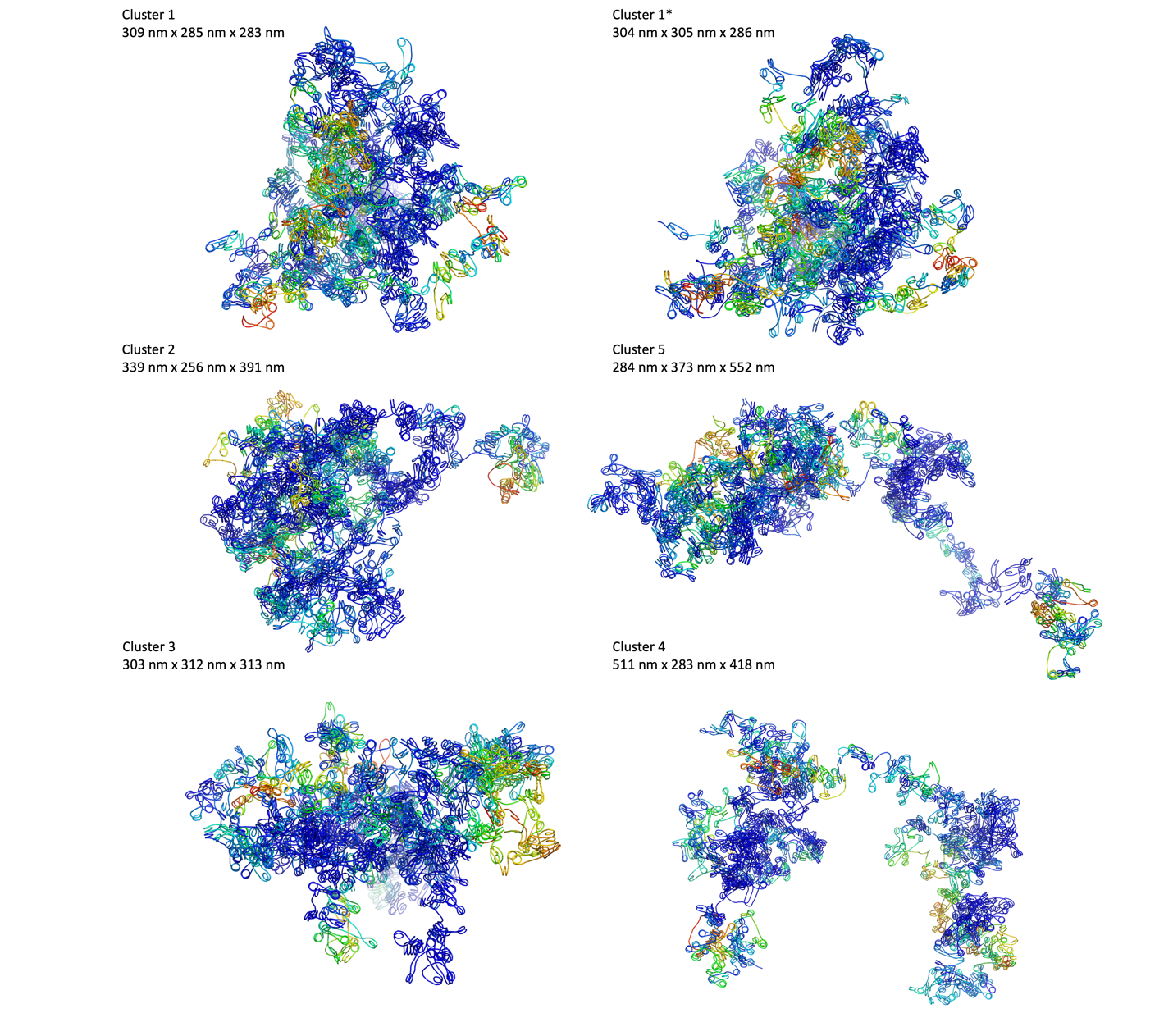

Figure S12: Reduced representation of the generated molecular models of *Nanog* locus. The DNA traces are shown at 20bp resolution and colored proportional to the increasing H3K27Ac signal (blue to red spectrum) for easier visualization. The models are oriented similarly to the mesoscopic representations in Figure 2E in the main text. Note: Cluster 1* is based on the mesoscopic reference structure of Cluster 1 but generated using an independent backmapping run.

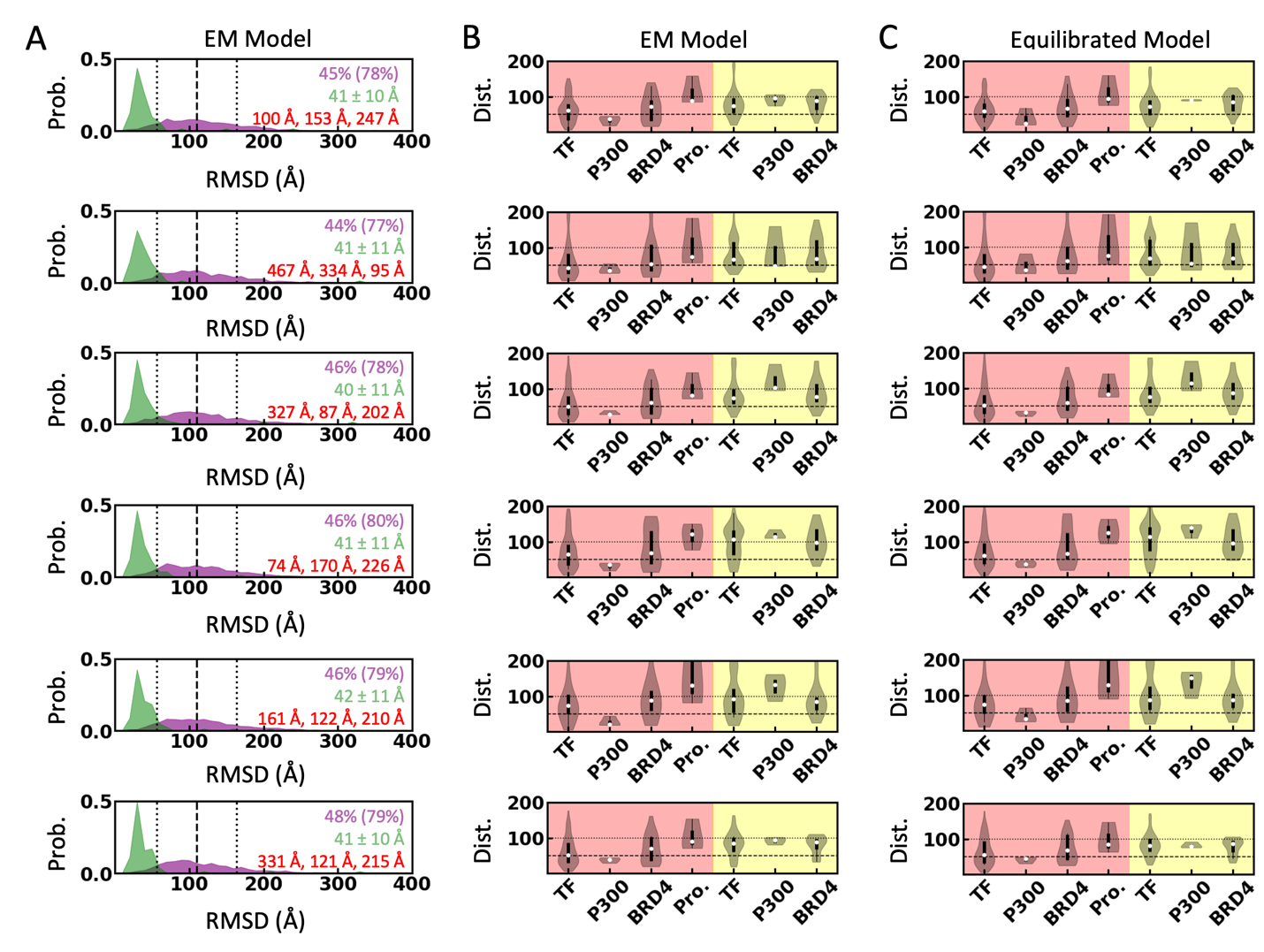

Figure S13: A) Distribution of distance of the modeled nucleosomes (purple) and TFs (green) from their reference mesoscopic bead and DNA binding site, respectively. The radius of the mesoscopic bead (11 nm, black dashed line) ± the radius of the nucleosome (5.5 nm, black dotted lines) are shown as references. The fraction of nucleosomes mapped within 11 (16.5) nm from the reference bead is given in purple, the average distance of the TFs from their corresponding DNA binding site is given in green, and the distance of the three P300 molecules from their target site is given in red. Each row from the top-to-bottom corresponds to clusters 1-5, respectively, and the last row represents Cluster 1*. Note: Cluster 1* is based on the mesoscopic reference structure of Cluster 1 but generated using an independent backmapping run. B and C) Distance distribution of the four TFs, P300, BRD4, and promoters to the nearest enhancers (red shaded) and promoters (yellow shaded) calculated from the energy minimized and equilibrated model.

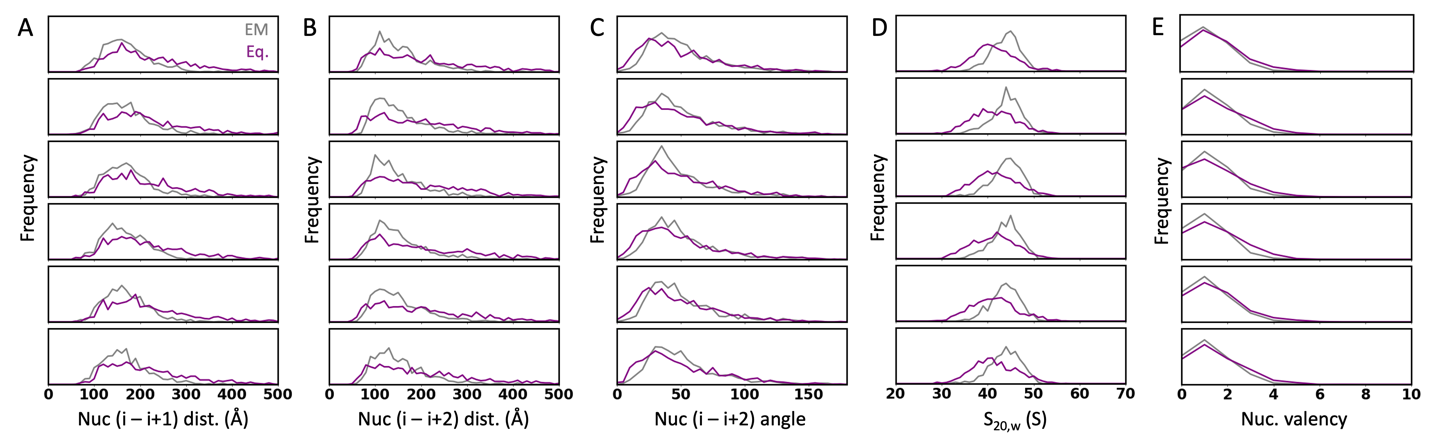

Figure S14: Distribution of inter-nucleosome properties – A) i-i+1 nucleosome distance, B) i-i+2 nucleosome distance, C) angle formed by the COM of nucleosomes i-i+1-i+2, D) sedimentation coefficients calculated for 12 nucleosome segments and E) nucleosome valency estimated using the 11 nm COM-COM distance cutoff – after energy minimization (grey) and equilibration using GENESIS (purple). Each row from the top-to-bottom corresponds to the clusters 1-5, respectively, and the last row represents Cluster 1*. Note: Cluster 1* is based on the mesoscopic reference structure of Cluster 1 but generated using an independent backmapping run.

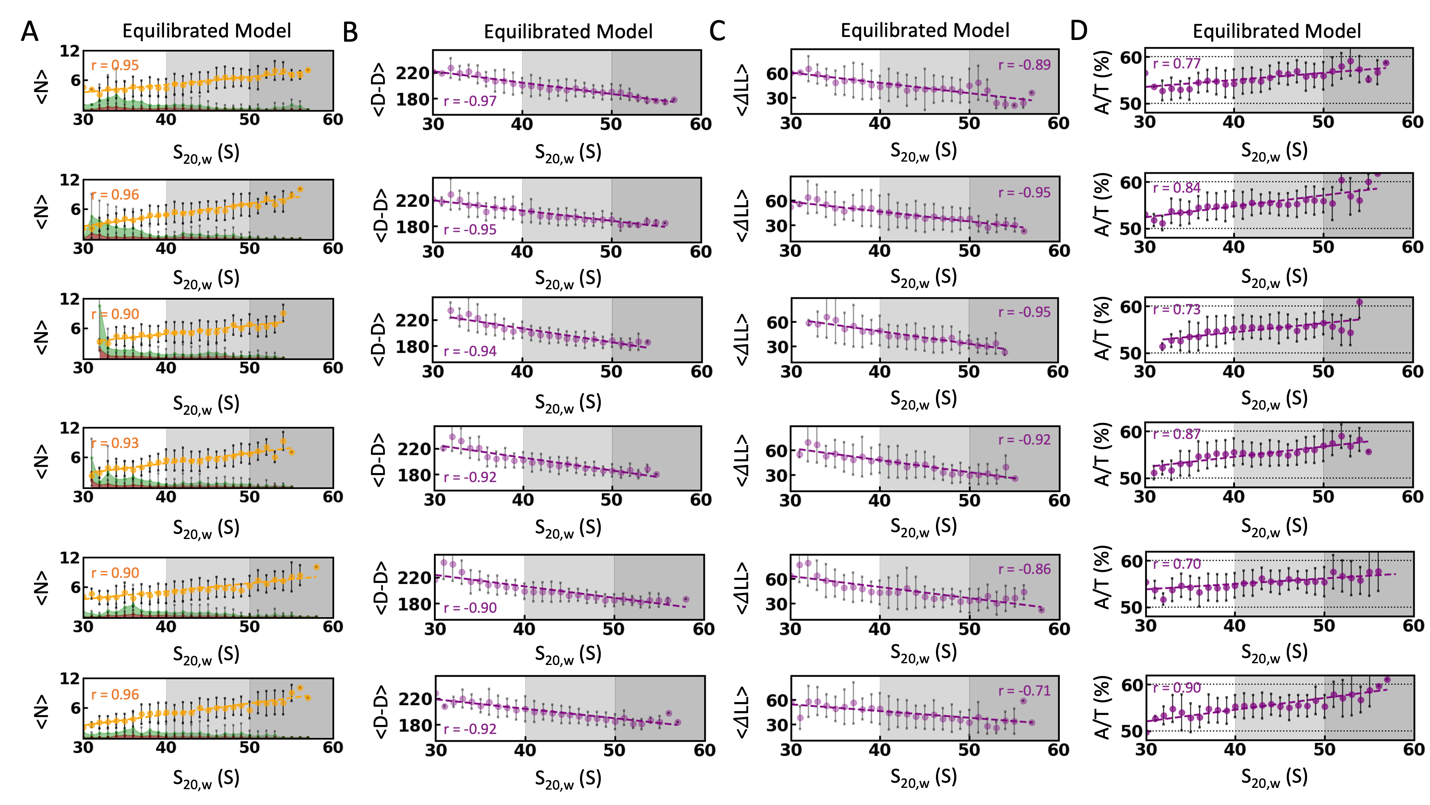

Figure S15: A) The number of linker histones (orange), TFs (green), and BRD4 (red) mapped as the function of the sedimentation coefficient (S_20,w_) of 12-nucleosome chromatin segments for each cluster. Each row from the top-to-bottom corresponds to the clusters 1-5, respectively, and the last row represents Cluster 1*. Note: Cluster 1* is based on the mesoscopic reference structure of Cluster 1 but generated using an independent backmapping run. The error bars are calculated as the standard deviation in each bin. The shaded backgrounds indicate the broadly categorized expanded (white), moderately compact (light grey), and compact (dark grey) chromatin segments. B-D) Same as panel A, but for dyad-dyad distance in bp (D-D; proxy for i-i+1 nucleosome separation), asymmetry in entry/exit linker DNA length (LL; proxy for i-i+2 nucleosome interaction) and A/T fraction (A/T %; proxy for nucleosome positioning signal) of the DNA segments. Pearson’s correlation coefficient (r) is given in the top corner of each panel.

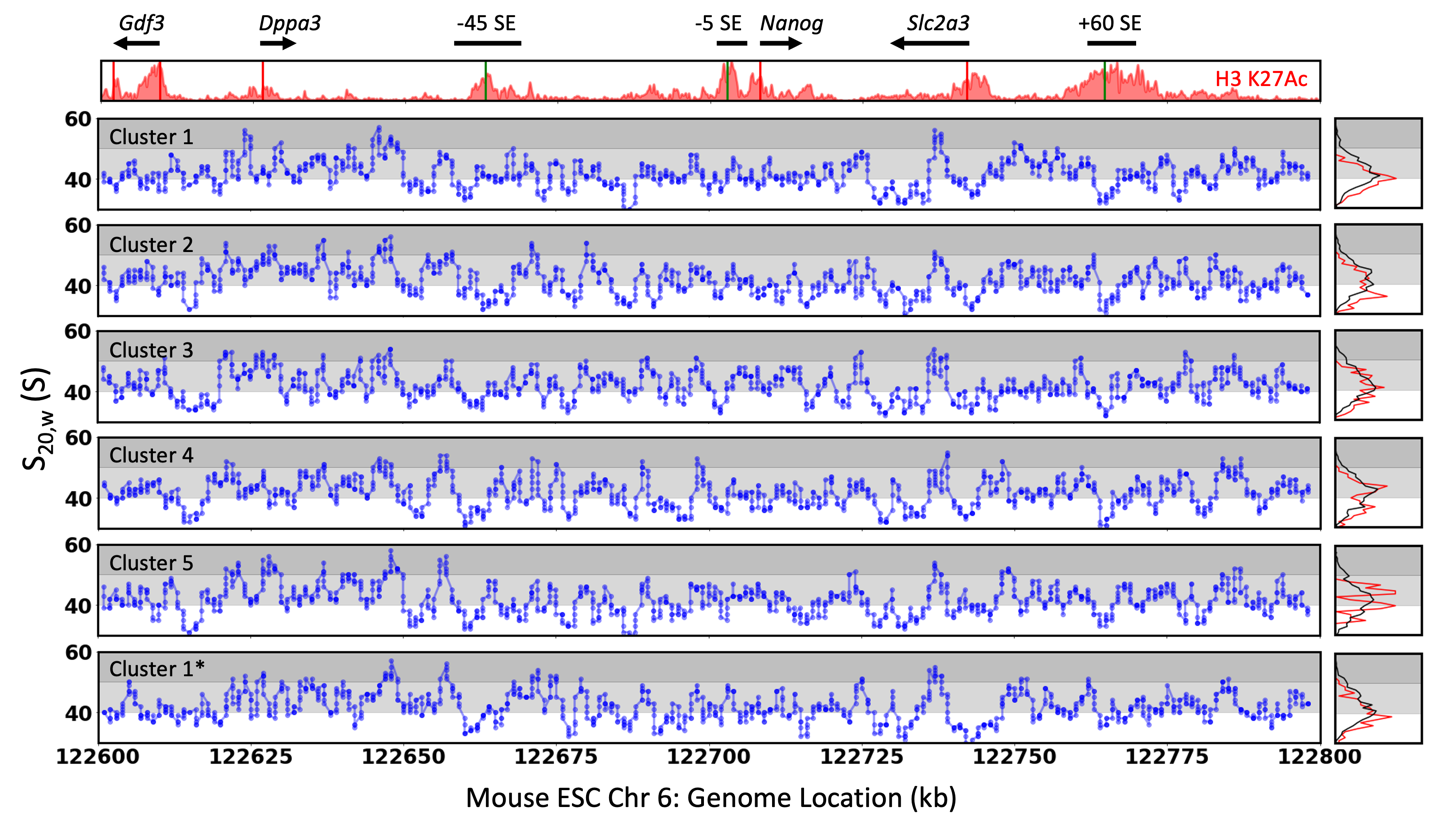

Figure S16: The S_20,w_ of the 12 nucleosome segments as the function of genomic position. The shaded backgrounds indicate the broadly categorized expanded (white), moderately compact (light grey), and compact (dark grey) chromatin segments. The annotations and the H3K27Ac ChIP-seq signals are shown at the top for reference. The distribution of S_20,w_ for H3K27Ac enriched segments (red) and the rest (black) are shown on the right. Note: Cluster 1* is based on the mesoscopic reference structure of Cluster 1 but generated using an independent backmapping run.

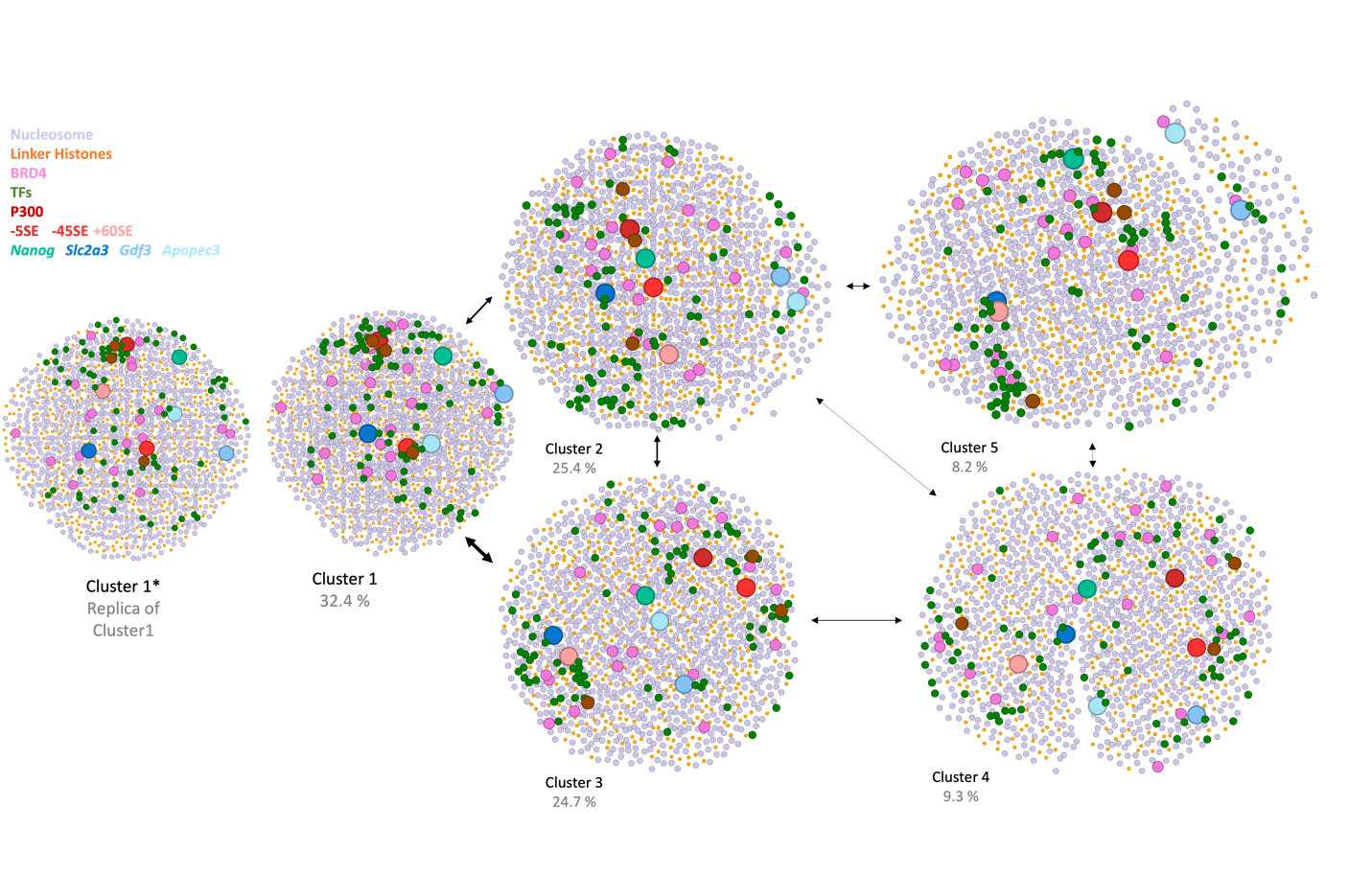

Figure S17: 2D projection of the representative models of the *Nanog* locus using graph theoretical methods. Each circle represents a molecular module (refer to the legend in the top left corner) with size proportional to their radius of gyration, and the proximity is proportional to the minimum distance between the molecules. The arrows indicate the transition between clusters and its width proportional to the transition probabilities calculated from the HiC-metainference simulations.

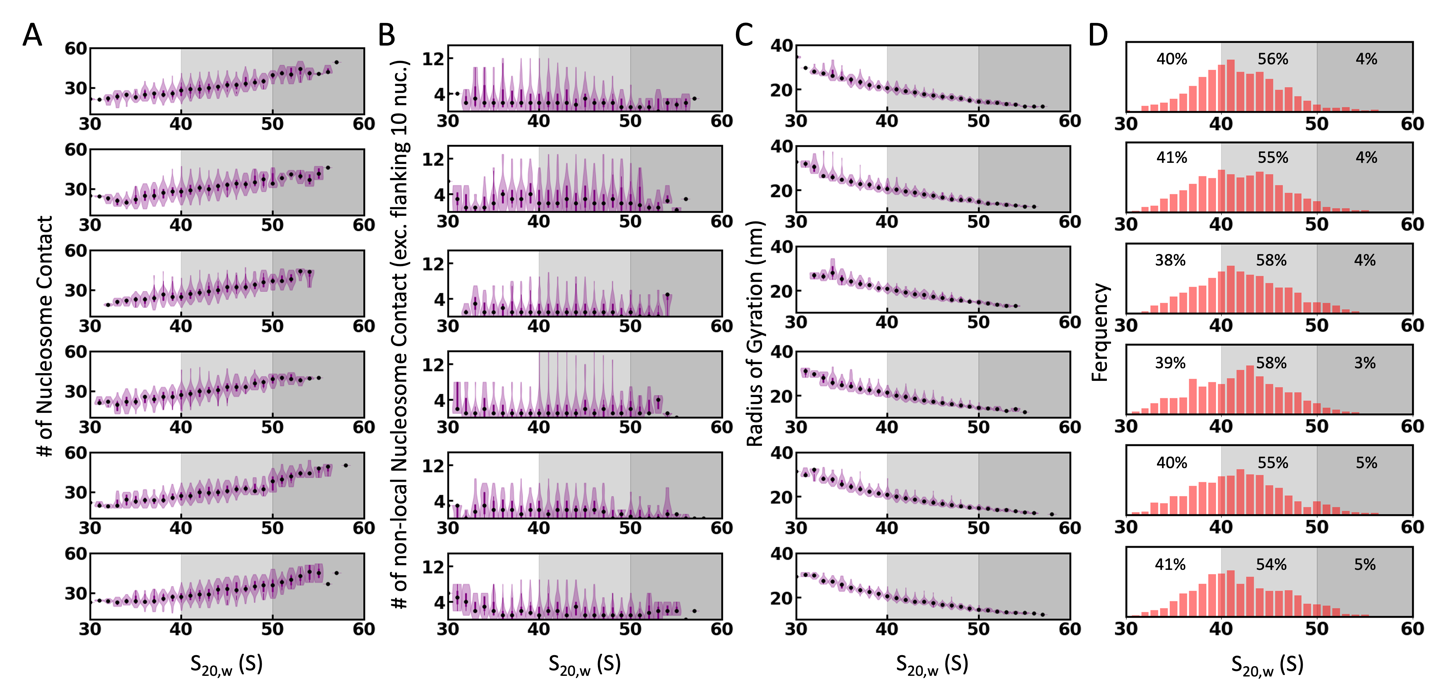

Figure S18: A) The distribution of the number of nucleosome contacts (COM-COM distance < 12 nm) as the function of the sedimentation coefficient (S_20,w_) of 12-nucleosome chromatin segments for each cluster. Each row from the top-to-bottom corresponds to the clusters 1-5, respectively, and the last row represents Cluster 1*. Note: Cluster 1* is based on the mesoscopic reference structure of Cluster 1 but generated using an independent backmapping run. The black circles indicate the median in each bin. The shaded backgrounds indicate the broadly categorized expanded (white), moderately compact (light grey), and compact (dark grey) chromatin segments. B-C) Same as panel A, but for the number of non-local nucleosome interactions and Radius of gyration in each model. D) Distribution of (S_20,w_) of 12-nucleosome chromatin segments for each cluster with the fraction of data from the three categories, identified using differently colored backgrounds, mentioned at the top.

Supplementary Videos 1-5: The videos demonstrate the backmapping of 5kb mesoscopic beads (transparent grey beads) to near-atomistic resolution. The conformations at residue-level resolution generated during the MC sampling are shown in grey, and the selected chromatin conformation based on the MC selection criteria (excluding proteins, for clarity) is shown in the color spectrum of red to white.

**References**

1.	Brandani, G. B., Gu, C., Gopi, S. & Takada, S. Multiscale Bayesian simulations reveal functional chromatin condensation of gene loci. *PNAS Nexus* **3**, (2024).

2.	Bonomi, M., Camilloni, C., Cavalli, A. & Vendruscolo, M. Metainference: A Bayesian inference method for heterogeneous systems. *Sci Adv* **2**, (2016).

3.	Hsieh, T.-H. S. *et al.* Resolving the 3D Landscape of Transcription-Linked Mammalian Chromatin Folding. *Mol Cell* **78**, 539-553.e8 (2020).

4.	Robinson, J. T. *et al.* Juicebox.js Provides a Cloud-Based Visualization System for Hi-C Data. *Cell Syst* **6**, 256-258.e1 (2018).

5.	Tribello, G. A., Bonomi, M., Branduardi, D., Camilloni, C. & Bussi, G. PLUMED 2: New feathers for an old bird. *Comput Phys Commun* **185**, 604–613 (2014).

6.	Plimpton, S. Fast Parallel Algorithms for Short-Range Molecular Dynamics. *J Comput Phys* **117**, 1–19 (1995).

7.	Scherer, M. K. *et al.* PyEMMA 2: A Software Package for Estimation, Validation, and Analysis of Markov Models. *J Chem Theory Comput* **11**, 5525–5542 (2015).

8.	Neph, S. *et al.* BEDOPS: high-performance genomic feature operations. *Bioinformatics* **28**, 1919–1920 (2012).

9.	Williams, L. H. *et al.* Pausing of RNA Polymerase II Regulates Mammalian Developmental Potential through Control of Signaling Networks. *Mol Cell* **58**, 311–322 (2015).

10.	Li, J. *et al.* Single-Molecule Nanoscopy Elucidates RNA Polymerase II Transcription at Single Genes in Live Cells. *Cell* **178**, 491-506.e28 (2019).

11.	Cao, K. *et al.* High-resolution mapping of h1 linker histone variants in embryonic stem cells. *PLoS Genet* **9**, e1003417 (2013).

12.	Zhang, Y., Liu, Z., Medrzycki, M., Cao, K. & Fan, Y. Reduction of Hox Gene Expression by Histone H1 Depletion. *PLoS One* **7**, e38829 (2012).

13.	Chen, J. *et al.* Single-Molecule Dynamics of Enhanceosome Assembly in Embryonic Stem Cells. *Cell* **156**, 1274–1285 (2014).

14.	Xie, L. *et al.* A dynamic interplay of enhancer elements regulates *Klf4* expression in naïve pluripotency. *Genes Dev* **31**, 1795–1808 (2017).

15.	Karr, J. P., Ferrie, J. J., Tjian, R. & Darzacq, X. The transcription factor activity gradient (TAG) model: contemplating a contact-independent mechanism for enhancer–promoter communication. *Genes Dev* **36**, 7–16 (2022).

16.	Castro-Mondragon, J. A. *et al.* JASPAR 2022: the 9th release of the open-access database of transcription factor binding profiles. *Nucleic Acids Res* **50**, D165–D173 (2022).

17.	Chronis, C. *et al.* Cooperative Binding of Transcription Factors Orchestrates Reprogramming. *Cell* **168**, 442-459.e20 (2017).

18.	Lee, B. T. *et al.* The UCSC Genome Browser database: 2022 update. *Nucleic Acids Res* **50**, D1115–D1122 (2022).

19.	Šali, A. & Blundell, T. L. Comparative Protein Modelling by Satisfaction of Spatial Restraints. *J Mol Biol* **234**, 779–815 (1993).

20.	Varadi, M. *et al.* AlphaFold Protein Structure Database: massively expanding the structural coverage of protein-sequence space with high-accuracy models. *Nucleic Acids Res* **50**, D439–D444 (2022).

21.	Jumper, J. *et al.* Highly accurate protein structure prediction with AlphaFold. *Nature* **596**, 583–589 (2021).

22.	Tan, C. & Takada, S. Nucleosome allostery in pioneer transcription factor binding. *Proceedings of the National Academy of Sciences* **117**, 20586–20596 (2020).

23.	Krivov, G. G., Shapovalov, M. V. & Dunbrack, R. L. Improved prediction of protein side-chain conformations with SCWRL4. *Proteins: Structure, Function, and Bioinformatics* **77**, 778–795 (2009).

24.	Wang, X. *et al.* LLPSDB v2.0: an updated database of proteins undergoing liquid–liquid phase separation *in vitro*. *Bioinformatics* **38**, 2010–2014 (2022).

25.	Palacio, M. & Taatjes, D. J. Merging Established Mechanisms with New Insights: Condensates, Hubs, and the Regulation of RNA Polymerase II Transcription. *J Mol Biol* **434**, 167216 (2022).

26.	Abraham, M. J. *et al.* GROMACS: High performance molecular simulations through multi-level parallelism from laptops to supercomputers. *SoftwareX* **1–2**, 19–25 (2015).

27.	Best, R. B. & Hummer, G. Optimized Molecular Dynamics Force Fields Applied to the Helix−Coil Transition of Polypeptides. *J Phys Chem B* **113**, 9004–9015 (2009).

28.	Tan, C. *et al.* Implementation of residue-level coarse-grained models in GENESIS for large-scale molecular dynamics simulations. *PLoS Comput Biol* **18**, e1009578 (2022).

29.	Tesei, G., Schulze, T. K., Crehuet, R. & Lindorff-Larsen, K. Accurate model of liquid–liquid phase behavior of intrinsically disordered proteins from optimization of single-chain properties. *Proceedings of the National Academy of Sciences* **118**, (2021).

30.	Zheng, G., Lu, X.-J. & Olson, W. K. Web 3DNA--a web server for the analysis, reconstruction, and visualization of three-dimensional nucleic-acid structures. *Nucleic Acids Res* **37**, W240–W246 (2009).

31.	Farr, S. E., Woods, E. J., Joseph, J. A., Garaizar, A. & Collepardo-Guevara, R. Nucleosome plasticity is a critical element of chromatin liquid–liquid phase separation and multivalent nucleosome interactions. *Nat Commun* **12**, 2883 (2021).

32.	Li, G. & Widom, J. Nucleosomes facilitate their own invasion. *Nat Struct Mol Biol* **11**, 763–769 (2004).

33.	Niina, T., Brandani, G. B., Tan, C. & Takada, S. Sequence-dependent nucleosome sliding in rotation-coupled and uncoupled modes revealed by molecular simulations. *PLoS Comput Biol* **13**, e1005880 (2017).

34.	Dabrowski-Tumanski, P., Rubach, P., Niemyska, W., Gren, B. A. & Sulkowska, J. I. Topoly: Python package to analyze topology of polymers. *Brief Bioinform* **22**, (2021).

35.	Tubiana, L., Polles, G., Orlandini, E. & Micheletti, C. KymoKnot: A web server and software package to identify and locate knots in trajectories of linear or circular polymers. *The European Physical Journal E* **41**, 72 (2018).

36.	Jung, J. *et al.* GENESIS: a hybrid-parallel and multi-scale molecular dynamics simulator with enhanced sampling algorithms for biomolecular and cellular simulations. *Wiley Interdiscip Rev Comput Mol Sci* **5**, 310–323 (2015).

37.	Jung, J., Tan, C. & Sugita, Y. GENESIS CGDYN: large-scale coarse-grained MD simulation with dynamic load balancing for heterogeneous biomolecular systems. *Nat Commun* **15**, 3370 (2024).

38.	Terakawa, T. & Takada, S. Multiscale Ensemble Modeling of Intrinsically Disordered Proteins: p53 N-Terminal Domain. *Biophys J* **101**, 1450–1458 (2011).

39.	Tan, C. & Takada, S. Dynamic and Structural Modeling of the Specificity in Protein–DNA Interactions Guided by Binding Assay and Structure Data. *J Chem Theory Comput* **14**, 3877–3889 (2018).

40.	Voong, L. N. *et al.* Insights into Nucleosome Organization in Mouse Embryonic Stem Cells through Chemical Mapping. *Cell* **167**, 1555-1570.e15 (2016).

41.	Kagey, M. H. *et al.* Mediator and cohesin connect gene expression and chromatin architecture. *Nature* **467**, 430–5 (2010).

42.	Norrie, J. L. *et al.* Nucleome Dynamics during Retinal Development. *Neuron* **104**, 512-528.e11 (2019).

43.	Shen, Y. *et al.* A map of the cis-regulatory sequences in the mouse genome. *Nature* **488**, 116–20 (2012).
